# Synchronization between chloroplastic and cytosolic protein synthesis for photosynthesis complex assembly

**DOI:** 10.1101/2024.05.03.592458

**Authors:** Taisei Wakigawa, Tomoya Fujita, Naohiro Kawamoto, Yukio Kurihara, Yuu Hirose, Takashi Hirayama, Hirotaka Toh, Tomoko Kuriyama, Atsushi Hashimoto, Eriko Matsuura-Suzuki, Keiichi Mochida, Minoru Yoshida, Minami Matsui, Shintaro Iwasaki

**Author notes:** Correspondence (S.I.). These authors contributed equally (Taisei Wakigawa, Tomoya Fujita, Naohiro Kawamoto).

## Abstract

Through symbiosis, subunits of chloroplastic complexes are encoded in distinct genomes in the nucleus and organelles. For plant cells to maintain the stoichiometry of subunits and respond to environmental cues, the orchestration of the nuclear and organellar gene expression systems is an essential task. However, the mechanism maintaining chloroplastic complexes remains largely enigmatic. Here, we simultaneously assessed the translatomes of the chloroplast and the cytoplasm via ribosome profiling and revealed the differential mechanisms employed by these two systems to cope with acute light/dark transitions: in chloroplasts, translational regulation is employed, whereas in the cytoplasm, control of the mRNA abundance is implemented. This strategy is widely conserved in land plants (*Arabidopsis* and the grass plant *Brachypodium*) and green algae (*Chlamydomonas*). The translational control in chloroplasts may be established based on organelle symbiosis; the primitive chloroplast in Glaucophyta (*Cyanophora*) was found to have already acquired translational control, whereas cyanobacteria (*Synechocystis*) control the mRNA abundance. Moreover, reduced plastoquinones and active cytosolic protein synthesis drive chloroplastic translation of the complex subunits in the light. Our work reveals an early origin of coordination of chloroplast and nuclear/cytoplasmic gene expression upon light exposure.

## Introduction

Most biological energy on Earth is supplied through photosynthesis. This reaction occurs in chloroplasts through the multiple protein complexes in plants ^1–5^. The photosynthetic apparatus is assembled from proteins encoded in two distinct genomes: the nuclear genome and the chloroplast genome ^6,7^. This complicated scheme is due to ancient cyanobacterial symbiosis and coevolution; some genes encoding subunits of the photosynthetic apparatus were transferred to the nuclear genome, whereas others are still encoded in the chloroplast genome. Ultimately, the subunits of the photosynthetic apparatus are translated in two distinct subcellular locations—the cytoplasm and the chloroplast stroma. Considering the subunits of photosystem I (PSI), photosystem II (PSII), cytochrome *b6f* complex (Cyt *b6f*), NADH dehydrogenase-like complex (NDH), and FoF1 ATP synthase in *Arabidopsis thaliana*, 47 subunits should be synthesized in the cytosol and 44 in the chloroplast stroma (Table S1). A balanced supply of subunits from these two origins is crucial to proteostasis in plant cells, since excess free subunits or immature complexes can result in detrimental effects on cells ^8,9^. However, the mechanisms underlying the coordinated protein synthesis between the cytosol and chloroplast stroma remain elusive.

Oxidative phosphorylation (OXPHOS) complexes in mitochondria are faced with a similar challenge, also requiring the coordination of gene expression between the nuclear genome and the mitochondrial genome ^10–12^. The two distinct gene expression systems in the cytosol and the mitochondrial matrix are synchronized at the translational level to allow adaptation to respiratory metabolism ^13,14^. Especially in yeast, cytosolic translation status dictates mitochondrial translation ^13^. Although it is logical to hypothesize the presence of similar synchronization of gene expression between the cytosol and chloroplasts in plants to support their response to environmental cues, the presence of such a mechanism is unclear.

To investigate the cooperation of gene expression systems for the nuclear genome and chloroplast genome, we harnessed ribosome profiling, a technique based on deep sequencing of ribosome-protected RNA fragments (*i.e.*, ribosome footprints) generated by RNase treatment ^15,16^. We employed this strategy, which has been used for the simultaneous assessment of cytosolic and mitochondrial translation in mammals ^17–19^, to monitor cytosolic and chloroplastic protein synthesis in land plants, green algae, and Glaucophyta. We found that upon environmental transitions between light and dark conditions, the supply of subunits for photosynthetic apparatuses aligned with the demands, *i.e.*, a lower synthesis rate under dark conditions and a higher synthesis rate under light conditions. However, the mechanistic basis of this orchestration differed between the subcellular locations: regulation of the RNA abundance in the cytosol and regulation of translation in chloroplasts. On the other hand, in cyanobacteria, the evolutionary origin of chloroplasts, the RNA abundance is regulated, suggesting the emergence of a translational regulatory mechanism upon symbiosis. We also found that a reduced form of plastoquinone (PQ) produced by PSII activates translation in chloroplasts. Moreover, the cytoplasmic translation process drives chloroplastic translation, but not vice versa. Our study provides fundamental insight into the synchronization of gene expression from two different genomes for environmental adaptation.

## Results

### Plant ribosome profiling simultaneously assesses translation in the cytosol and chloroplast stroma

Given that the subunits of the photosynthesis complexes in the thylakoid membrane are encoded by dual genomes, we inferred the presence of coordination between the two gene expression systems. To monitor protein synthesis in both the cytosol and chloroplasts simultaneously (Figure 1A), we obtained whole-cell lysates containing both cytosolic and chloroplastic ribosomes (Figure 1A) (see the Materials and Methods section for details). For concurrent assessment of two translation systems, we explored the library preparation conditions, titrating micrococcal nuclease (MNase) and RNase I, well-used RNases for ribosome profiling, for *Arabidopsis thaliana* seedlings and green algae *Chlamydomonas reinhardtii* (Figure S1A-D and Table S2). In both species, high RNase I conditions generated a representative size of reads originating from nuclear genome-encoded mRNAs (*i.e.*, cytoribosome footprints) and chloroplastic mRNA fragments (*i.e.*, chlororibosome footprints) (Figure S1A-B). Since three-nucleotide periodicity of reads was a hallmark of ribosome footprints, we evaluated the quality by “periodicity score” (see Methods for details) ^20^. Basically, RNase I conditions provided high 3-nt periodicity in a dose-dependent manner (Figure S1C-D). Importantly, both cytoribosome footprints and chlororibosome footprints exhibited a similar trend in data quality across the tested conditions; better conditions for cytoribosome footprints were also better for chlororibosome footprints. Thus, we used high-RNase I conditions in the downstream analysis to assess footprints from cytoribosomes and chlororibosomes.

**Figure 1.**
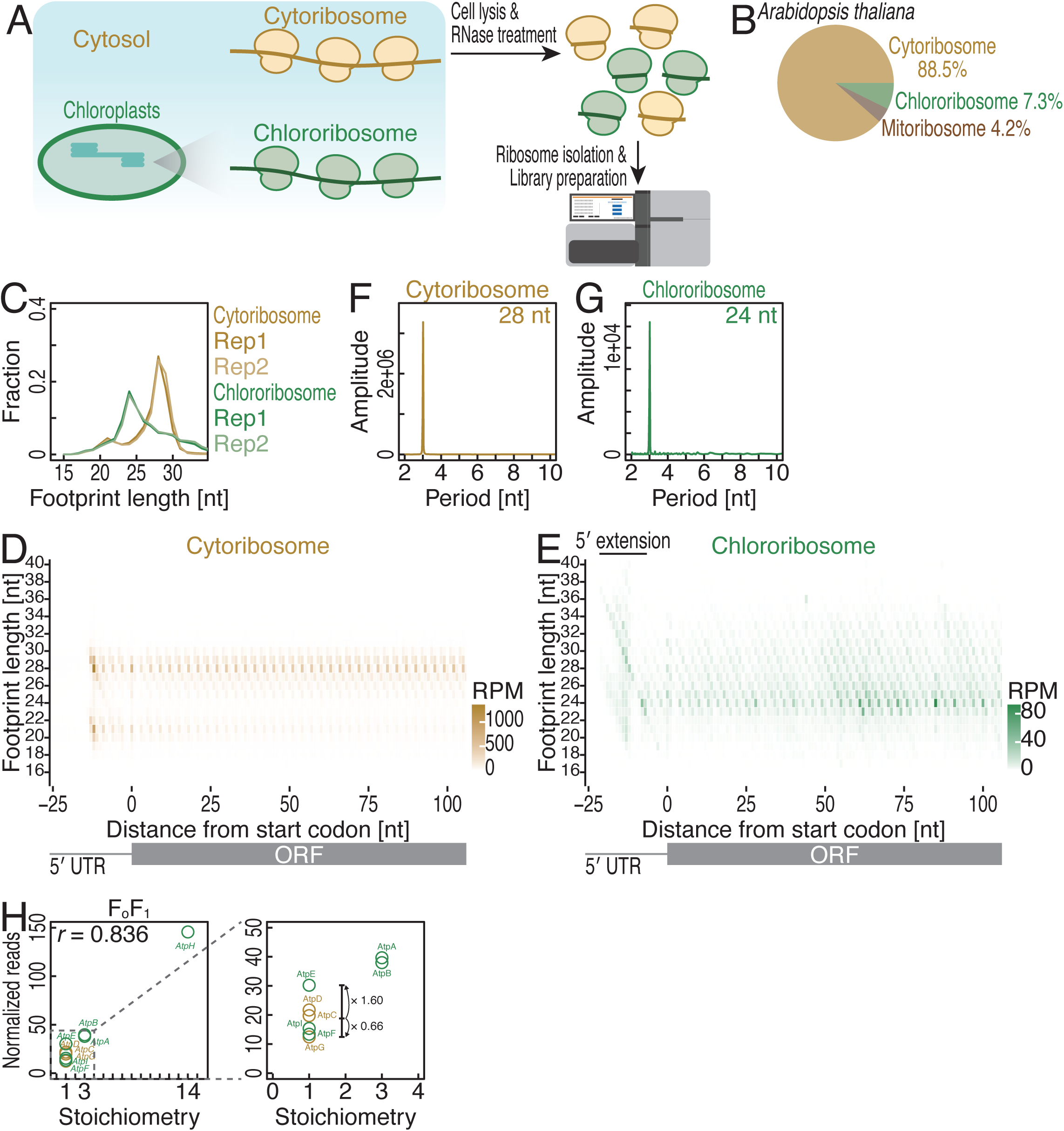
Simultaneous assessment of cytosolic and chloroplastic translations by ribosome profiling. (A) Schematic representation of the procedure for plant ribosome profiling, which recovers footprints stemming from cytoribosomes and chlororibosomes. (B) Pie chart showing the proportions of footprints from cytoribosomes, mitoribosomes, and chlororibosomes in *A. thaliana*. (C) Length distribution of cytoribosome footprints and chlororibosome footprints. (D and E) Metagene plots of the 5′ ends of cytoribosome footprints (D) and chlororibosome footprints (E) along the length around start codons (the first nucleotide of the start codon was set to 0). The color scale indicates the read abundance. RPM, reads per million mapped reads. (F and G) Discrete Fourier transform of cytoribosome footprints (28 nt) (F) and chlororibosome footprints (24 nt) (G) downstream of the start codon (from position −9 to position 300). (H) Correlation between the normalized read density (reads/codon) and the subunit stoichiometry for the chloroplast FoF1 ATP synthase, with the zoomed-in inset highlighting data for one component factor. See also Figures S1-2 and Tables S1-3.

Here, we started surveying cytoplasmic and chloroplastic translations focusing on *Arabidopsis thaliana* seedlings. The majority (88.5%) of the reads originated from cytoribosome footprints, while chlororibosome footprints constituted a small but significant fraction (7.3%) of the reads (Figure 1B). The lengths of ribosome footprints are determined by the sizes and conformational status of ribosomes and thus often fall into characteristic size ranges ^21–27^. As observed in other eukaryotes ^21,25,27^, cytosolic ribosomes showed two distinct peak sizes—short, ∼21 nucleotides (nt); and long, ∼28 nt (Figures 1C and S2A). These sizes represent the absence and presence of an A-site tRNA on the ribosome, respectively ^25^. On the other hand, the chloroplast ribosome footprint lengths were restricted to ∼24 nt (Figures 1C and S2A), consistent with a recent study ^28^. Notably, although earlier studies showed a wider distribution of longer footprints from chloroplasts ^29–35^, this observation may have been due to insufficient RNase digestion or conformational dynamics of chlororibosomes (see the “Limitations of the study” section). The presence of three-nucleotide periodicity in footprints ensures codon resolution in the data. Indeed, the metagene plots showed a periodic pattern of reads along the open reading frame (ORF) for both cytoribosome footprints and chlororibosome footprints (Figures 1D-E and S2B). This observation was further supported by discrete Fourier transform calculations (Figure 1F and 1G). These data indicated that our libraries were qualified for the simultaneous assessment of cytosolic and chloroplastic translation. We also found that our ribosome profiling data were highly reproducible in both ORF-wise (Figure S2C) and codon-wise (Figure S2D) assessments.

In our metagene analysis of chloroplast ribosome footprints, we observed the 5′ extension of the footprints around the start codon (Figure 1E), reminiscent of bacterial ribosome footprints (Figure S1E). The increased lengths of bacterial ribosome footprints around the start codon can be explained by the failure of RNA cleavage due to the RNA duplexes formed between the Shine‒Dalgarno (SD) sequence in an mRNA and the anti-SD sequence in a 16S rRNA ^22,26^. Although SD/anti-SD base pairing facilitates translation initiation in bacteria ^36,37^, the importance for chloroplast translation has remained controversial ^38–45^. Our data suggested that the region upstream of the AUG codon, including the SD sequence, may interact with an anti-SD sequence in the chloroplast 16S rRNA.

### Chloroplastic ATP synthase subunits are synthesized in proportion to the stoichiometry of the complex

Multisubunit complexes may contain each protein subunit in different stoichiometries. Imbalanced subunit synthesis leads to nonfunctional complexes or orphan proteins, which may have adverse effects and thus should be degraded ^9^. To reduce the cellular cost, cells synthesize protein subunits in proportion to the stoichiometry of the complex ^46–48^. The bacterial FoF1 ATP synthase, which consists of subunits at a stoichiometric ratio of 1:2:3:10, is a notable example ^46^. The synthesis of the homologous complex on the chloroplast thylakoid membrane is more challenging; the subunit proteins are encoded by genes that are not in the same operon, as they are in bacteria; are encoded in two genomes; and are translated in different subcellular domains.

Even with these difficulties, the subunits of the chloroplastic FoF1 ATP synthase are still proportionally synthesized, irrespective of the translation system (Figure 1H). The variation in subunit synthesis, particularly for one component factor, remained within a 2-fold range (Figure 1H), consistent with observations in bacteria, yeasts, mice, humans, and zebrafish ^46–48^. A similar pattern of proportional subunit synthesis was suggested for maize, *Arabidopsis*, and *Chlamydomonas* via ribosome profiling ^30,49,50^. However, these studies did not consider the subunits translated by cytoribosomes. Here, we showed that dual translation systems coordinately control the supply of subunits.

### Light-dependent synchronization of dual gene expression systems for chloroplastic multisubunit complexes

Since FoF1 ATP synthase is a component of the photosynthesis system, we investigated the dynamics of its synthesis rate during acute light-to-dark and dark-to-light transitions (Figures 2A, S2A-E, and Table S3). With reduced light exposure, we observed decreases in the footprints of both translation systems for FoF1 ATP synthase subunits (Figure 2B, left). Reillumination of the plant restored subunit synthesis (Figure 2B, left). Thus, the synchronization of the two gene expression systems adjusted to meet the demand for FoF1 ATP synthase in response to environmental changes.

**Figure 2.**
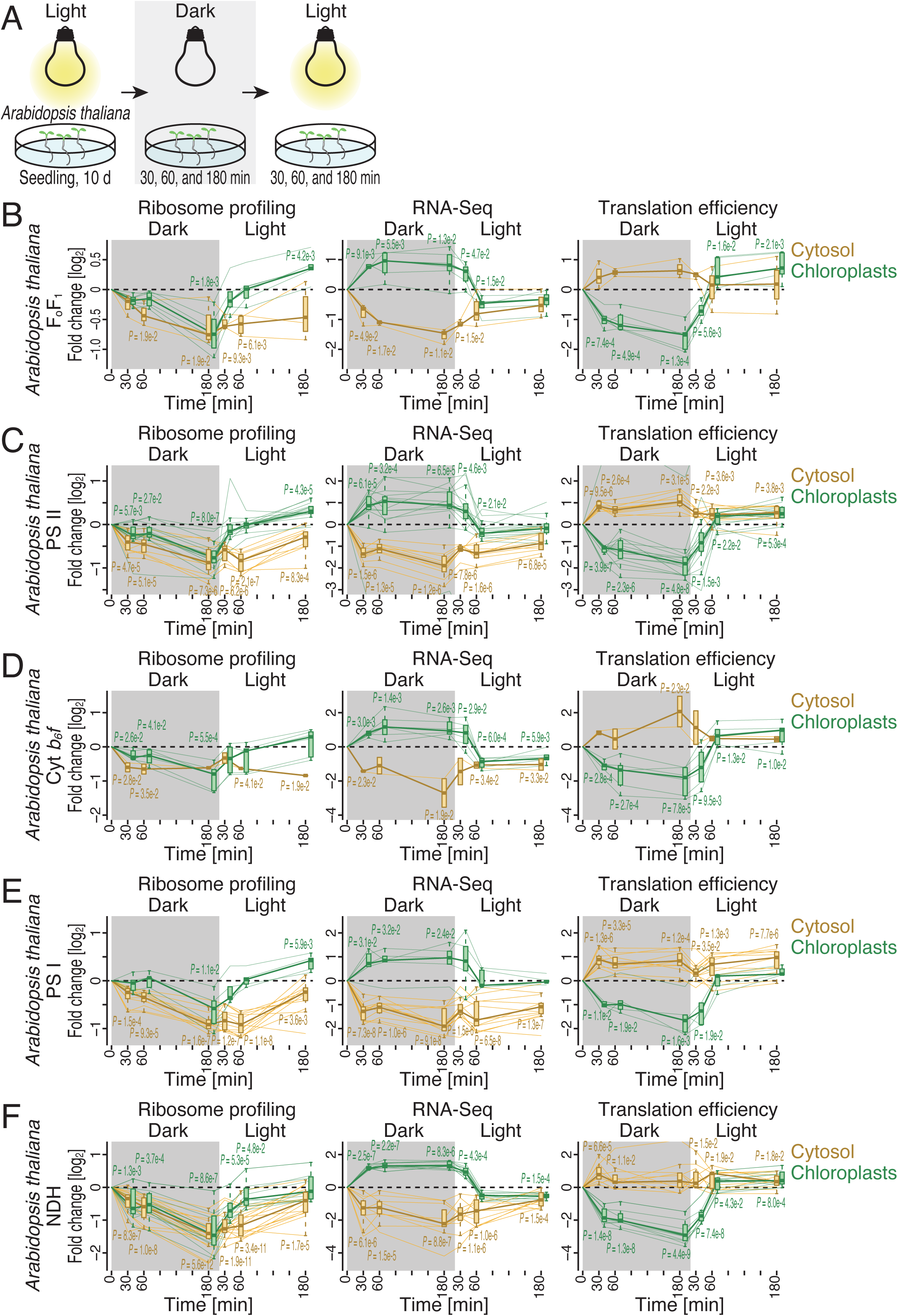
Light/dark transitions coordinate the protein supply via dual gene expression systems. (A) Schematic representation of the experimental design used to monitor the light- dependent synthesis of the subunits of thylakoid membrane-embedded complexes. (B-F) Changes in ribosome profiling, RNA-Seq, and translation efficiency during the light-to-dark and dark-to-light transitions for the subunits of FoF1 ATP synthase (B), PSII (C), Cyt *b6f* (D), PSI (E), and NDH (F) in *A. thaliana*. The box plots (B-F) show the median (center line), upper/lower quartiles (box limits), and 1.5× interquartile range (whiskers). The black line indicates the median at each time point. Significance was determined by the Mann‒Whitney *U* test. See also Figure S2 and Tables S1-3.

However, strikingly, the mechanistic basis differed between the two gene expression systems. Simultaneous RNA-Seq analysis (Figure S2C, S2E, and Table S3) revealed drastic reductions in the levels of cytoplasmic mRNAs encoding FoF1 ATP synthase subunits under dark conditions and the restoration of their levels upon reillumination (Figure 2B, middle). Thus, the alterations in cytoribosome footprints (Figure 2B, left) originated predominantly from transcript availability. Consequently, the change in the “translation efficiency”, a relative score indicating the over- or underrepresentation of ribosome footprints relative to RNA-Seq fragments (Figure S2E and Table S3), did not follow the changes in the footprint (Figure 2B, right). In contrast, the changes in chloroplastic mRNA levels could not explain the footprint dynamics upon light exposure (especially under dark conditions, when chloroplastic mRNAs were enriched, unlike footprint changes) (Figure 2B, middle). Rather, the translation efficiency was a major regulatory mechanism (Figure 2B, right).

In addition to FoF1 ATP synthase, the other photosynthesis complexes, including PSII, Cyt *b6f*, PSI, and NDH, are multisubunit assemblies that require gene expression via dual systems. Thus, we further extended our analysis to those complexes. Indeed, the subunits of all these complexes showed essentially the same dynamics as FoF1 ATP synthase; the dark and light transitions induced synchronized subunit synthesis from the two gene expression systems, as evidenced by the ribosome profiling results (Figures 2C- F and S2F). RNA-centric regulation was dominant in the cytosol, whereas translation-centric regulation was mainly observed in chloroplasts.

We investigated whether chloroplastic RNA processing events occurring before translation may explain the dynamics of ribosome loading on chloroplastic mRNAs during light-dark transitions. Given that a subset of chloroplastic mRNAs possess group I and II introns ^7,51,52^, we explored whether splicing is affected by light-dark transition via RNA-Seq data. Across 15 introns in mRNAs ^52^, 6 introns are found in the mRNAs encoding photosynthesis complexes. We did not observe the dynamic changes in the intron retention profiles in our conditions (Figure S2G).

We also analyzed RNA editing rates. C-to-U RNA editing is omnipresent in land plants’ chloroplasts ^7,53,54^. Our RNA-Seq also detected the C-to-U conversion in many of the known editing sites (28/37) ^55^. Twenty of 28 sites were found in the genes encoding for subunits of photosynthesis complexes. Again, we could not detect drastic editing rate changes that explain the translation changes (Figure S2H). We note that chloroplasts in green algae generally lack RNA editing ^56^, while drastic translation changes upon light-dark transition were conserved in *Chlamydomonas* (see below).

Notably, these regulations were found in a subset of mRNA and thus could not explain translation regulations on wider chloroplastic mRNAs we observed, if such regulation indeed exists.

### Conservation of synchronization between dual gene expression systems

These observations led us to expand our analysis to include green lineages, such as *Brachypodium distachyon* (Figure S3A-I and Table 4), a model grass plant ^57^, and green algae *Chlamydomonas reinhardtii* (Figure S4A-I and Table S5). As was possible for *A. thaliana*, ribosome profiling of these species allowed us to monitor both cytosolic and chloroplastic protein synthesis, as evidenced by the 3-nt read periodicities (Figures S3C- E and S4C-E).

Both green lineages also exhibited environment-tailored protein synthesis for thylakoid membrane-embedded complexes, which was downregulated in the dark and restored in the light (Figure 3B-C), as observed in *Arabidopsis* (Figure 3A). Moreover, the modes of regulation were also conserved; protein subunits generated in the cytosol were regulated at the RNA level, whereas chloroplastic subunits were regulated at the translational level (Figures 3A-C, S3J-O, and S4J-N).

**Figure 3.**
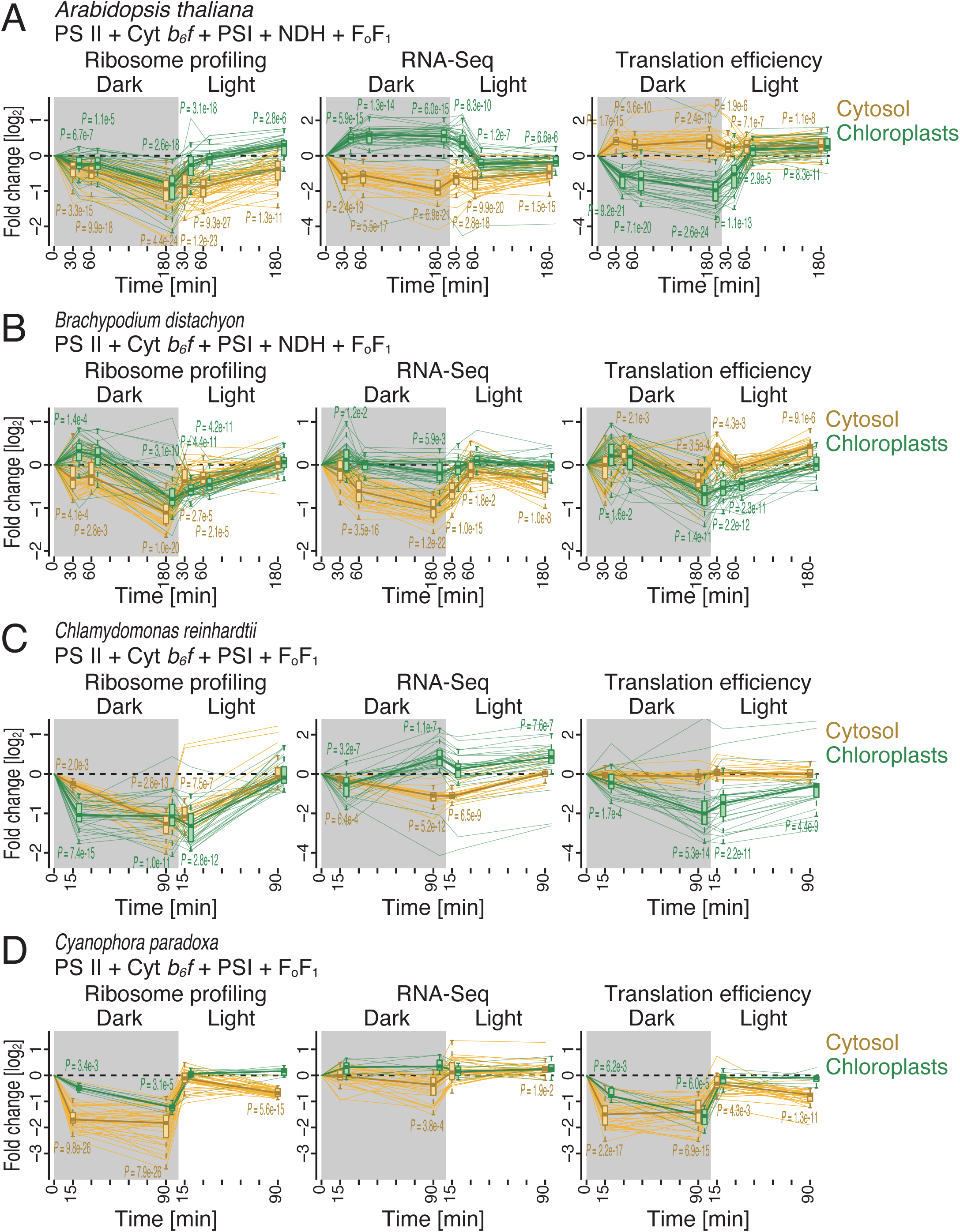
Conservation of the regulation of cytosolic mRNA abundances and chloroplastic translation. (A-D) Changes in ribosome profiling, RNA-Seq, and translation efficiency during the light-to-dark and dark-to-light transitions for the subunits of FoF1 ATP synthase, PSII, Cyt *b6f*, PSI, and NDH in *A. thaliana* (A), *B. distachyon* (B), *C. reinhardtii* (C), and *C. paradoxa* (D). *C. reinhardtii* and *C. paradoxa* do not possess NDH. The box plots (A-D) show the median (center line), upper/lower quartiles (box limits), and 1.5× interquartile range (whiskers). The black line indicates the median at each time point. Significance was determined by the Mann‒Whitney *U* test. See also Figures S3-5 and Tables S1-2 and S4-6.

Ribosome footprint analysis in *B. distachyon* may require additional caution. The nuclear genome of *B. distachyon* contains a large number of chloroplast gene duplications due to gene transfer (nuclear plastid genes) ^58^. Thus, the reads mapped to the chloroplast genome could also be assigned to nuclear plastid genes because of their high sequence similarity (Figure S3P-Q). However, we assumed that the reads mapped to chloroplast transcripts predominantly represented chloroplastic translation because 1) the majority of the nuclear plastid genes were known to be pseudogenes ^58^ and 2) the footprints mapped to the chloroplast genome had different sizes than those mapped to the nuclear genome (Figure S3B).

The conservation of the light-dark response mechanism for subunit synthesis led us to investigate primitive plant cells. Here, we used Glaucophyta (*Cyanophora paradoxa*), which is considered the most ancient lineage in the kingdom Plantae ^59,60^. Chloroplasts of *C. paradoxa* resemble their cyanobacterial origin with the peptidoglycan “cell wall” between the membranes ^61^. Ribosome profiling of this species revealed the footprints of cytoribosomes and chlororibosomes (Figure S5A-I and Table S6). We found that primordial chloroplasts already possessed a translational regulatory mechanism activated in response to light (Figures 3D and S5J-N). Thus, these data suggested the early implementation of translational control of chloroplastic gene expression during cyanobacterial symbiosis. Notably, in contrast to the RNA level control observed in the green lineages, cytoplasmic synthesis of subunits was regulated at the translational level in Glaucophyta (Figures 3D and S5J-N), implying the divergence of regulatory strategies during evolution.

### Switching of the regulatory mechanism upon endosymbiotic gene transfer

During coevolution, chloroplast-to-nuclear gene transfer (endosymbiotic gene transfer) shapes plant genomes. Although this gene transfer was largely completed at the early stage of symbiosis ^62^, independent events occurred in the green lineages. For example, PsaI in the PSI complex and PetN in Cyt *b6f* were still encoded in the chloroplast genome in *Arabidopsis* but were transferred into the nuclear genome in *Chlamydomonas*. The mechanisms controlling light-dark transition-associated expression are closely aligned with the gene locus (Figure 4A-B); specifically, the expression of chloroplast genome-encoded *psaI* and *petN* was regulated at the translational level in *Arabidopsis*, whereas the expression of these genes, encoded in the nuclear genome in *Chlamydomonas*, was altered at the RNA level in this species.

**Figure 4.**
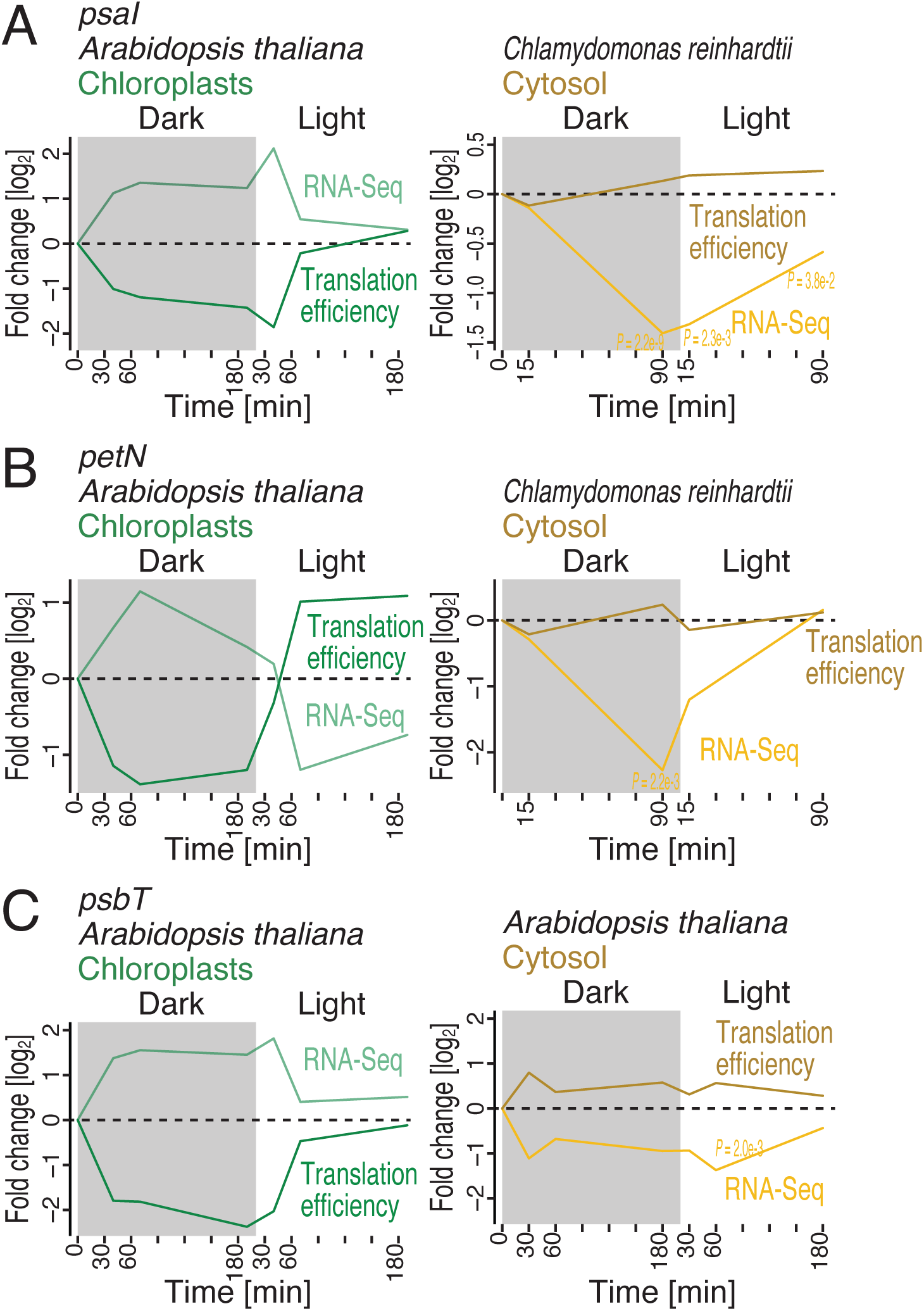
Switch in the regulatory mechanism associated with endosymbiotic gene transfer. (A-B) Changes in ribosome profiling, RNA-Seq, and translation efficiency during the light-to-dark and dark-to-light transitions for *psaI* (A, a subunit of the PSI complex) and *petN* (B, a subunit of Cyt *b6f*) in *A. thaliana* and *C. reinhardtii*. (C) Changes in ribosome profiling, RNA-Seq, and translation efficiency during the light-to-dark and dark-to-light transitions for paralogs of *psbT* (a subunit of PSII) in *A. thaliana*. In A-C, significance was determined by the Mann‒Whitney *U* test. See also Tables S1-2.

Moreover, the same gene transfer-coupled regulatory switch was found in paralogs within the same species. *Arabidopsis* possesses two paralogs of the PSII subunit PsbT, one encoded in the nuclear genome and the other in the chloroplast genome. Again, the responses of these *psbT* paralogs to light/dark environments were driven by the general strategy predominant in each subcellular location: the paralog synthesized in the chloroplast stroma was under translational control, and the paralog synthesized in the cytoplasm was under RNA control (Figure 4C).

These data support the idea that the switch in the regulatory mechanism is associated with gene transfer from the chloroplast genome to the nuclear genome.

### RNA level control is an ancestral mechanism in cyanobacteria

Considering the above findings, we sought to determine how cyanobacteria, the origin of chloroplasts, cope with changes in light conditions. Ribosome profiling and RNA-Seq analysis in *Synechocystis* sp. PCC6803 (Figure S6A-H and Table S7) revealed that the FoF1 and NDH complexes (among others) exhibited on-demand dynamics in response to light cues (Figures 5A-B and S6I-L). However, this regulation relied mainly on changes in RNA abundances rather than on translational control (Figure 5A-B). Thus, these data suggested the conversion of the regulatory mechanism during chloroplast symbiosis from RNA-centric regulation in cyanobacteria to translation-centric regulation in chloroplasts (Figures 5C and Figure S6M).

**Figure 5.**
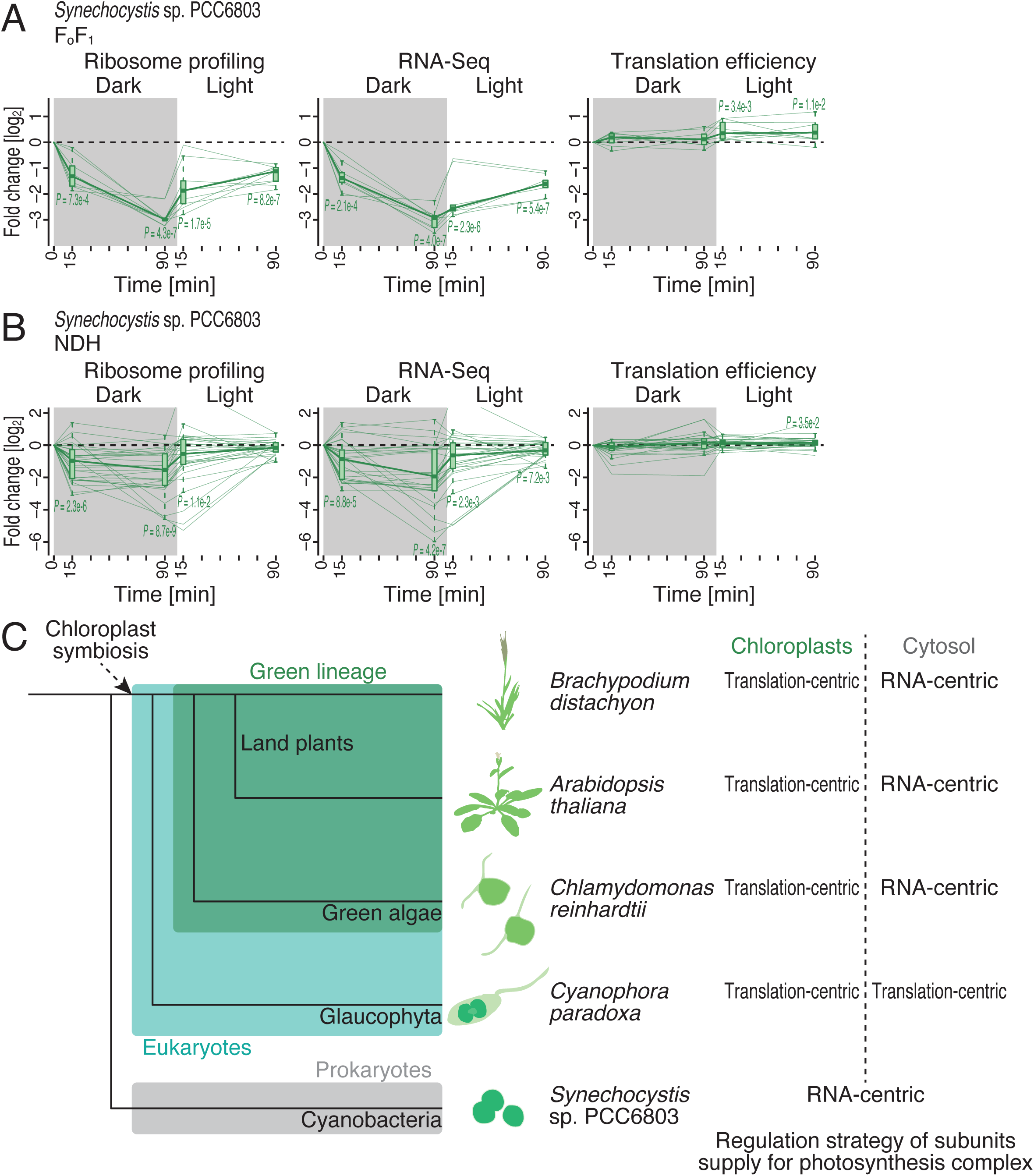
Absence of light-dependent translational regulation in cyanobacteria. (A-B) Changes in ribosome profiling, RNA-Seq, and translation efficiency during the light-to-dark and dark-to-light transitions for the subunits of FoF1 ATP synthase (A) and NDH (B) in *Synechocystis* sp. PCC6803. (C) Model of light-dependent regulation of the assembly of the photosynthetic apparatus among species. The box plots (A-B) show the median (center line), upper/lower quartiles (box limits), and 1.5× interquartile range (whiskers). The black line indicates the median at each time point. Significance was determined by the Mann‒Whitney *U* test. See also Figure S6 and Tables S1-2 and S7.

### Reduced PQ increases translation in chloroplasts

Given the finding of light-dependent translational regulation in chloroplasts, we reasoned that photosynthesis may generate a signal to facilitate protein synthesis. To test this hypothesis, we treated *Arabidopsis* seedlings with 3-(3,4-dichlophenyl)1,1-dimethylurea (DCMU), a PSII inhibitor that blocks electron flux from PSII to PQ ^63,64^ and decreases the abundance of plastoquinol (PQH2), the reduced form of PQ (Figure 6A), in the dark and then exposed them to light (Figure 6B). The dark-to-light transition increased the translation efficiency of the chloroplastic mRNAs encoding complex subunits (Figures 6C, S7A-C, and Table S8), as observed in the previous experiments (Figures 2 and 3A). However, DCMU treatment or a higher PQ/PQH2 ratio attenuated this response (Figure 6C), indicating the pivotal role of the photosynthesis pathway in translational activation in chloroplasts.

**Figure 6.**
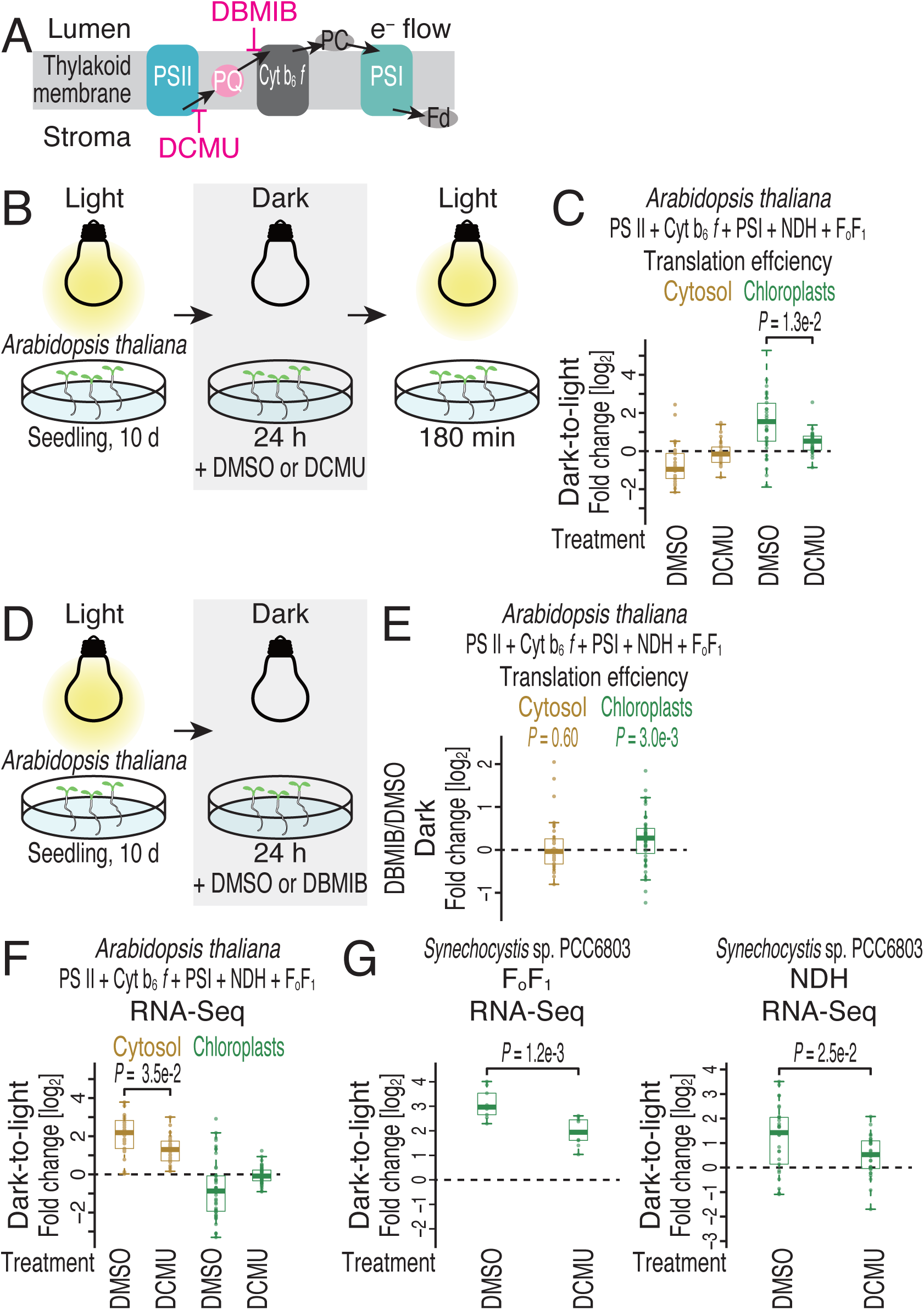
PQH2 triggers translational activation in chloroplasts. (A) Schematic representation of the inhibitory mechanisms of DCMU and DBMIB. (B) Schematic representation of the experimental design used to monitor the response of cytosolic and chloroplastic gene expression to DCMU treatment. (C) Fold change in the translation efficiency following DCMU treatment of *A. thaliana* during the dark-to-light transition. Cytosolic mRNAs and chloroplast mRNAs encoding the subunits of FoF1 ATP synthase, PSII, Cyt *b6f*, PSI, and NDH were analyzed. (D) Schematic representation of the experimental design used to monitor the response of cytosolic and chloroplastic gene expression to DBMIB treatment. (E) Fold change in the translation efficiency following DBMIB treatment of *A. thaliana* in the dark. Cytosolic mRNAs and chloroplast mRNAs encoding the subunits of FoF1 ATP synthase, PSII, Cyt *b6f*, PSI, and NDH were analyzed. (F) Fold changes in RNA abundances following DCMU treatment of *A. thaliana* during the dark-to-light transition. Cytosolic mRNAs and chloroplast mRNAs encoding the subunits of FoF1 ATP synthase, PSII, Cyt *b6f*, PSI, and NDH were analyzed. (G) Fold changes in RNA abundances following DCMU treatment of *Synechocystis* sp. PCC6803 during the dark-to-light transition. mRNAs encoding the subunits of FoF1 ATP synthase (left) and NDH (right) were analyzed. The box plots (C and E-G) show the median (center line), upper/lower quartiles (box limits), and 1.5× interquartile range (whiskers). Significance was determined by the Mann‒Whitney *U* test. See also Figure S7 and Tables S1-2 and S8-10.

Given the decreased PQH2 by DCMU, we evaluated the importance of PQH2 for chloroplastic translation by treatment with 2,5-dibromo-3-methyl-6- isopropylbenzoquinone (DBMIB). This compound inhibits electron flux to Cyt *b6f* ^63–65^ (Figure 6A) and thus maintains the PQ pool in the form of PQH2 (or lower PQ/PQH2 ratio). Indeed, treatment of *Arabidopsis* seedlings with this compound under dark conditions (Figure 6D) induced effects mimicking those of light exposure, as we observed increases in the translation efficiency of complex subunits in chloroplasts (Figures 6E, S7A-B, S7D, and Table S9).

### Coupling with photosynthesis and cytosolic mRNA abundances

On the other hand, our data suggest that PQH2 itself is not involved in signaling controlling cytosolic mRNA abundances. DCMU decreased the efficiency of the induction of cytosolic mRNAs encoding chloroplastic complex subunits during the dark-to-light transition (Figures 6F and S7E). Thus, retrograde signaling—from the chloroplast to the cytoplasm/nucleus—plays a role in this response. On the other hand, DBMIB treatment under dark conditions did not increase the expression of these mRNAs (Figure S7F-G).

To investigate the evolutionary origin of this response, we repeated DCMU treatment in cyanobacteria, in which FoF1 and NDH complex synthesis is controlled at the RNA level (Figures 5A-B). DCMU attenuated the accumulation of the corresponding mRNAs in response to light (Figures 6G, S7H-L, and Table S10), suggesting that photosynthesis is a shared trigger for mRNA accumulation both in cyanobacteria and in the *Arabidopsis* cytosol.

### Ongoing cytosolic translation drives chloroplastic translation activation and cytosolic mRNA accumulation

Considering that cytosolic translation directs mitochondrial translation in yeasts ^13^, we hypothesized that a similar communication between translation systems may play an important role. To address this point, we incubated *Arabidopsis* seedlings with cytosolic translation inhibitor cycloheximide before light exposure (Figure 7A). Strikingly, cytosolic translation shutoff by this compound blocked the translational efficiency activation of chloroplastic mRNAs (Figures 7B, S8A-C, and Table S11). We repeated similar experiments in *Chlamydomonas reinhardtii* and found the same counteraction of chloroplastic translation activation by cycloheximide (Figure 7C, S8D-F, and Table S12). Thus, ongoing cytosolic translation is important for light-dependent chloroplastic translational activation. This dependency on the cytoplasmic translation was quite similar to the mitochondrial translation regulation found in yeasts ^13^, suggesting a similar mechanistic basis behind them. For cytosolic mRNA accumulation upon light exposure, we also observed that productive cytoribosome traversal is necessary in both *Arabidopsis* and *Chlamydomonas* (Figures 7D-E and S8G-H).

**Figure 7.**
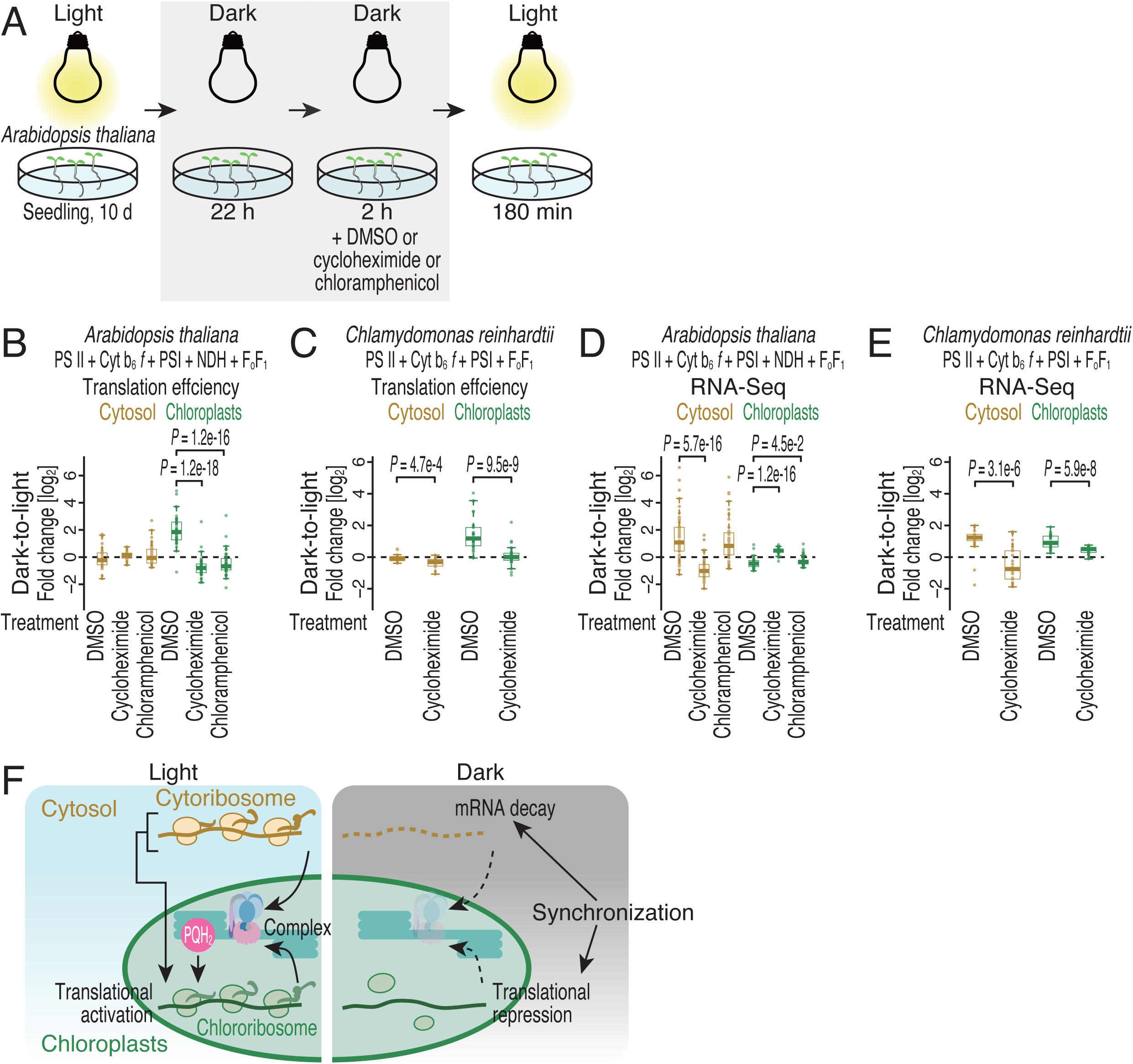
Active cytosolic protein synthesis triggers translational activation in chloroplasts. (A) Schematic representation of the experimental design used to monitor the response of cytosolic and chloroplastic gene expression to translation inhibitor treatment. (B) Fold change in the translation efficiency following translation inhibitor treatment of *A. thaliana* during the dark-to-light transition. Cytosolic mRNAs and chloroplast mRNAs encoding the subunits of FoF1 ATP synthase, PSII, Cyt *b6f*, PSI, and NDH were analyzed. (C) Fold change in the translation efficiency following translation inhibitor treatment of *C. reinhardtii* during the dark-to-light transition. Cytosolic mRNAs and chloroplast mRNAs encoding the subunits of FoF1 ATP synthase, PSII, Cyt *b6f*, and PSI were analyzed. (D) Fold change in RNA abundances following translation inhibitor treatment of *A. thaliana* during the dark-to-light transition. Cytosolic mRNAs and chloroplast mRNAs encoding the subunits of FoF1 ATP synthase, PSII, Cyt *b6f*, PSI, and NDH were analyzed. (E) Fold change in RNA abundances following translation inhibitor treatment of *C. reinhardtii* during the dark-to-light transition. Cytosolic mRNAs and chloroplast mRNAs encoding the subunits of FoF1 ATP synthase, PSII, Cyt *b6f*, and PSI were analyzed. (F) Model of the distinct mechanisms employed for cytoplasmic and chloroplastic gene expression in response to light exposure. The box plots (B-E) show the median (center line), upper/lower quartiles (box limits), and 1.5× interquartile range (whiskers). Significance was determined by the Mann‒ Whitney *U* test. See also Figure S8 and Tables S1-2 and S11-12.

In *Arabidopsi*s, we also conducted reciprocal experiments, shutting off chloroplastic translation by chloramphenicol (Figures 7A-B, 7D, S8A-C, S8G, and Table S11). As expected, this inhibitor did not lead to chloroplastic translation activation by light (Figure 7B and S8G). However, the accumulation of cytosolic mRNAs normally occurred (Figure 7D). We note that chloramphenicol should also block mitochondrial translation. Despite this technical caveat of this inhibitor, the ineffectiveness on cytosolic mRNA accumulation suggested that coordination between the two gene expression systems is only mediated in a unidirectional manner (*i.e.*, cytoplasm to chloroplast).

Considering these data collectively, we concluded that chloroplasts use PQH2 as a signaling molecule to balance the load of complex subunits used in the chloroplast stroma under light/dark conditions (Figure 7F). Moreover, chloroplastic translation activation upon light exposure depends on progressive cytoribosome traversal (Figure 7F). This mechanism provides another checkpoint for the trigger for protein synthesis in chloroplasts, avoiding uneven protein supply from the two translation systems.

## Discussion

Orchestration of the subunit supply via dual gene expression systems is challenging. For this orchestration, in yeasts, cytosolic translation tunes mitochondrial translation ^13^. The present study highlights another mechanism underlying the balance in protein synthesis between dual genomes: control of RNA abundances in the cytosol and control of translation in chloroplasts. Light-dependent chloroplastic translation was initially reported 3 decades ago ^66–68^ and has more recently been assessed at high resolution via chloroplast-tailored ribosome profiling ^32,69^. Our ribosome profiling for simultaneous assessment of translation via dual systems integrated the knowledge of chloroplastic translational regulation into a global view of the economy for multisubunit complexes.

Given the rapid decreases in cytosolic RNA abundances caused by light-to-dark shifts, mRNA decay is the most likely mechanism involved. This decay could be coordinated with transcriptional tuning, which may turn off mRNA synthesis under dark conditions and turn on under light conditions. In contrast to PQH2-mediated mRNA splicing driven by exposure to light ^64,70^, the retrograde regulation of mRNA abundances reported in the present study may be driven by other signaling molecules, such as reactive oxygen species (ROS), isoprenoid precursors [methylerythritol cyclodiphosphate (MEcPP)], carotenoid derivatives, heme, and a phosphonucleotide [3′-phosphoadenosine 5′-phosphate (PAP)] ^71^. Since PAP was reported to function as an inhibitor of the 5′-to-3′ RNA exonuclease XRN2 in retrograde signaling ^72^, future investigations are warranted to determine its involvement.

On the other hand, dark-to-light shift may require transcriptional activation. The requirement of cytosolic translation (Figure 7D-E) could be explained by *de novo* synthesis of transcriptional factors at the beginning of the light exposure, which is associated with photosynthesis (Figure 6F).

In *Arabidopsis*, we observed the accumulation of chloroplastic mRNAs in the dark (Figures 2B-F, 3A, and 4A-C). Although the biological significance was still unclear, we found that the chloroplastic RNA accumulation in the dark was associated with the PQ/PQH2 ratio and cytosolic translation status. While dark-to-light transition reduced the chloroplastic mRNAs, DCMU and cycloheximide counteracted the effects (Figures 6F and 7D). We also observed that DBMIB treatment in the dark mimics the light response in terms of the reduction of chloroplastic mRNA level (Figure S7G). Thus, the chloroplastic mRNA levels were probably controlled by the same mechanism of chloroplastic translation regulation.

In cyanobacteria, changes in mRNA abundances during light/dark transitions have previously been studied. Consistent with our observation, the occurrence of active mRNA decay under dark conditions has been suggested ^73^. Conversely, light exposure inhibits mRNA decay ^73^ and simultaneously induces transcription through two-component regulatory systems involving histidine kinases and response regulators ^74,75^. Importantly, our ribosome profiling results showed that protein synthesis predominantly follows alterations in mRNA abundances, with no further need for translational regulation, at least for the short periods of darkness in our experiments. Longer exposure to dark conditions may trigger translational repression through the stringent response mediated by (p)ppGpp for adaptation ^76^.

Most likely, light-dependent chloroplastic translation is regulated at translation initiation, considering that this process is a global rate-limiting step in protein synthesis in growing cells ^77–83^ and generally determines ribosome footprint numbers on ORFs ^84–86^. Indeed, we found the reduction and the restoration of the basal level of footprints along ORFs in response to dark-light transitions (Figures S2F, S3O, S4N, and S5N). This process may be mediated through mRNA-specific RNA-binding proteins and/or global translational regulators (through interactions with chlororibosomes) ^7^.

In addition to translation initiation, the elongation process can be subject to light-dependent regulation. We observed dynamic changes in cytoribosome and chlororibosome pausing in acute light/dark transitions (Figures S2I, S3R, S4O, S5O, and S6N). *PsbA* (encoding D1, a catalytic center subunit of PSII) is a notable example of chlororibosome pausing ^87,88^. Our ribosome profiling results revealed stalling at this position (after exposure of the 4th transmembrane domain), consistent with earlier reports in *Arabidopsis* ^87,88^ (Figure S2J-K). Since this pausing on *psbA* mRNA may exhibit interspecies variation (Figures S3S, S4P, S5P, and S6O), analysis of chloroplastic translation by disome profiling ^27,89–92^ will be useful.

## Limitations of the study

We generally observed a wider range of chlororibosome footprints (Figures 1C, S2A, S3B, S4B, and S5B) than cytoribosome footprints. The diversity of chlororibosome footprints may be related to the conformational varieties of chlororibosomes. Indeed, even broader distribution of footprints could be found in bacterial ribosomes ^22,26^ and attributed to conformational diversity that could be reflected in RNase digestion. We also found species-dependent differences in chlororibosome footprint length (Figures 1C, S2A, S3B, S4B, and S5B). This could be associated with the conformation/composition of chlororibosomes in each species. Given that mitoribosomes have quite different structures/factor incorporations among species ^12,93^, it is not surprising that similar diversity occurs in chlororibosomes. Future studies surveying the structure-footprint relationship will provide more concrete rationales for the chlororibosome footprint heterogeneity.

## Resource availability

### Lead contact

Requests for further information and resources should be directed to and will be fulfilled by the lead contact, Shintaro Iwasaki.

### Materials availability

The regents, commercial assays, model organisms/strains, software, and equipment can be found in the key resources table. Further information and requests for resources and reagents should be directed to and will be available upon completion of appropriate material transfer agreements by the lead contact, Shintaro Iwasaki.

### Data and code availability

The ribosome profiling and RNA-Seq data obtained in this study (GSE267365) were deposited in the National Center for Biotechnology Information (NCBI) database. We also used bacterial ribosome profiling data published in an earlier work (GSE180482) ^94^. All the GEO-deposited data were adapter-trimmed and demultiplexed according to linker sequences.

For ribosome profiling and RNA-Seq data analysis, we deposited codes used in this study [Zenodo 15139971 (https://zenodo.org/records/15139971)].

## Acknowledgments

We are grateful to all the members of the Iwasaki laboratory for fruitful discussion and technical help. We also thank Dr. Hiro-oki Iwakawa for the constructive advice. *Synechocystis* sp. PCC6803 is a kind gift from Dr. Kan Tanaka. The supercomputer HOKUSAI Sailing Ship in RIKEN was used for computations. This study was supported by the Japan Society for the Promotion of Science (JSPS) (JP19H02959, JP23H02415, and JP23H00095 to S.I.; JP19J14480 to T.F.; JP23KJ0444 to T.W.; JP23KJ2175 to N.K.; JP22K20765, JP23KJ2178, and JP23K14173 to H.T.; JP19K22434 and JP21H02509 to T.H.; JP23H05473 and JP23H04882 to M.Y.); the Ministry of Education, Culture, Sports, Science and Technology (MEXT) (JP20H05784 and JP24H02307 to S.I.); RIKEN (Pioneering Project “Biology of Intracellular Environments” to S.I. and M.M.; Incentive Research Project to H.T. and N.K.); and The University of Tokyo (C2205 and C2306 to T.W.). Some of the DNA libraries were sequenced by QB3 Genomics, UC Berkeley, Berkeley, CA (RRID: SCR_022170), which facilities were supported by an NIH Instrumentation Grant (S10 OD018174). T.F. was a recipient of the RIKEN Junior Research Associate Program and a JSPS Research Fellow (DC2). T.W. received fellowships from JSPS (DC2), JST SPRING (JPMJSP2108), and the ANRI. H.T. was a JSPS Research Fellow (PD). N.K. received the support of JSPS Research Fellow (PD) and the RIKEN Special Postdoctoral Researchers Program.

## Author Contributions

Conceptualization: T.F. and S.I.;

Methodology: T.W., T.F., N.K., Y.K., Y.H., T.H., H.T., A.H., E.M.-S., and S.I.;

Formal analysis: T.W., T.F., N.K., H.T., and S.I.;

Investigation: T.W., T.F., N.K., Y.K., Y.H., T.H., H.T., T.K., A.H., and E.M.-S.;

Writing – Original Draft: N.K. and S.I.;

Writing – Review & Editing: T.W., T.F., N.K., Y.K., Y.H., T.H., H.T., T.K., A.H., E.M.- S., K.M., M.Y., M.M., and S.I.;

Visualization: T.W., T.F., N.K., and S.I.;

Supervision: Y.K., Y.H., T.H., K.M., M.Y., M.M., and S.I.;

Funding Acquisition: T.W., T.F., N.K., T.H., H.T., M.Y., M.M., and S.I.

## STAR Methods

### Experimental model and study participant details

#### Culture of A. thaliana

Sterilized *A. thaliana* Col-0 (Arabidopsis Biological Resource Center) seeds were sown on growth medium [1× Murashige and Skoog salt mixture (SHIOTANI M.S. CO), 10 g/l sucrose, and 0.8% agar; adjusted to pH 5.8], stored in the dark at 4°C for 3 d, and subsequently grown at 23°C under continuous exposure to white light (50 µmol/m^2^/s) for 10 d. The plants were subsequently harvested before initiation of light/dark transitions (0 min); 30, 60, and 180 min after the light-to-dark transition; and 30, 60, and 90 min after the dark-to-light transition. The samples were flash-frozen in liquid nitrogen and stored at −80°C.

For photosynthesis inhibitor treatment, 20 ml of the drug solution [10 mM MES (pH 5.8) with 10 µM DCMU [nacalai tesque], 30 µM DBMIB [nacalai tesque], or a corresponding amount of DMSO (solvent)] was poured onto the growth medium, and the drug-treated seedlings were subsequently grown in the dark at 23°C for 24 h. For the experiments with DCMU treatment, the plants were further incubated in the light for 180 min after the dark period.

For translation inhibitor treatment, 10-d seedlings were transferred to dark conditions and incubated for 21 h. Ten milliliters of the drug solution [10 mM MES (pH 5.8) with 100 µM cycloheximide (Sigma–Aldrich), 100 µM chloramphenicol (FUJIFILM Wako Pure Chemical Corporation), or a corresponding amount of DMSO (solvent, 0.028%)] ^95^ was poured onto the growth medium. After 3-h incubation, seedlings were harvested before or after incubation in the light for 180 min.

#### Culture of B. distachyon

*B. distachyon* (Bd21, psb00001, RIKEN BioResource Center) was grown on soil at 23°C on a 14-h light/10-h dark cycle for 25 d, followed by constant light conditions for 10 d. The plants were harvested before initiation of light/dark transitions (0 min); 30, 60, and 180 min after the light-to-dark transition; and 30, 60, and 90 min after the dark-to-light transition. The aboveground tissues from ten plants were pooled to generate one sample, flash-frozen in liquid nitrogen, and stored at −80°C.

#### Culture of C. reinhardtii

*C. reinhardtii* (NIES-2235) was obtained from the National Institute of Environmental Studies (NIES) culture collection. The cells were cultured in 50 ml of TAP medium ^96^ in a 100-ml conical flask at 26°C with shaking at 100 rpm under constant exposure to white fluorescent light (MIR-254, PHCbi). The cells in a 30-ml volume of culture were collected on a 0.45-μm filter (Millipore) by vacuum filtration. The collected cells on the filter were immediately frozen in liquid nitrogen and stored at −80°C. Cultures were harvested before initiation of light/dark transitions(0 min); 15 and 90 min after the light-to-dark transition; and 15 and 90 min after the dark-to-light transition.

For translation inhibitor treatment, 200 ml of *C. reinhardtii* culture in 500-ml conical flask were transferred to dark conditions, incubated for 22 h, and treated with 20 µl of the drug solution [100 mg/ml cycloheximide or a corresponding amount of DMSO (solvent)] ^97^ for 2 h. Thirty milliliters of the culture were harvested before or after incubation in the light for 90 min.

#### Culture of C. paradoxica

*C. paradoxa* (NIES-547) was obtained from the NIES culture collection. The cells were aseptically cultured in 6 square glass flasks that contained 800 ml of Bold 3N liquid medium for 14 d at 26°C under constant exposure to white fluorescent light (10 μE/m^2^/s) with 1% CO2 (v/v) bubbling ^98^. The cells in a 200-ml volume of culture were collected on two 0.45-μm filters (Millipore) by vacuum filtration. The collected cells on the filters were immediately frozen in liquid nitrogen and stored at −80°C. Cultures were harvested before initiation of light/dark transitions (0 min); 15 and 90 min after the light-to-dark transition; and 15 and 90 min after the dark-to-light transition.

#### Culture of Synechocystis sp. PCC6803

*Synechocystis* sp. PCC6803 (kindly shared by Dr. Kan Tanaka) ^99^ was cultivated in BG- 11 liquid medium ^100^ at 30°C under continuous exposure to white light (50 µmol/m^2^/s) with shaking (160 rpm). The cells in a 30-ml volume of culture were collected on a 0.45- μm filter (Millipore) by vacuum filtration. The collected cells on the filter were immediately frozen in liquid nitrogen and stored at −80°C. Cultures were harvested before initiation of light/dark transitions (0 min); 15 and 90 min after the light-to-dark transition; and 15 and 90 min after the dark-to-light transition.

For compound treatment, cells were cultured in 30 ml of medium containing 10 µM DCMU, 10 µM DBMIB, or a corresponding amount of DMSO (solvent). The drug-treated cells were incubated in the dark at 30°C for 24 h. For the experiments with DCMU treatment, the cells were further incubated in the light for 90 min after the dark period.

### Method details

#### Lysate preparation for ribosome profiling and RNA-Seq analysis

For *A. thaliana* and *B. distachyon*, frozen samples were pulverized with 400 µl of frozen lysis buffer [100 mM Tris-HCl (pH 7.5), 40 mM KCl, 20 mM MgCl2, 1 mM dithiothreitol (DTT), 1% Triton X-100, 300 µg/ml chloramphenicol, and 100 µg/ml cycloheximide] using a Multi-beads Shocker instrument (Yasui Kikai) at 2800 rpm for 10 s. For MNase digestion, 5 mM CaCl2 was added to the lysis buffer.

For *C. reinhardtii*, pulverization was performed by 2 cycles at 1500 rpm for 15 s with 650 µl of frozen lysis buffer. The lysate was thawed on ice and immediately clarified by centrifugation at 3,000 × g for 5 min at 4°C. The supernatant was incubated with 25 U/ml Turbo DNase (Thermo Fisher Scientific) and clarified by centrifugation at 20,000 × g for 10 min at 4°C. For MNase digestion, 5 mM CaCl2 was added to the lysis buffer.

For *C. paradoxa*, pulverization was performed with 650 µl of frozen lysis buffer at 2800 rpm for 10 s. The lysate was thawed on ice and filtered through a Spin-X centrifuge tube filter 0.22-µm (Costar) by centrifugation at 16,000 × g for 5 min at 4°C. Then, the flowthrough was incubated with 25 U/ml Turbo DNase and clarified by centrifugation at 20,000 × g for 10 min at 4°C.

For *Synechocystis* sp. PCC6803, pulverization was performed by 2 cycles of 1500 rpm for 15 s with 400 µl of frozen bacterial lysis buffer [20 mM Tris-HCl (pH 7.5), 150 mM NH4Cl, 10 mM MgCl2, 5 mM CaCl2, 1 mM DTT, 1% Triton X-100, and 100 µg/ml chloramphenicol]. The lysate was thawed on ice and immediately clarified by centrifugation at 3,000 × g for 5 min at 4°C. The supernatant was incubated with 25 U/ml Turbo DNase on ice and clarified by centrifugation at 20,000 × g for 10 min at 4°C.

#### Library preparation for ribosome profiling

Library preparation for ribosome profiling was performed as previously described ^101^. The RNA content in each lysate was measured with a Qubit RNA BR Assay Kit (Thermo Fisher Scientific). The lysates were treated with 20 U of RNase I (LGC Biosearch Technologies) for *A. thaliana* (30 µg of RNA), *B. distachyon* (30 µg of RNA), and *C. paradoxa* (30 µg of RNA); with 10 U of RNase I for *C. reinhardtii* (20 µg of RNA); or with 15 U of MNase (Roche) for *Synechocystis* sp. PCC6803 (20 µg of RNA). All of the above enzymatic treatments were performed in a 300-µl reaction (scaled up by the amount of lysis buffer or bacterial lysis buffer) at 25°C for 45 min. After the ribosomes were collected on a sucrose cushion, RNA was extracted with TRIzol (Thermo Fisher Scientific) and a Direct-zol RNA MicroPrep Kit (Zymo Research). After separation on a 15% urea polyacrylamide gel electrophoresis (PAGE) gel, RNA fragments containing 17 to 50 nt were excised from the gel, dephosphorylated, and ligated with linkers. rRNAs were depleted with the following kits: a Ribo-Zero Gold rRNA Removal Kit (Plant Leaf) (Illumina) for *A. thaliana, C. reinhardtii*, and *B. distachyon*; a RiboMinus Plant Kit for RNA-Seq (Thermo Fisher Scientific) for *C. paradoxa*; and a Ribo-Zero Gold rRNA Removal Kit (Bacteria) (Illumina) or Pan-Bacteria riboPOOLs (siTOOLs Biotech) for *Synechocystis* sp. PCC6803. cDNAs were synthesized by reverse transcription, subjected to circular ligation, and amplified by PCR. The DNA libraries were sequenced on the HiSeq 4000, HiSeq X, or NovaSeq 6000 platform. The libraries used for this study are summarized in Table S2.

For the experiments with titrated RNase (Figure S1A-D), *A. thaliana* lysates containing 30 µg of total RNA and *C. reinhardtii* lysates containing 20 µg of total RNA were used. For RNase I, 20, 40, or 80 U of the enzyme for *A. thaliana* and 10, 20, or 40 U for *C. reinhardtii* was used. For MNase, lysates were scaled up to 300 µl with lysis buffer containing 5 mM CaCl2, treated with 10, 20, or 40 U of the enzyme, and then quenched by the addition of 20 µl of 0.1 M EGTA.

#### Library preparation for RNA-Seq

Total RNA was extracted from the same lysates used for ribosome profiling with TRIzol LS (Thermo Fisher Scientific) and a Direct-zol RNA MicroPrep Kit (Zymo Research). After rRNA depletion (see the “*Library preparation for ribosome profiling*” section), the libraries were prepared with a TruSeq Stranded mRNA Library Prep Kit (Illumina) or SEQuoia Express Standard RNA Library Prep Kit (Bio-Rad) and sequenced on the HiSeq 4000 or HiSeq X platform. The libraries used for this study are summarized in Table S2.

### Quantification and statistical analysis

#### Data analysis

The deep sequencing data were processed as reported previously with modifications ^102^. Read quality filtering and adapter trimming were performed with fastp (version 0.21.0) ^103^, with the following parameters: -a AGATCGGAAGAGCACACGTCTGAA to trim adapter sequences, --length_limit 149 to discard reads longer than 149 bp, -w 12 to enable multithreaded processing with 12 threads. After all reads aligned to noncoding RNAs were removed using STAR (version 2.7.0a) ^104^, the unaligned reads were mapped to the corresponding nuclear, mitochondrial, and chloroplastic genomes [*A. thaliana*, TAIR10; *B. distachyon*, GCA_000005505.4 (nuclear genome) and NC_011032.1 (chloroplast genome); *C. reinhardtii*, GCA_000002595.3 (nuclear genome), U03843.1 (mitochondrial genome), and NC_005353.1 (chloroplast genome); *C. paradoxa*, ASM443141v1 (nuclear genome), NC_017836.1 (mitochondrial genome), and NC_001675.1 (chloroplast genome); and *Synechocystis* sp. PCC6803, ASM972v1] using STAR (version 2.7.0a). For *Escherichia coli*, reads were first aligned to noncoding RNAs using Bowtie2 (version 2.4.1) ^105^, and these aligned reads were excluded from the downstream analysis. The remaining reads were mapped to the *E. coli* genome sequence (NC_000913.2) with Bowtie2 (version 2.4.1). For ribosome profiling, the A-site position offsets from the 5′ ends of the reads were empirically determined for each read length, as summarized in Table S2. For RNA-Seq analysis, an offset of 15 was applied.

Reads assigned to ORFs were counted with a custom script (https://github.com/ingolia-lab/RiboSeq), and reads whose A sites were located in the first and last 5 codons were excluded. The relative enrichment of ribosome profiling and RNA-Seq reads was calculated with the DESeq2 package (version 1.34.0) ^106^. Translation efficiency was calculated via a generalized linear model with the DESeq2 package (version 1.34.0). The detailed scripts are found in Zenodo [Zenodo 15139971 (https://zenodo.org/records/15139971)].

To compute the ribosome occupancy, the reads of each codon were normalized to the average reads per codon of the corresponding transcript. We note that library L72 (*Brachypodium distachyon*, dark 180 min) was excluded from the codon-wise analysis, due to the low reproducibility (Figure S3G).

The periodicity score was calculated as reported previously ^20^. The relative entropy of *Hj* for read length *j* was defined as follows:

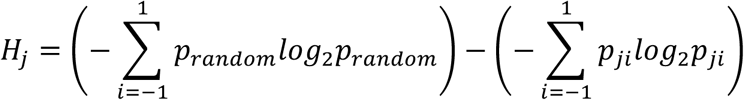

where *pji* is the fraction of ribosomes with footprint length *j* at frame *i,* and *prandom* is 0.333. Then, the periodicity score (*P*) was defined as follows:

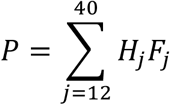

where *Fj* represents the fraction of ribosomes with footprint length *j*.

To calculate reads corresponding to the introns, RNA-Seq reads mapped to the *A. thaliana* chloroplastic genome were counted by featureCounts (version 2.0.3) ^107^. Reads per kilobase (RPK) values for the sum of the reads of introns and those from exon-intron junctions were normalized by RPK values for reads of introns, exon–intron junctions, and exons.

To evaluate RNA editing rate, RNA-Seq reads were aligned to the chloroplast reference genome. Previously characterized chloroplast RNA editing sites ^55^ were considered in the analysis. For each sample, the “*mpileup*” command of SAMtools (version 1.10) ^108^ was employed to quantify the number of reads possessing a C or T base at the corresponding positions.

---

**Figure S1.**
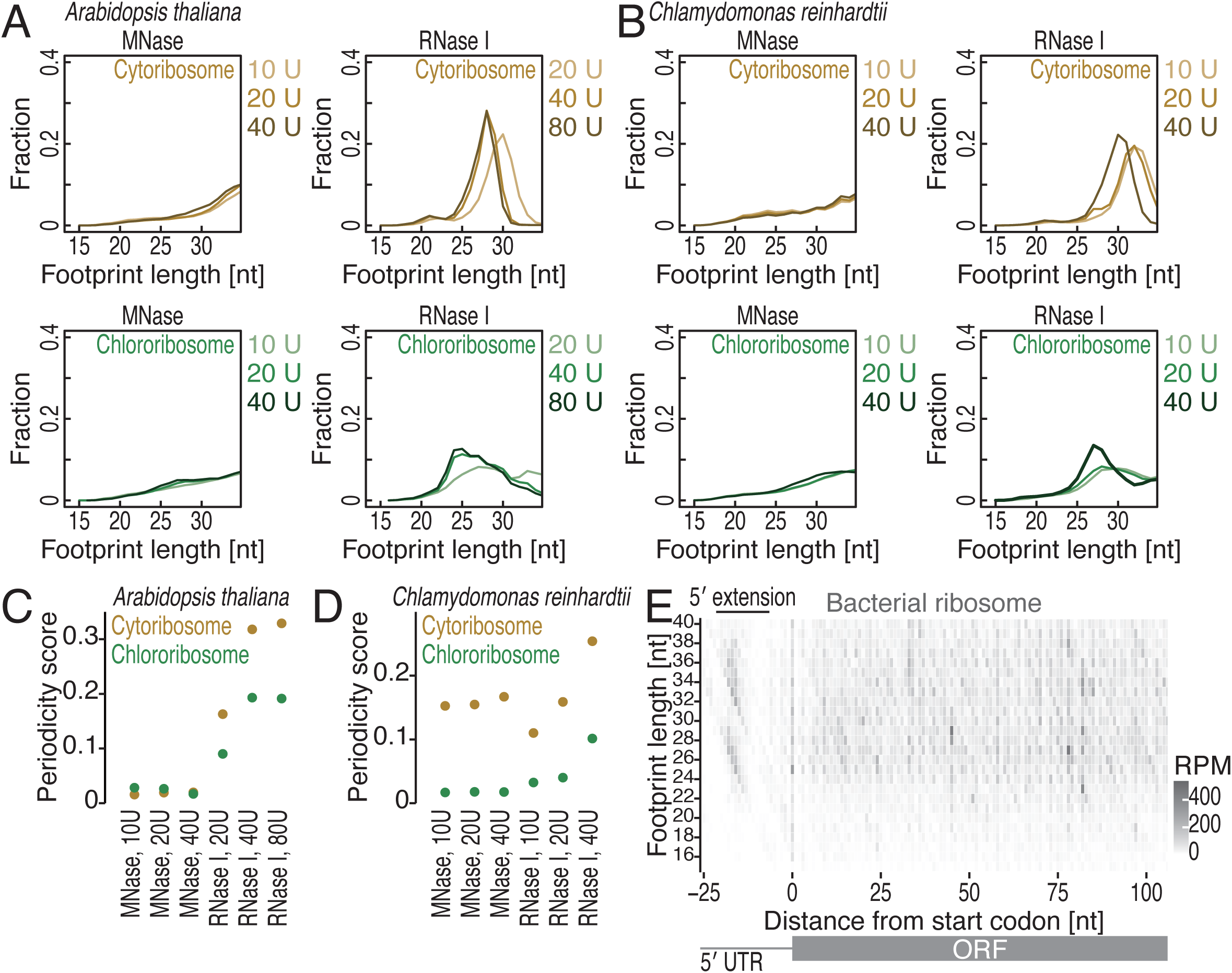
Optimization of plant ribosome profiling, related to Figure 1. (A-B) Length distributions of cytoribosome footprints and chlororibosome footprints in *A. thaliana* (A) and *C. reinhardtii* (B). The indicated RNase conditions were tested for library preparation. (C-D) Periodicity scores in the indicated RNase treatment conditions. (E) Metagene plots of the 5′ ends of bacterial ribosome footprints along the length around start codons (the first nucleotide of the start codon was set to 0). The color scale indicates the read abundance. RPM, reads per million mapped reads.

**Figure S2.**
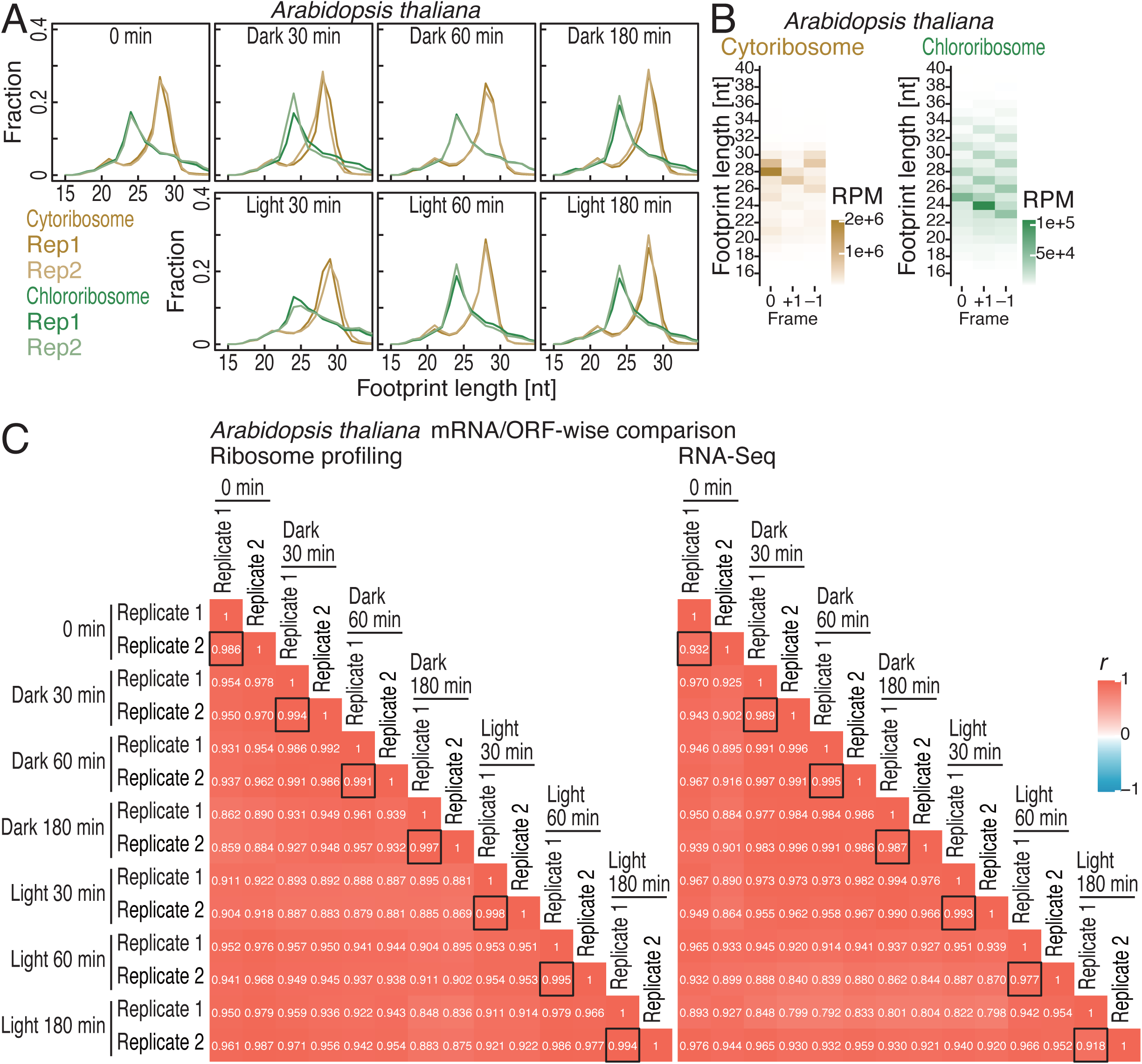
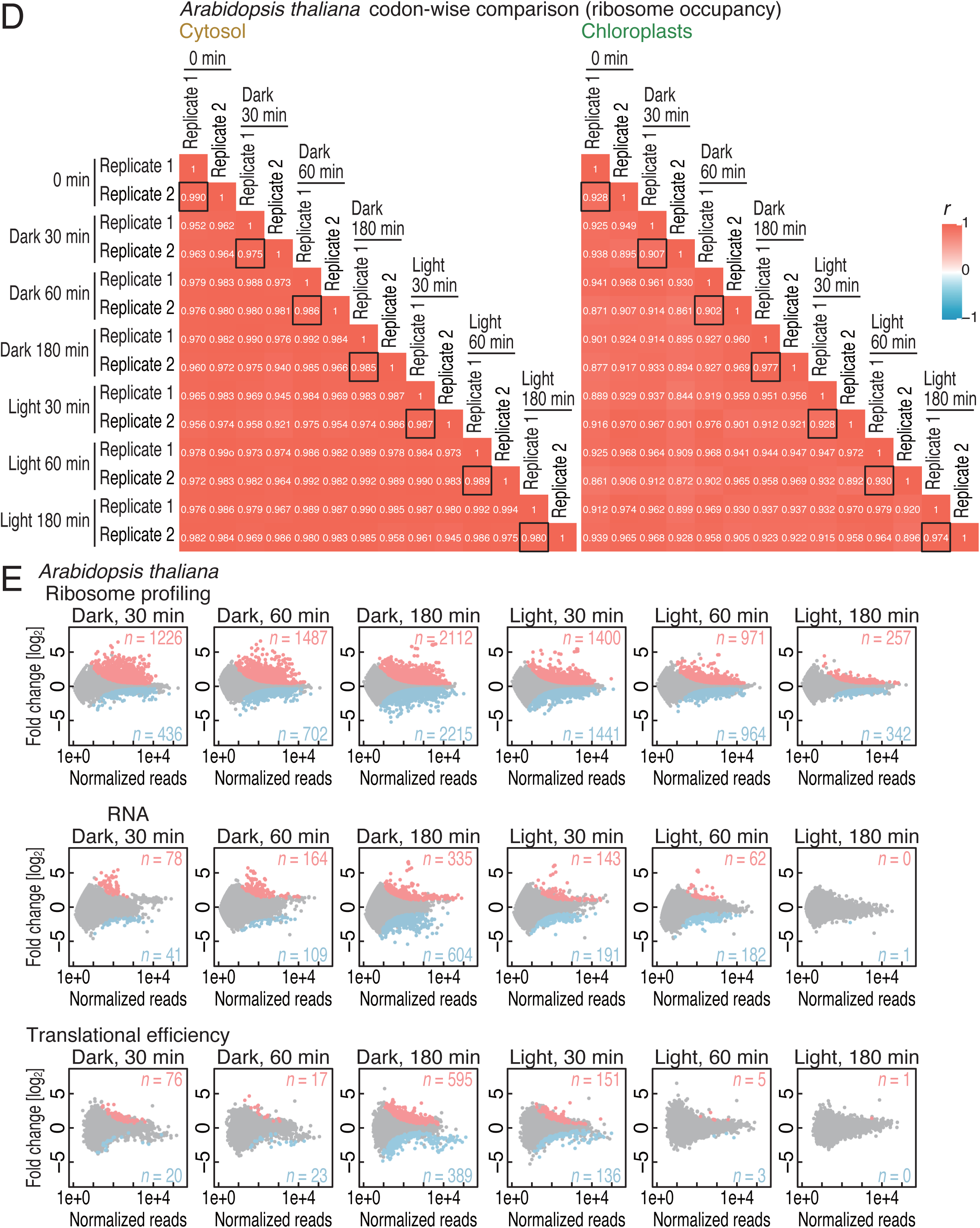
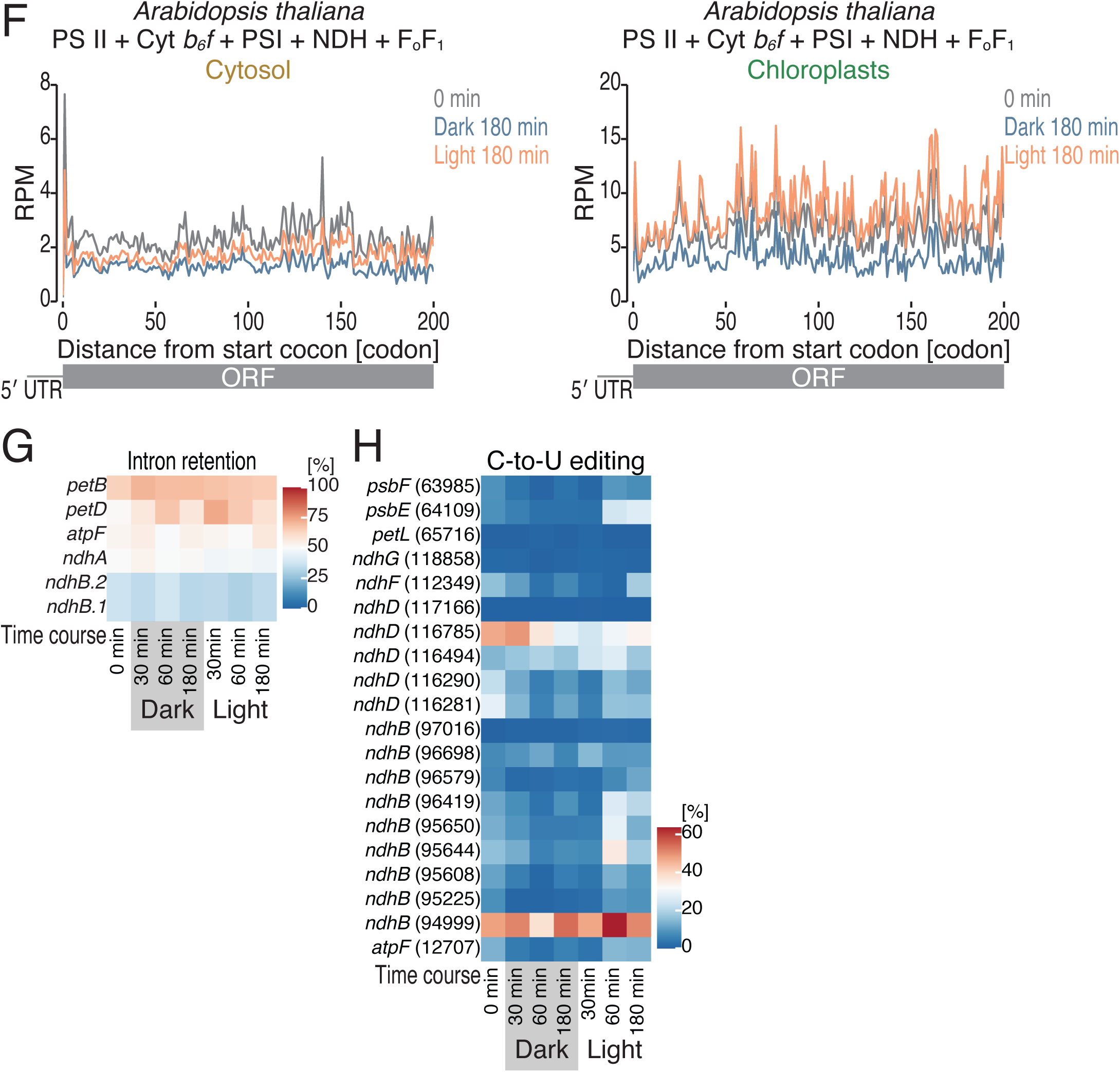
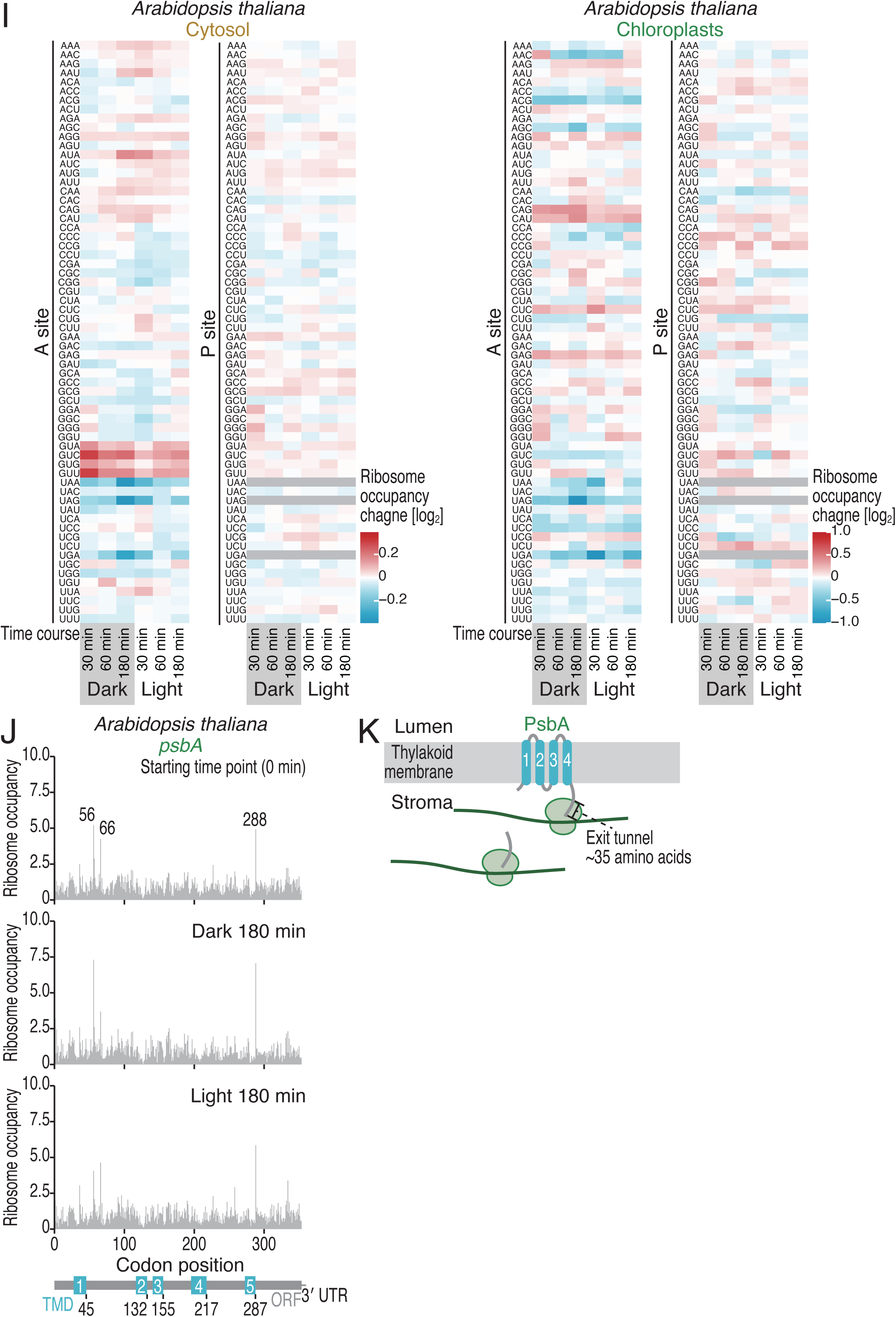
Characterization of ribosome profiling data in *A. thaliana*, related to Figures 1-2. (A) Length distributions of cytoribosome footprints and chlororibosome footprints in *A. thaliana* under the indicated conditions. The data for “0 min” were the same for Figure 1C. (B) Tile plots of corresponding reading frames for the 5′ end of cytoribosome footprints (left) and chlororibosome footprints (right). The color scale indicates the read abundance. RPM, reads per million mapped reads. (C) Heatmap of the Pearson’s correlation coefficients (*r*) of the ORF-mapped reads in ribosome profiling and RNA-Seq under the indicated conditions. The value of *r* is indicated by the color scale. The black box tiles present the correlations of replicates. (D) Heatmap of the Pearson’s correlation coefficients (*r*) of the cytoribosome footprints and chlororibosome footprints on each codon under the indicated conditions. The value of *r* is indicated by the color scale. The black box tiles present the correlations of replicates. (E) MA (M, log ratio; A, mean average) plot for changes in ribosome profiling, RNA-Seq, and translation efficiency in the indicated conditions. Transcripts with significant changes (BH-adjusted *P* value < 0.05) are highlighted. (F) Metagene plots for cryotribosome and chlororibosome footprints (the A site position) around start codons in the indicated conditions. mRNAs encoding the subunits of FoF1 ATP synthase, PSII, Cyt *b6f*, PSI, and NDH were analyzed. (G) A heatmap for intron retention in the indicated conditions. The color scale indicates the percentage of intron retention. (H) A heatmap for C-to-U editing rate in the indicated conditions. The color scale indicates the percentage of editing. (I) Heatmaps for the cytoribosome and chlororibosome occupancy changes at A-site and P-site codons in the indicated conditions. The color scale indicates ribosome occupancy change. (J) Distributions of chlororibosome footprints along the *psbA* mRNA under the indicated conditions. The A-site position of each read (the start codon was set to 0) is indicated. (K) Schematic representation of configurations of chlororibosome pause sites found in *psbA* with cotranslational insertion of the nascent peptide into the thylakoid membrane.

**Figure S3.**
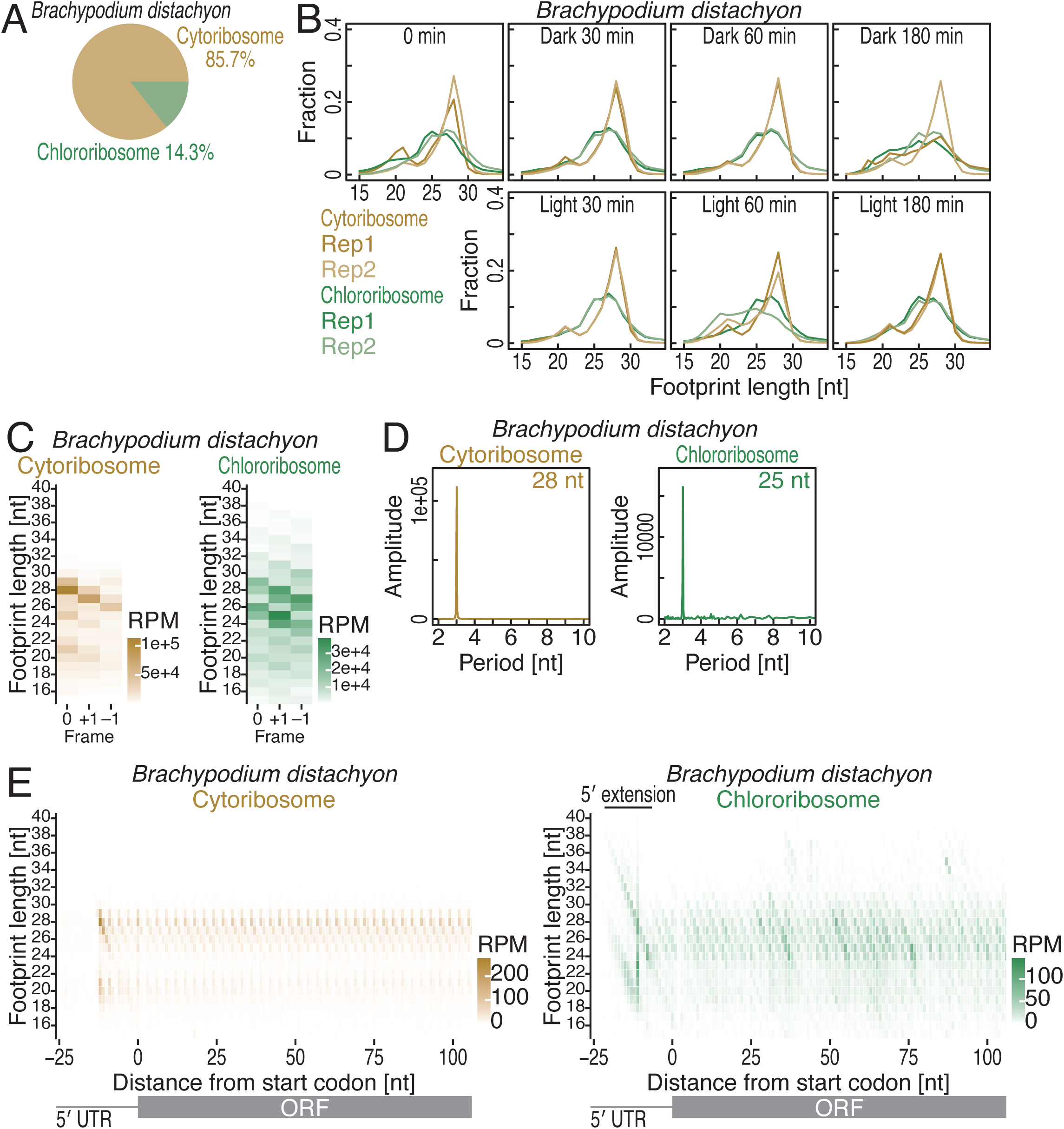
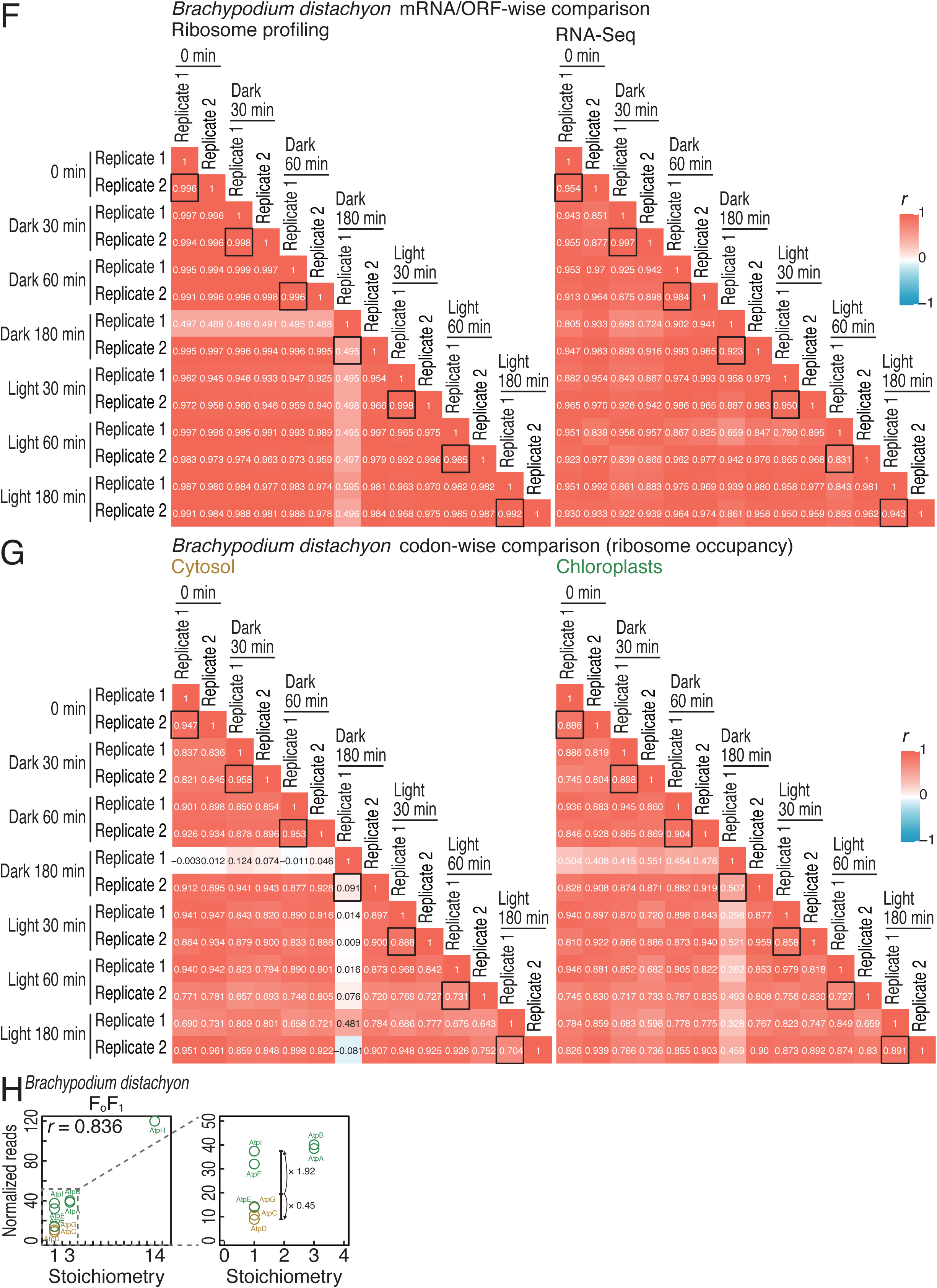
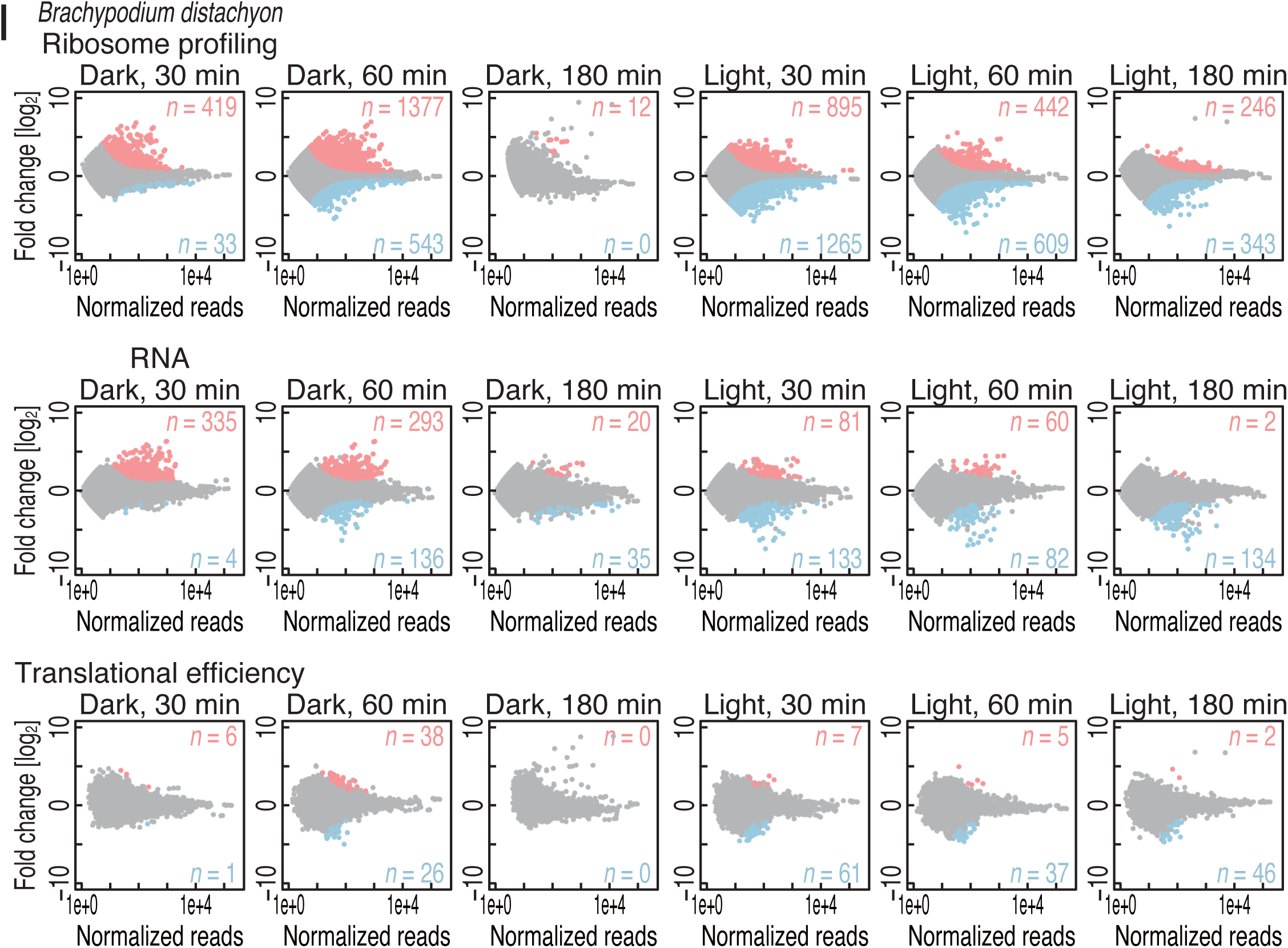
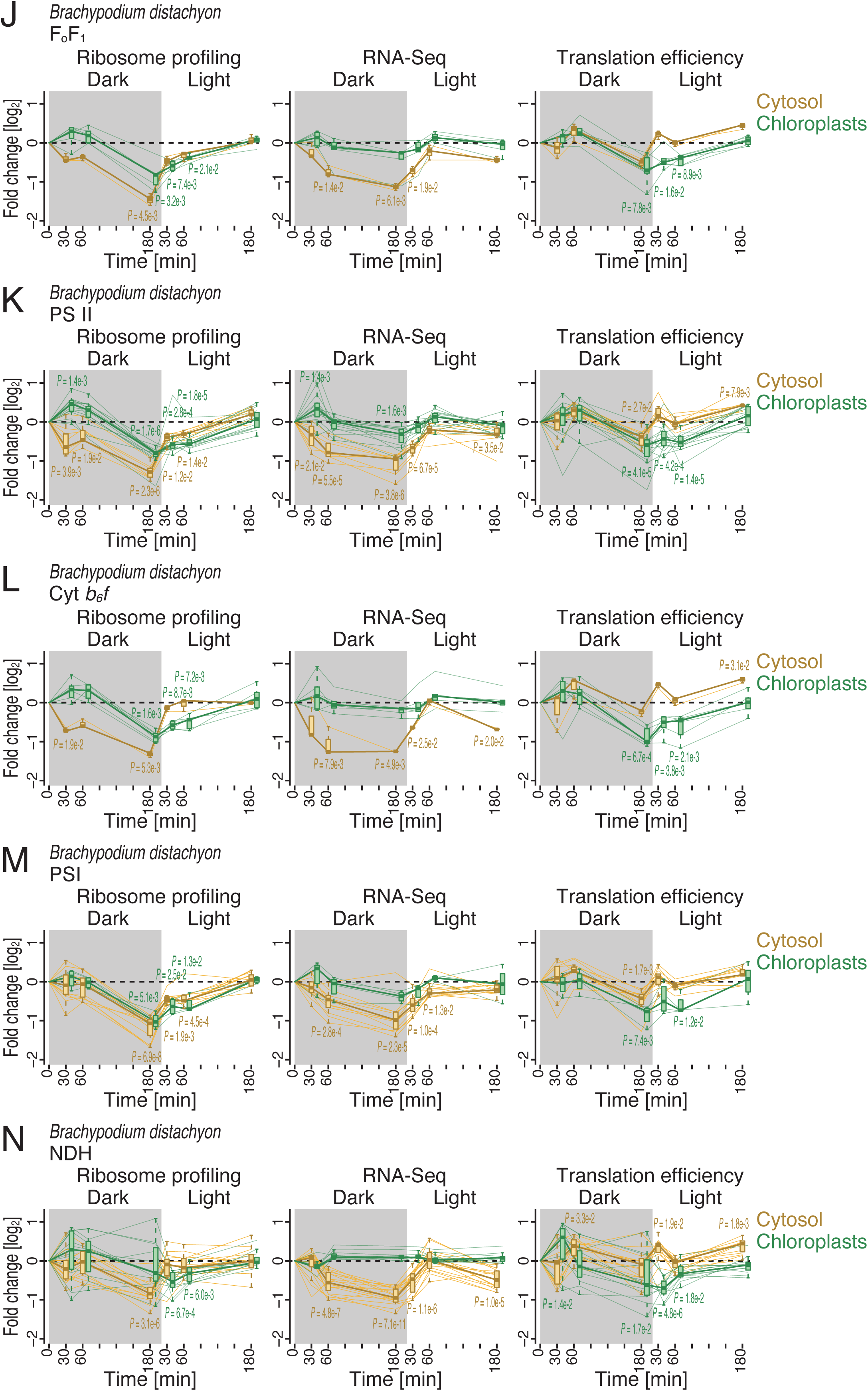
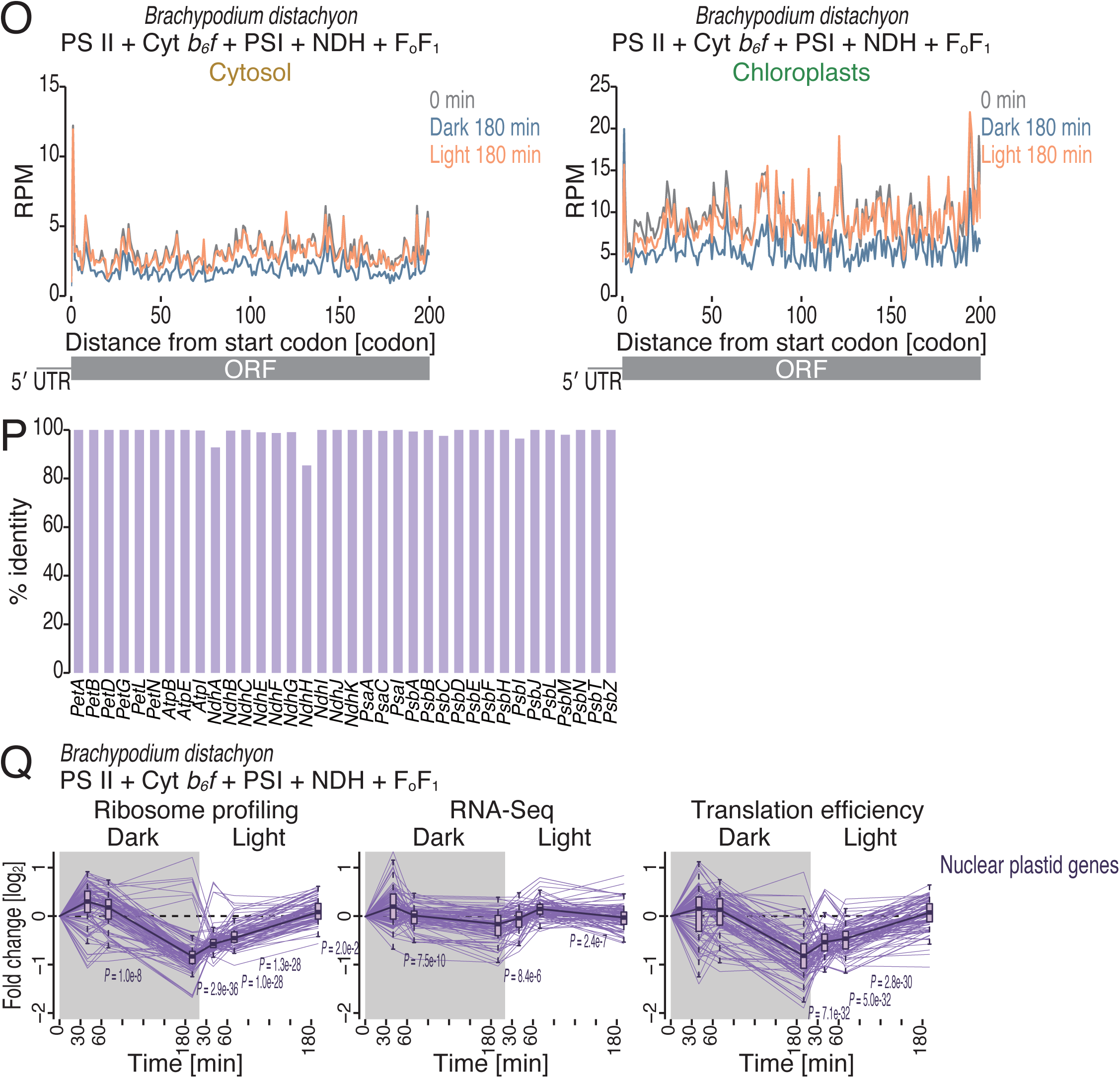
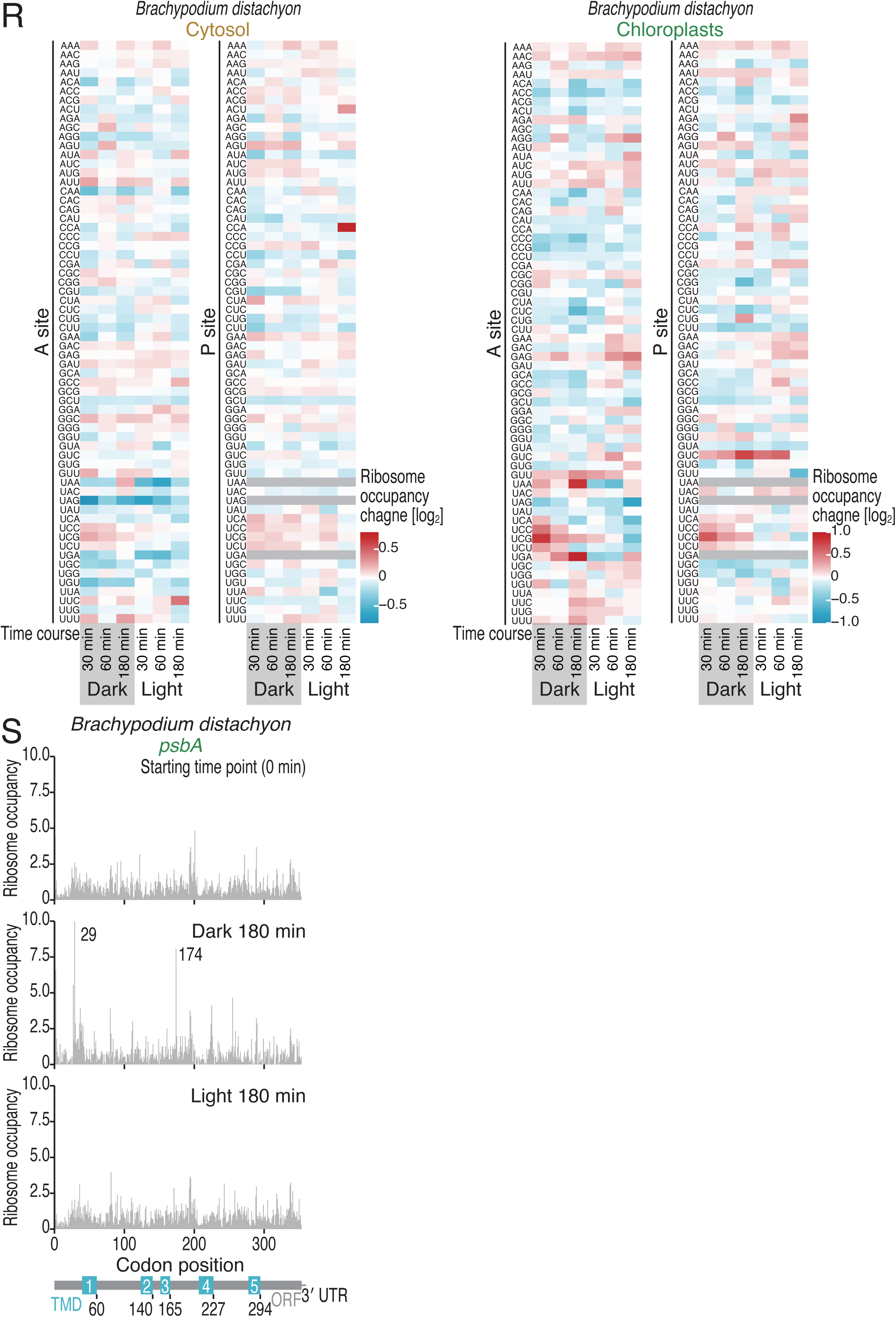
Characterization of ribosome profiling data in *B. distachyon*, related to Figure 3. (A) Pie chart showing the proportions of footprints from cytoribosomes and chlororibosomes in *B. distachyon*. Mitoribosome footprints were not assigned due to a lack of the mitochondrial genome sequence. (B) Length distributions of cytoribosome footprints and chlororibosome footprints in *B. distachyon* under the indicated conditions. (C) Tile plots of corresponding reading frames for the 5′ end of cytoribosome footprints (left) and chlororibosome footprints (right). The color scale indicates the read abundance. RPM, reads per million mapped reads. (D) Discrete Fourier transform of cytoribosome footprints (28 nt) (left) and chlororibosome footprints (25 nt) (right) downstream of the start codon (from position −9 to position 300). (E) Metagene plots of the 5′ ends of cytoribosome footprints (left) and chlororibosome footprints (right) along the length around start codons (the first nucleotide of the start codon was set to 0). The color scale indicates the read abundance. RPM, reads per million mapped reads. (F) Heatmap of the Pearson’s correlation coefficients (*r*) of the ORF-mapped reads in ribosome profiling and RNA-Seq under the indicated conditions. The value of *r* is indicated by the color scale. The black box tiles present the correlations of replicates. (G) Heatmap of the Pearson’s correlation coefficients (*r*) of the cytoribosome footprints and chlororibosome footprints on each codon under the indicated conditions. The value of *r* is indicated by the color scale. The black box tiles present the correlations of replicates. (H) Correlation between the normalized read density (reads/codon) and the subunit stoichiometry for the chloroplast FoF1 ATP synthase, with the zoomed-in inset highlighting data for one component factor. (I) MA (M, log ratio; A, mean average) plot for changes in ribosome profiling, RNA-Seq, and translation efficiency in the indicated conditions. Transcripts with significant changes (BH-adjusted *P* value < 0.05) are highlighted. (J-N) Changes in ribosome profiling, RNA-Seq, and translation efficiency during the light-to-dark and dark-to-light transitions for the subunits of FoF1 ATP synthase (J), PSII (K), Cyt *b6f* (L), PSI (M), and NDH (N) in *B. distachyon*. (O) Metagene plots for cryotribosome and chlororibosome footprints (the A site position) around start codons in the indicated conditions. mRNAs encoding the subunits of FoF1 ATP synthase, PSII, Cyt *b6f*, PSI, and NDH were analyzed. (P) Percent identity between chloroplastic genome-encoded genes and nuclear plastid genes. Based on local pairwise alignments, percent identity was calculated as 100 × identical positions / (aligned positions + internal gap positions). (Q) Changes in ribosome profiling, RNA-Seq, and translation efficiency during the light-to-dark and dark-to-light transitions for the subunits of FoF1 ATP synthase, PSII, Cyt *b6f*, PSI, and NDH among the nuclear plastid genes in *B. distachyon*. (R) Heatmaps for the cytoribosome and chlororibosome occupancy changes at A-site and P-site codons in the indicated conditions. The color scale indicates ribosome occupancy change. (S) Distributions of chlororibosome footprints along the *psbA* mRNA under the indicated conditions. The A-site position of each read (the start codon was set to 0) is indicated. The box plots (J-N and Q)show the median (center line), upper/lower quartiles (box limits), and 1.5× interquartile range (whiskers). The black line indicates the median at each time point.

**Figure S4.**
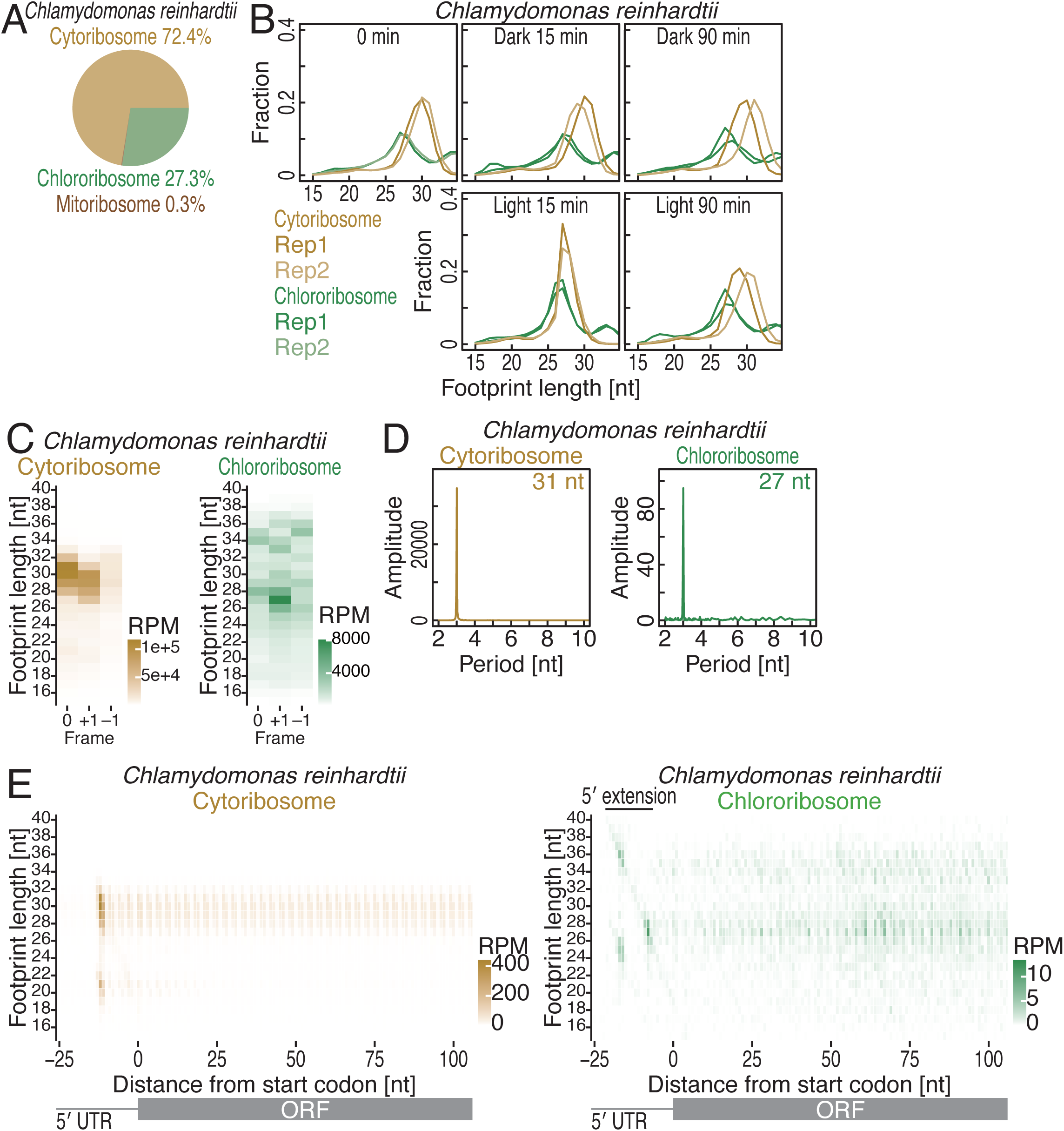
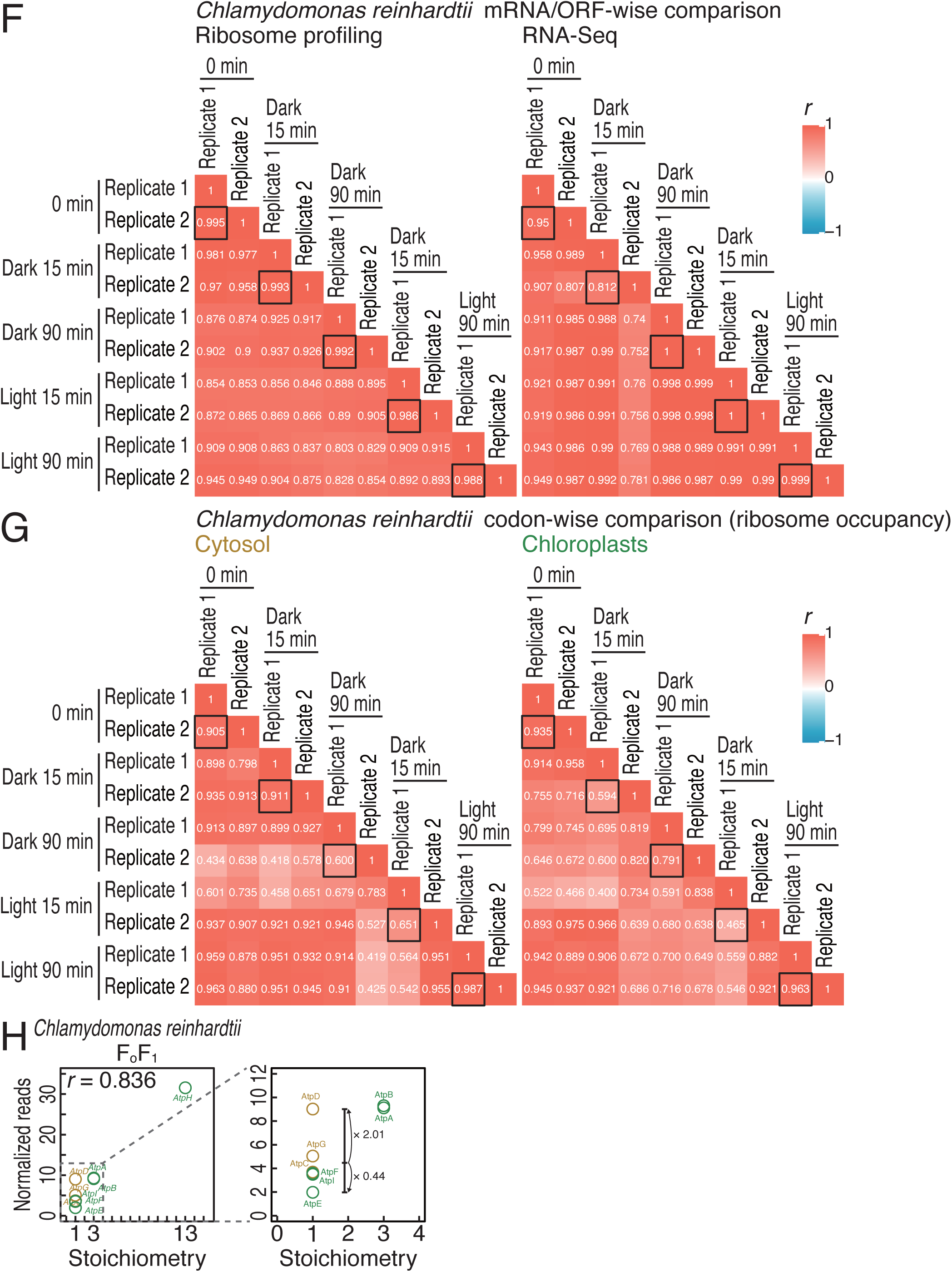
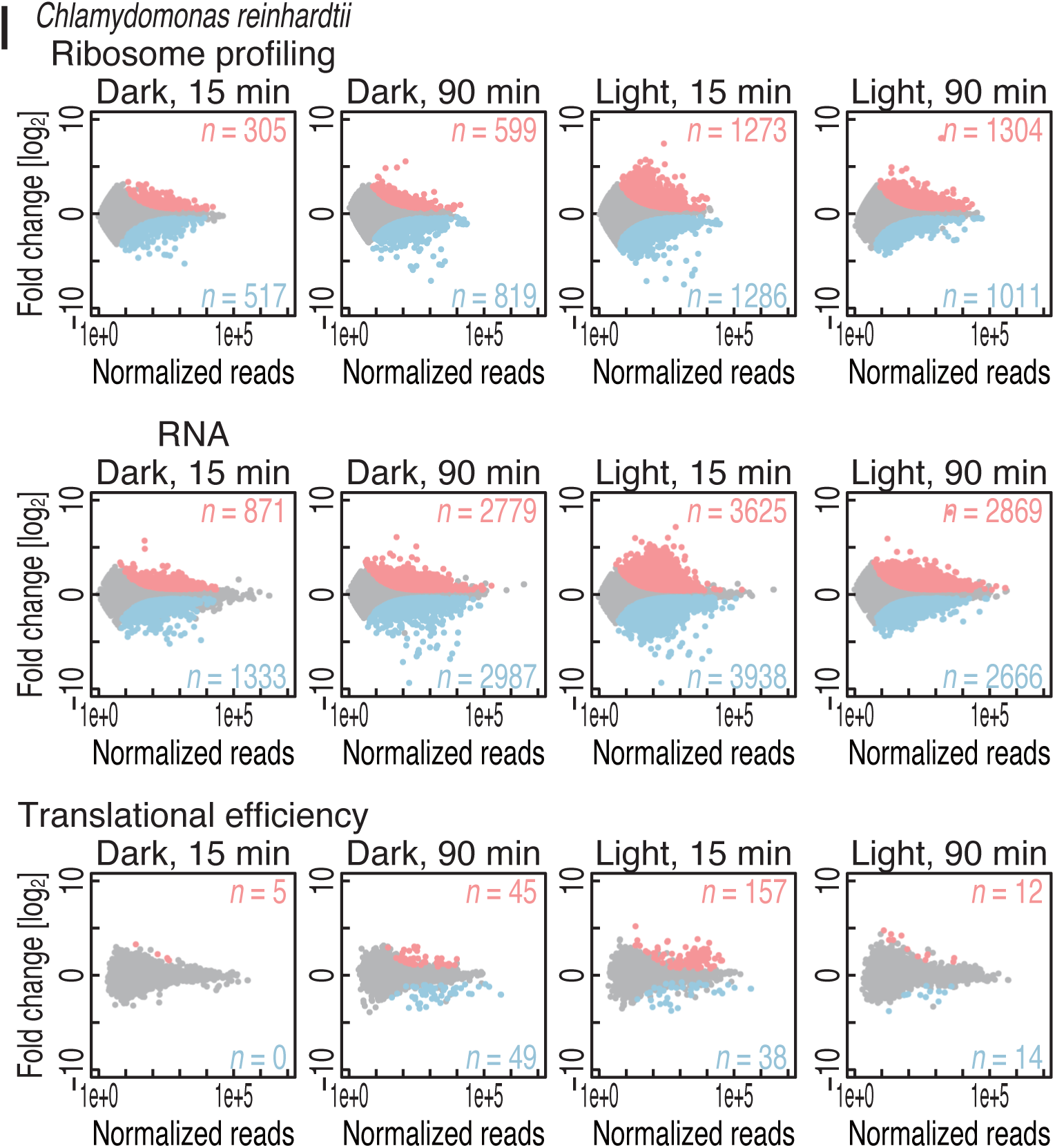
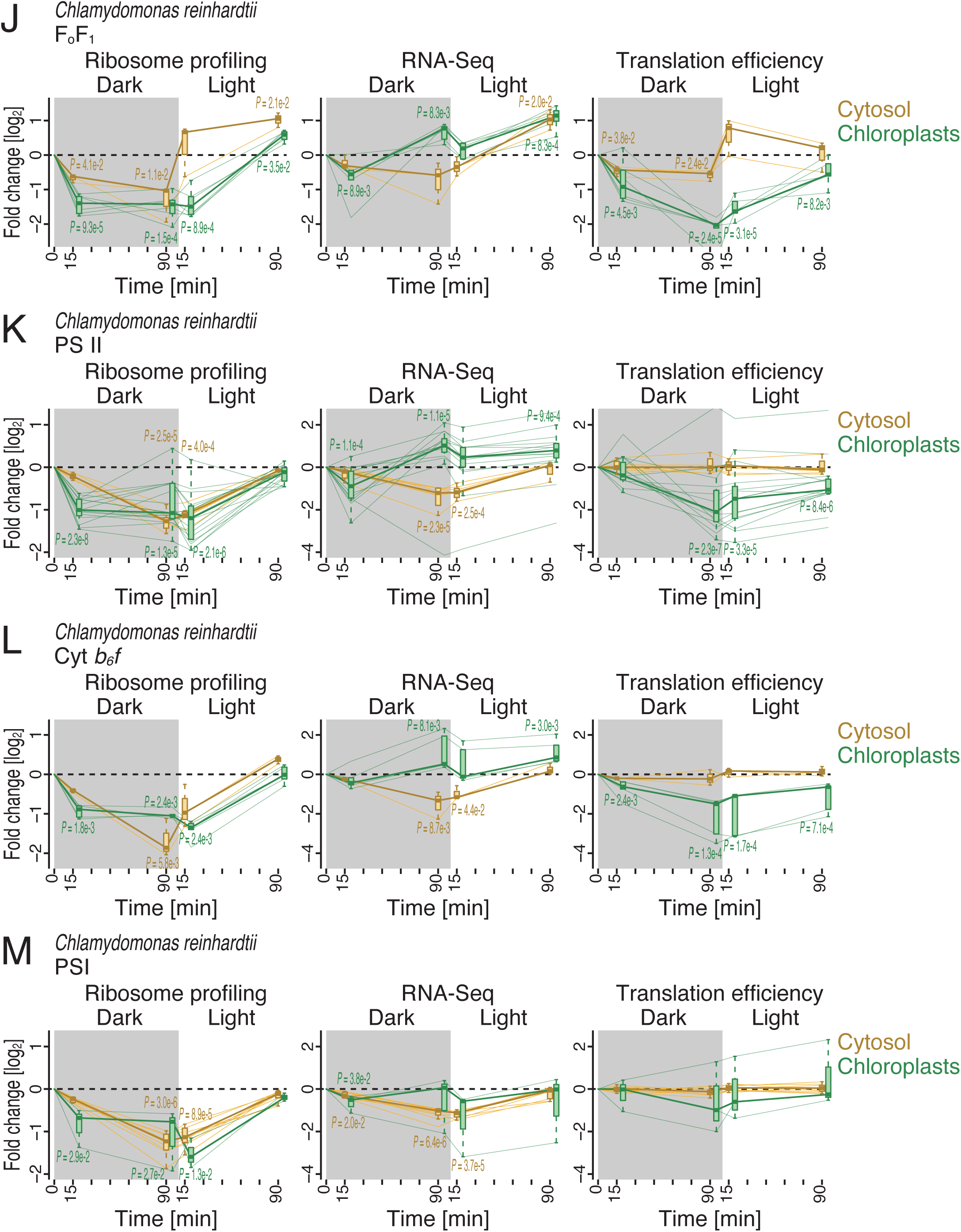
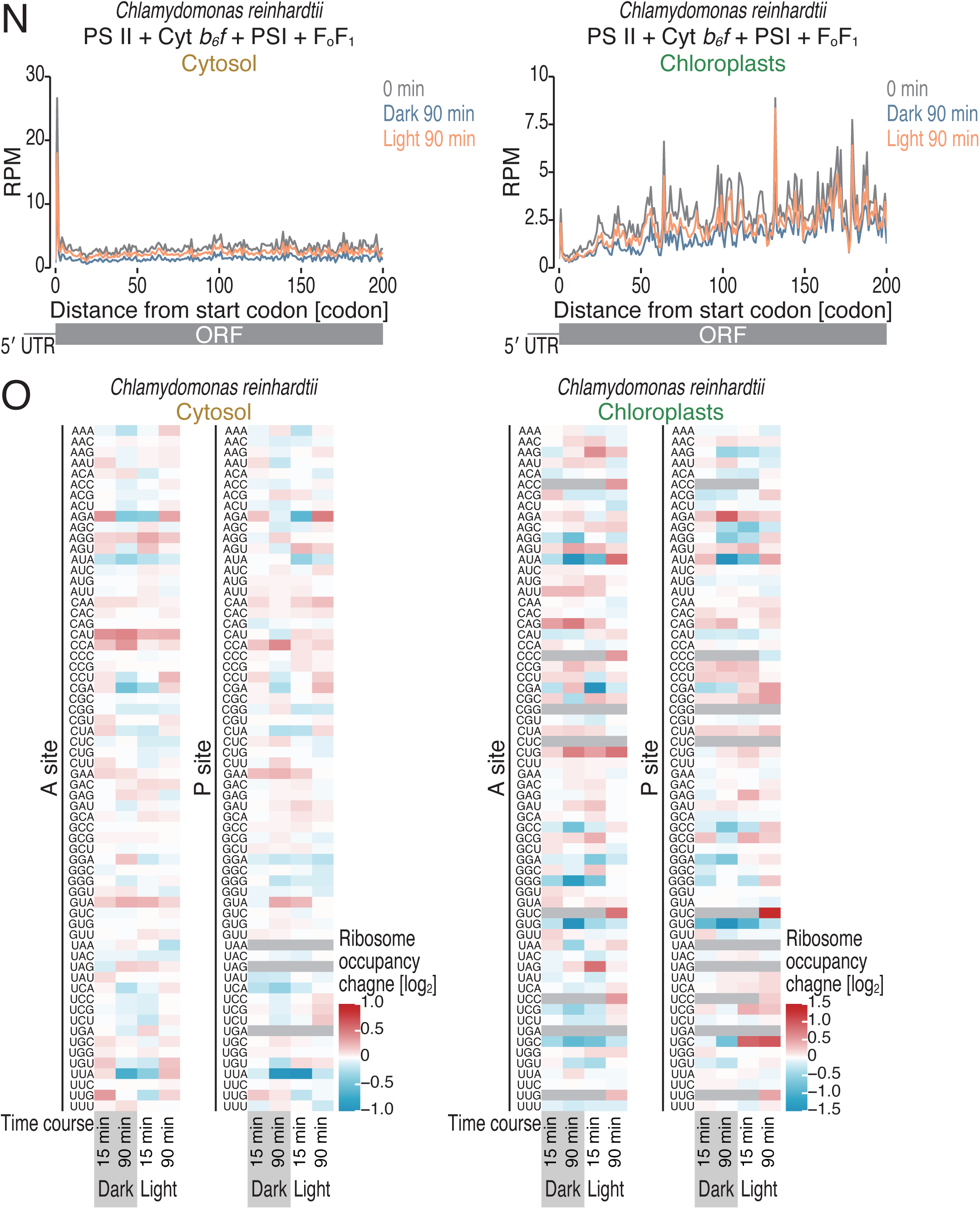
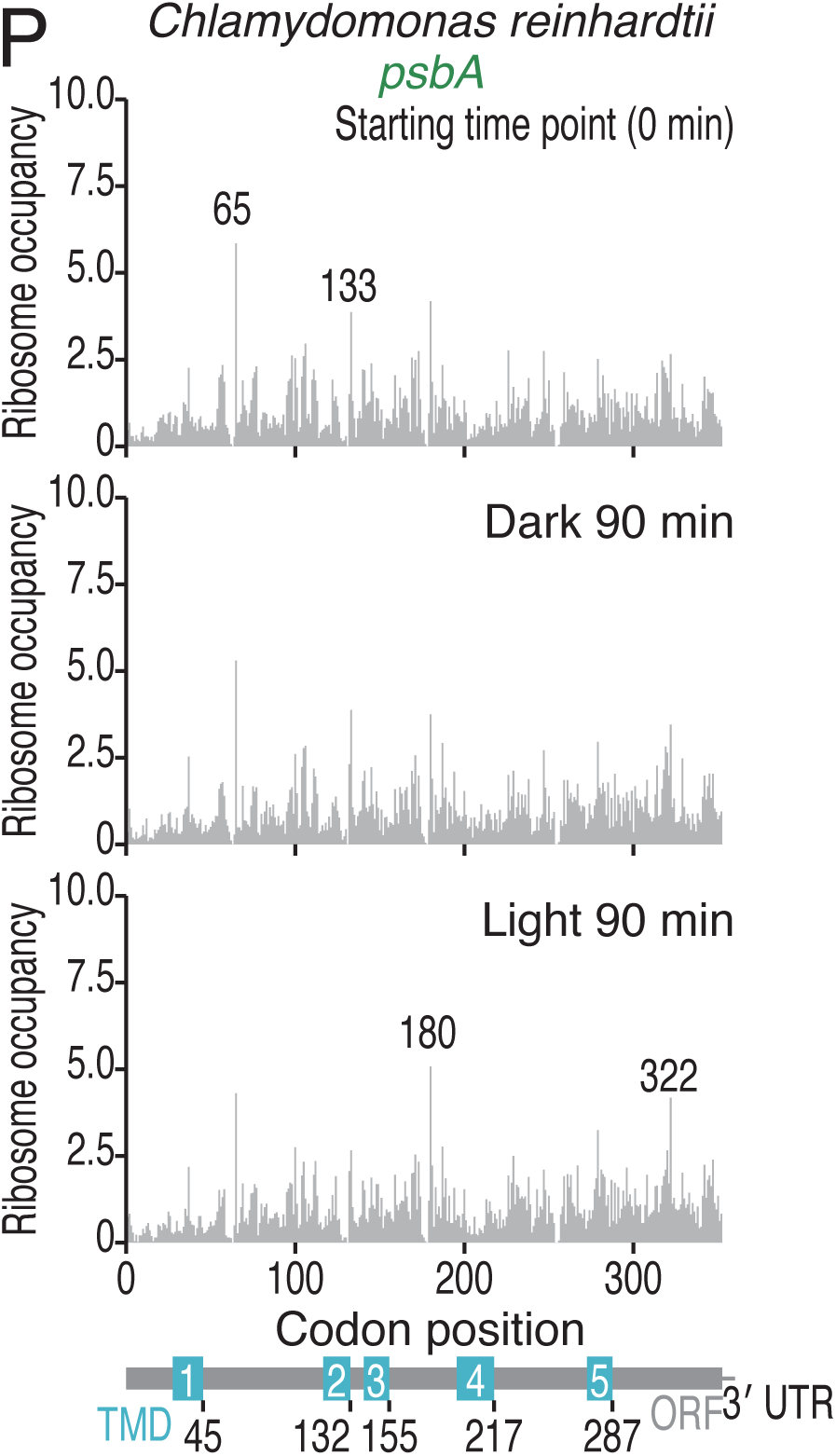
Characterization of ribosome profiling data in *C. reinhardtii*, related to Figure 3. (A) Pie chart showing the proportions of footprints from cytoribosomes and chlororibosomes in *C. reinhardtii*. (B) Length distributions of cytoribosome footprints and chlororibosome footprints in *C. reinhardtii* under the indicated conditions. (C) Tile plots of corresponding reading frames for the 5′ end of cytoribosome footprints (left) and chlororibosome footprints (right). The color scale indicates the read abundance. RPM, reads per million mapped reads. (D) Discrete Fourier transform of cytoribosome footprints (31 nt) (left) and chlororibosome footprints (27 nt) (right) downstream of the start codon (from position −9 to position 300). (E) Metagene plots of the 5′ ends of cytoribosome footprints (left) and chlororibosome footprints (right) along the length around start codons (the first nucleotide of the start codon was set to 0). The color scale indicates the read abundance. RPM, reads per million mapped reads. (F) Heatmap of the Pearson’s correlation coefficients (*r*) of the ORF-mapped reads in ribosome profiling and RNA-Seq under the indicated conditions. The value of *r* is indicated by the color scale. The black box tiles present the correlations of replicates. (G) Heatmap of the Pearson’s correlation coefficients (*r*) of the cytoribosome footprints and chlororibosome footprints on each codon under the indicated conditions. The value of *r* is indicated by the color scale. The black box tiles present the correlations of replicates. (H) Correlation between the normalized read density (reads/codon) and the subunit stoichiometry for the chloroplast FoF1 ATP synthase, with the zoomed-in inset highlighting data for one component factor. (I) MA (M, log ratio; A, mean average) plot for changes in ribosome profiling, RNA-Seq, and translation efficiency in the indicated conditions. Transcripts with significant changes (BH-adjusted *P* value < 0.05) are highlighted. (J-M) Changes in ribosome profiling, RNA-Seq, and translation efficiency during the light-to-dark and dark-to-light transitions for the subunits of FoF1 ATP synthase (J), PSII (K), Cyt *b6f* (L), and PSI (M) in *C. reinhardtii*. (N) Metagene plots for cryotribosome and chlororibosome footprints (the A site position) around start codons in the indicated conditions. mRNAs encoding the subunits of FoF1 ATP synthase, PSII, Cyt *b6f*, and PSI were analyzed. (O) Heatmaps for the cytoribosome and chlororibosome occupancy changes at A-site and P-site codons in the indicated conditions. The color scale indicates ribosome occupancy change. (P) Distributions of chlororibosome footprints along the *psbA* mRNA under the indicated conditions. The A-site position of each read (the start codon was set to 0) is indicated. The box plots (J-M) show the median (center line), upper/lower quartiles (box limits), and 1.5× interquartile range (whiskers). The black line indicates the median at each time point.

**Figure S5.**
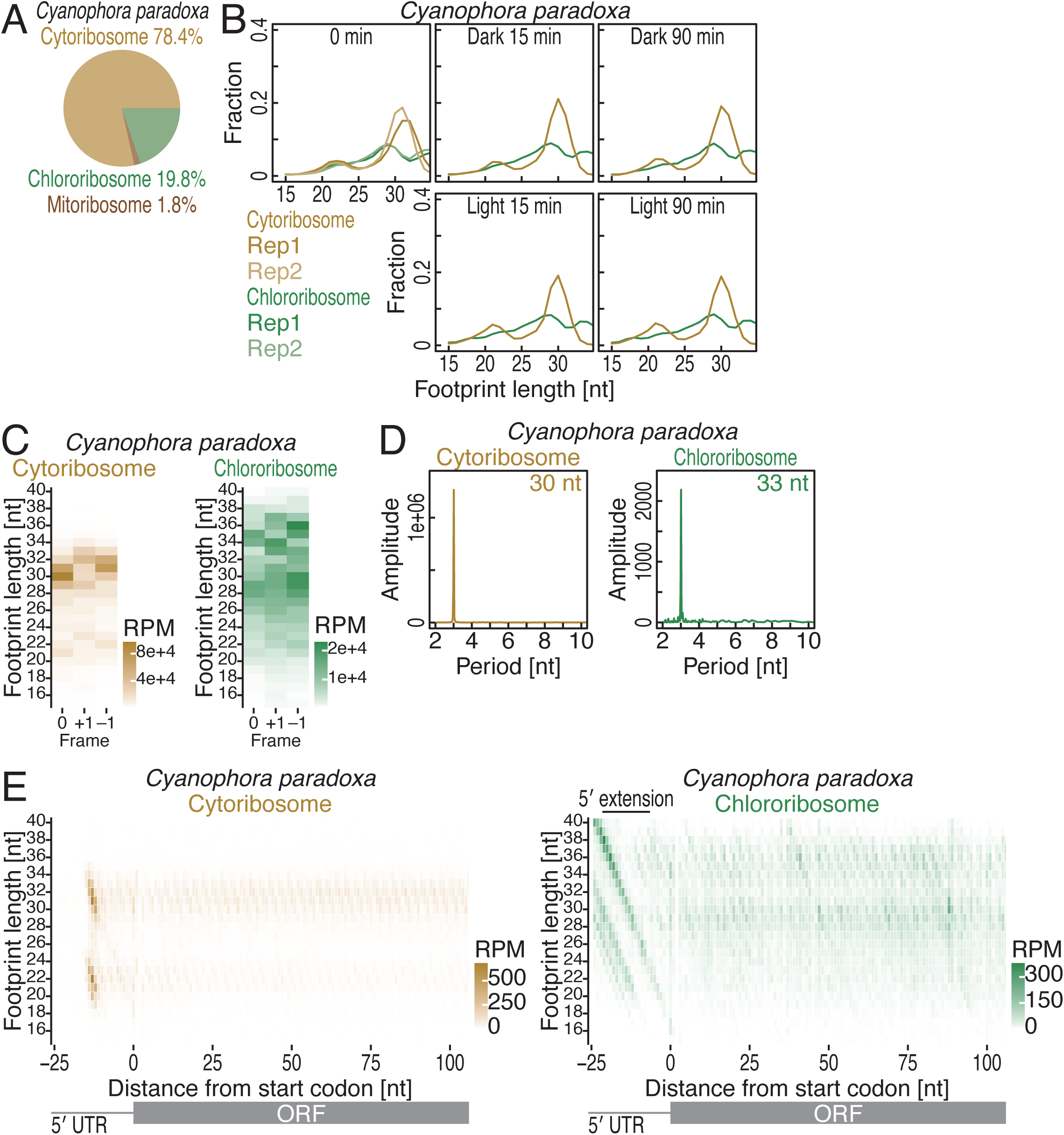
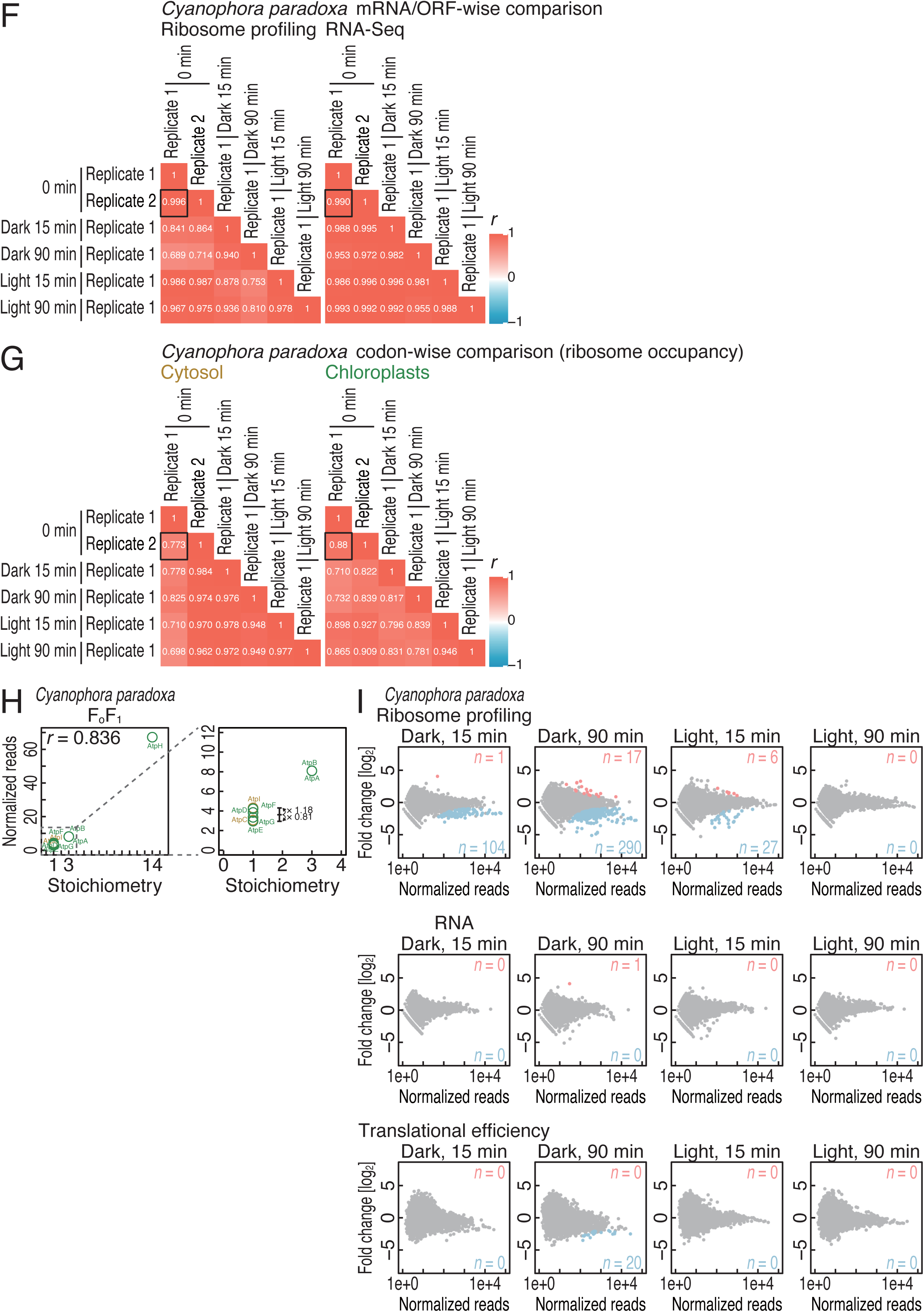
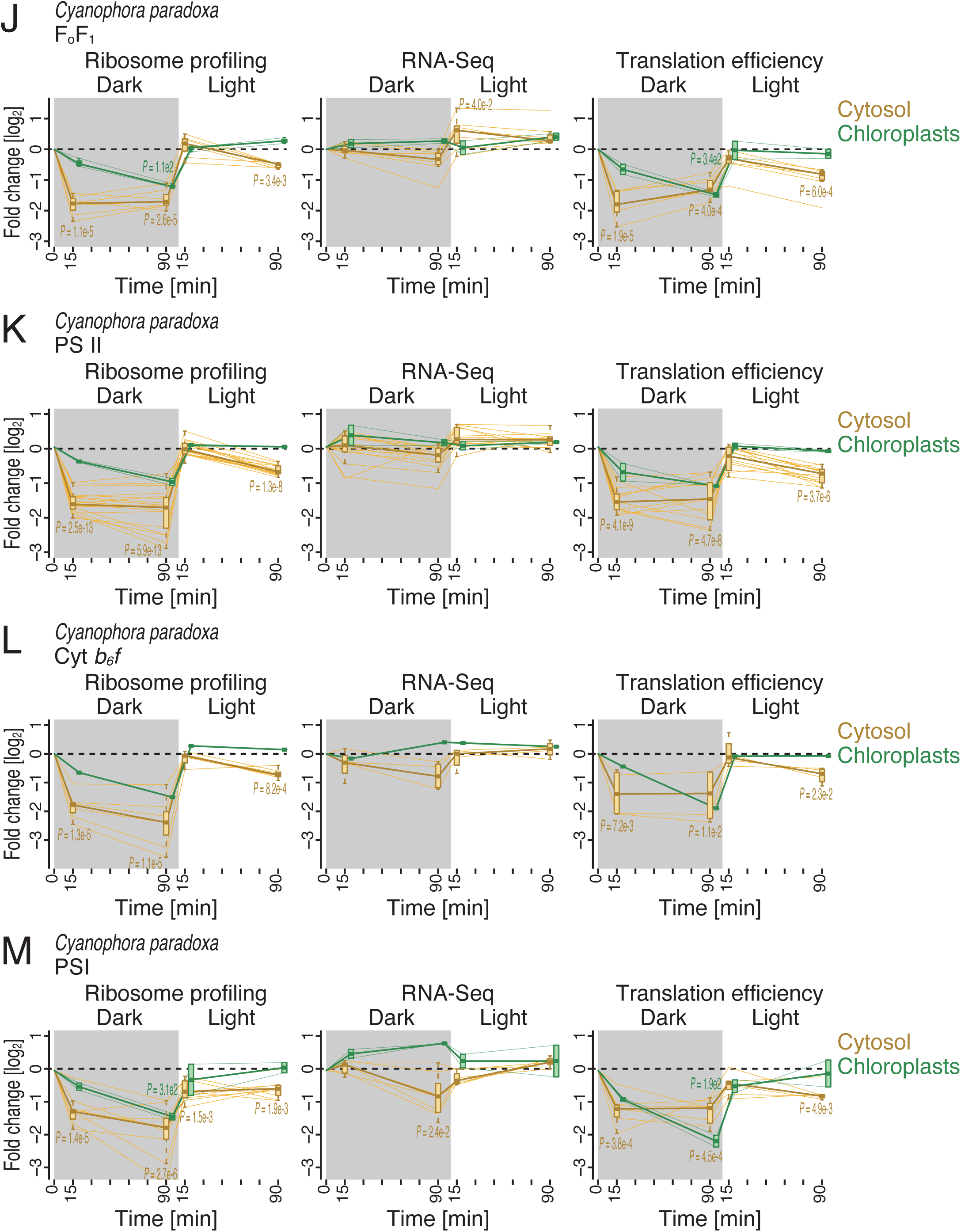
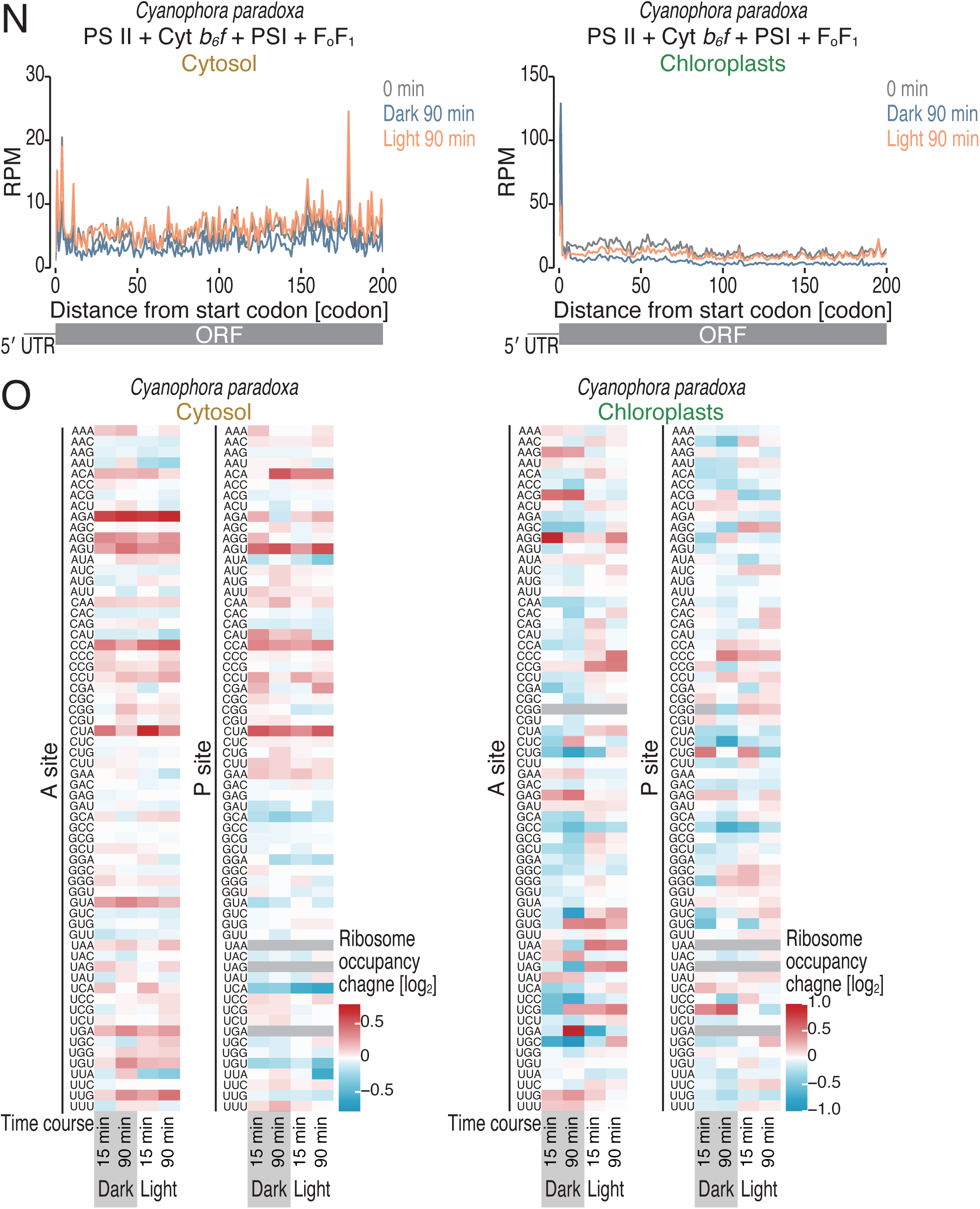
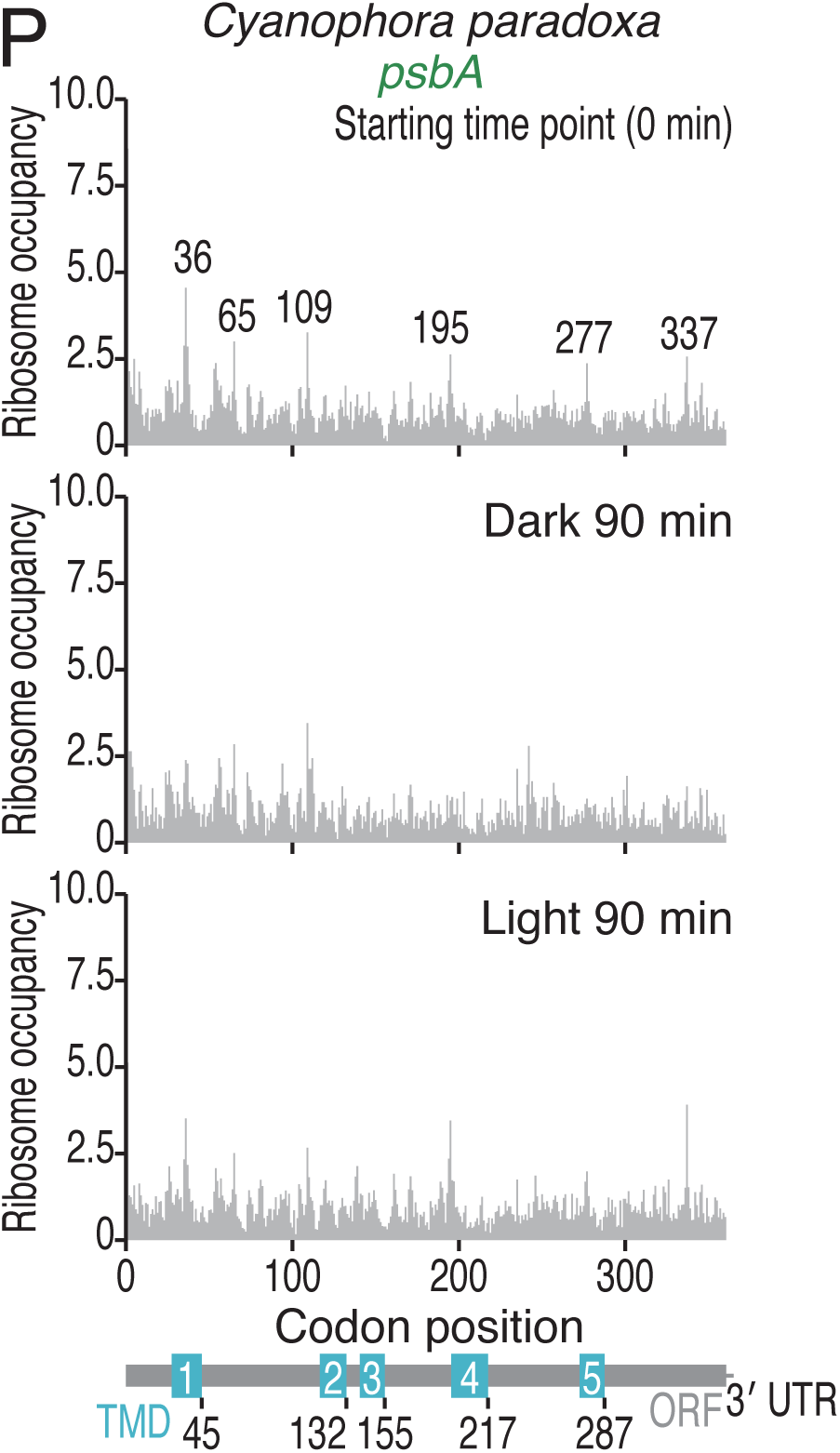
Characterization of ribosome profiling data for *C. paradoxa*, related to Figure 3. (A) Pie chart showing the proportions of footprints from cytoribosomes and chlororibosomes in *C. paradoxa*. (B) Length distributions of cytoribosome footprints and chlororibosome footprints in *C. paradoxa* under the indicated conditions. (C) Tile plots of corresponding reading frames for the 5′ end of cytoribosome footprints (left) and chlororibosome footprints (right). The color scale indicates the read abundance. RPM, reads per million mapped reads. (D) Discrete Fourier transform of cytoribosome footprints (30 nt) (left) and chlororibosome footprints (33 nt) (right) downstream of the start codon (from position −9 to position 300). (E) Metagene plots of the 5′ ends of cytoribosome footprints (left) and chlororibosome footprints (right) along the length around start codons (the first nucleotide of the start codon was set to 0). The color scale indicates the read abundance. RPM, reads per million mapped reads. (F) Heatmap of the Pearson’s correlation coefficients (*r*) of the ORF-mapped reads in ribosome profiling and RNA-Seq under the indicated conditions. The value of *r* is indicated by the color scale. The black box tiles present the correlations of replicates. (G) Heatmap of the Pearson’s correlation coefficients (*r*) of the cytoribosome footprints and chlororibosome footprints on each codon under the indicated conditions. The value of *r* is indicated by the color scale. The black box tiles present the correlations of replicates. (H) Correlation between the normalized read density (reads/codon) and the subunit stoichiometry for the chloroplast FoF1 ATP synthase, with the zoomed-in inset highlighting data for one component factor. (I) MA (M, log ratio; A, mean average) plot for changes in ribosome profiling, RNA-Seq, and translation efficiency in the indicated conditions. Transcripts with significant changes (BH-adjusted *P* value < 0.05) are highlighted. (J-M) Changes in ribosome profiling, RNA-Seq, and translation efficiency during the light-to-dark and dark-to-light transitions for the subunits of FoF1 ATP synthase (J), PSII (K), Cyt *b6f* (L), and PSI (M) in *C. paradoxa*. (N) Metagene plots for cryotribosome and chlororibosome footprints (the A site position) around start codons in the indicated conditions. mRNAs encoding the subunits of FoF1 ATP synthase, PSII, Cyt *b6f*, and PSI were analyzed. (O) Heatmaps for the cytoribosome and chlororibosome occupancy changes at A-site and P-site codons in the indicated conditions. The color scale indicates ribosome occupancy change. (P) Distributions of chlororibosome footprints along the *psbA* mRNA under the indicated conditions. The A-site position of each read (the start codon was set to 0) is indicated. The box plots (J-M) show the median (center line), upper/lower quartiles (box limits), and 1.5× interquartile range (whiskers). The black line indicates the median at each time point.

**Figure S6.**
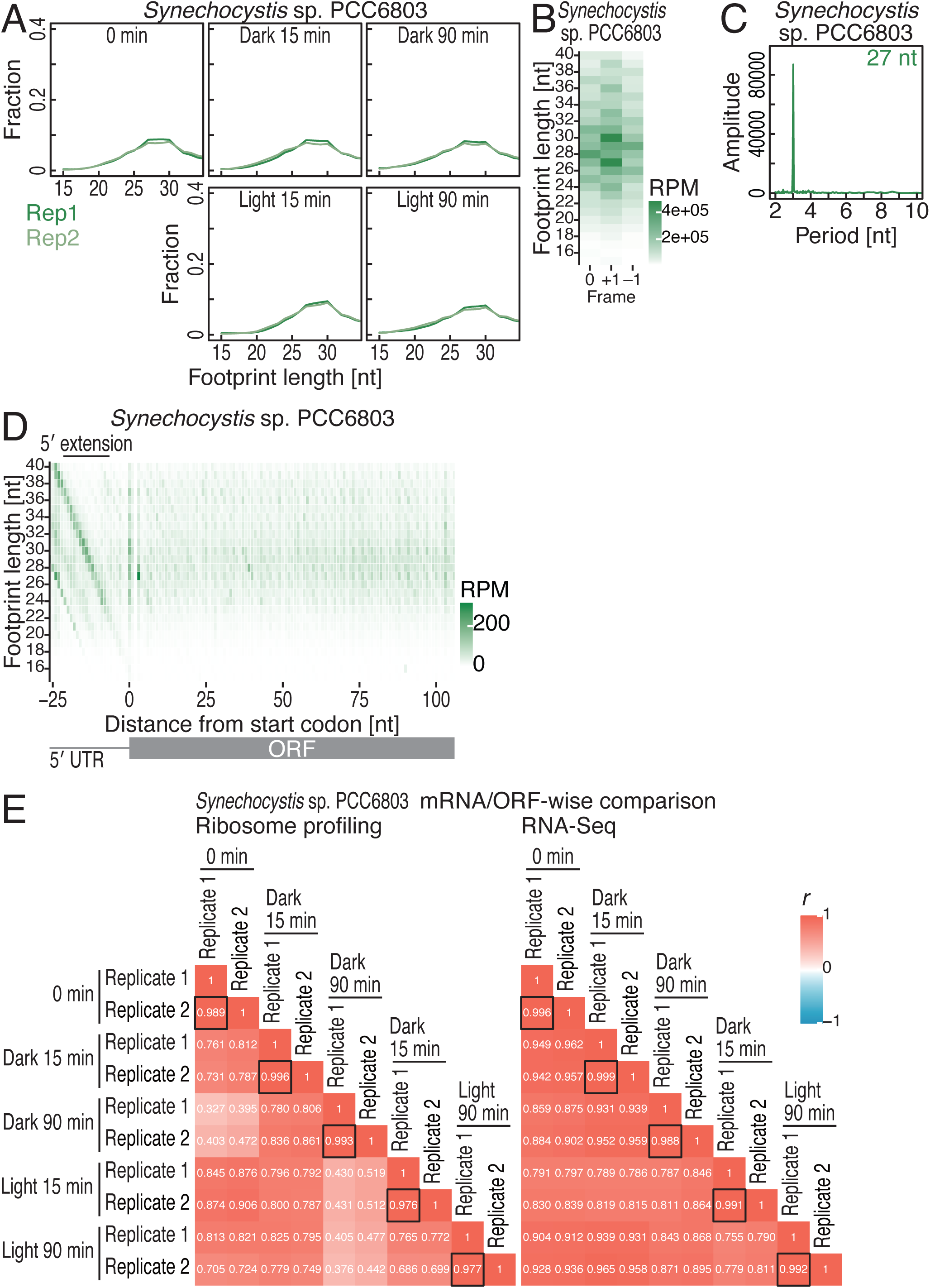
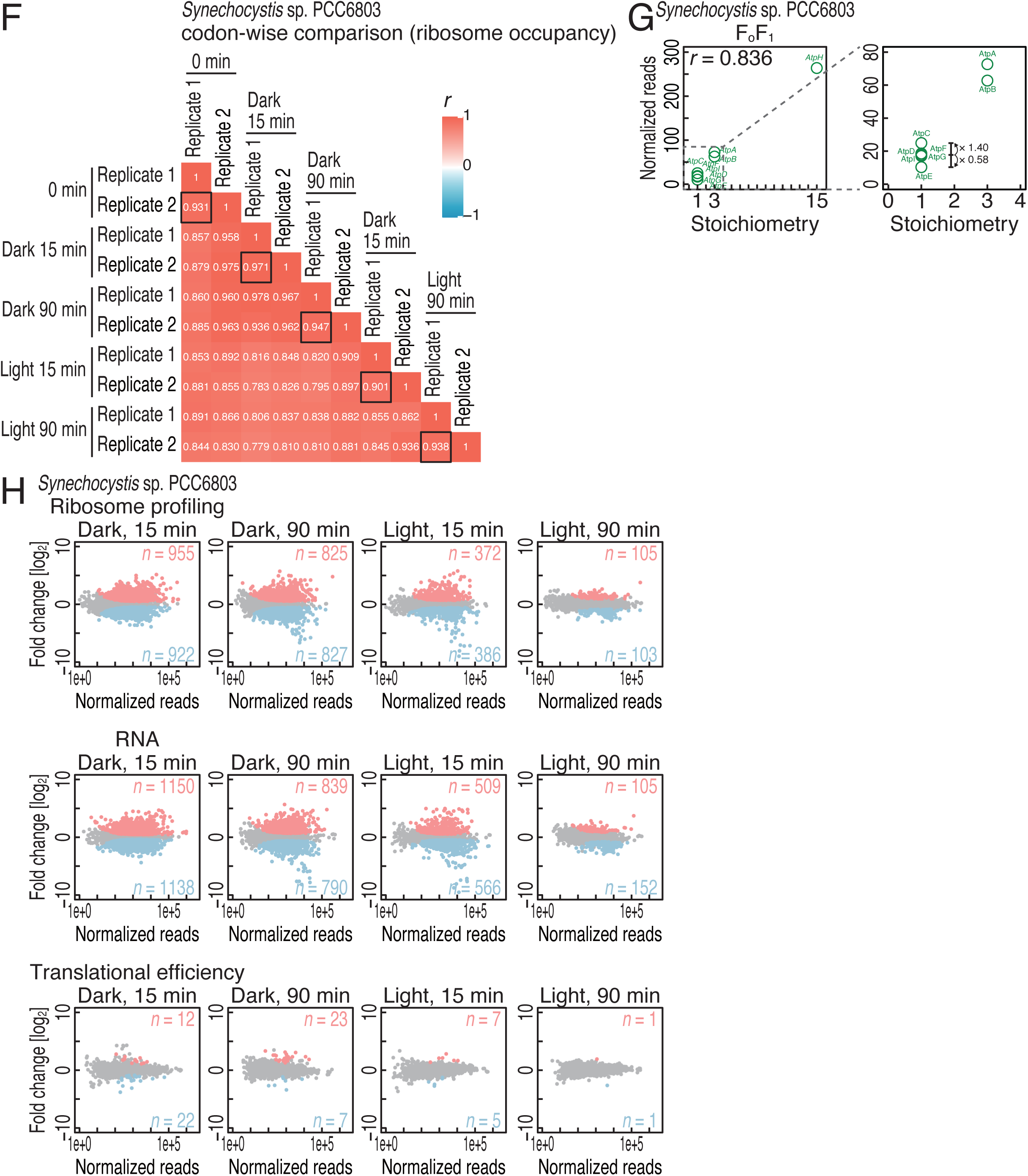
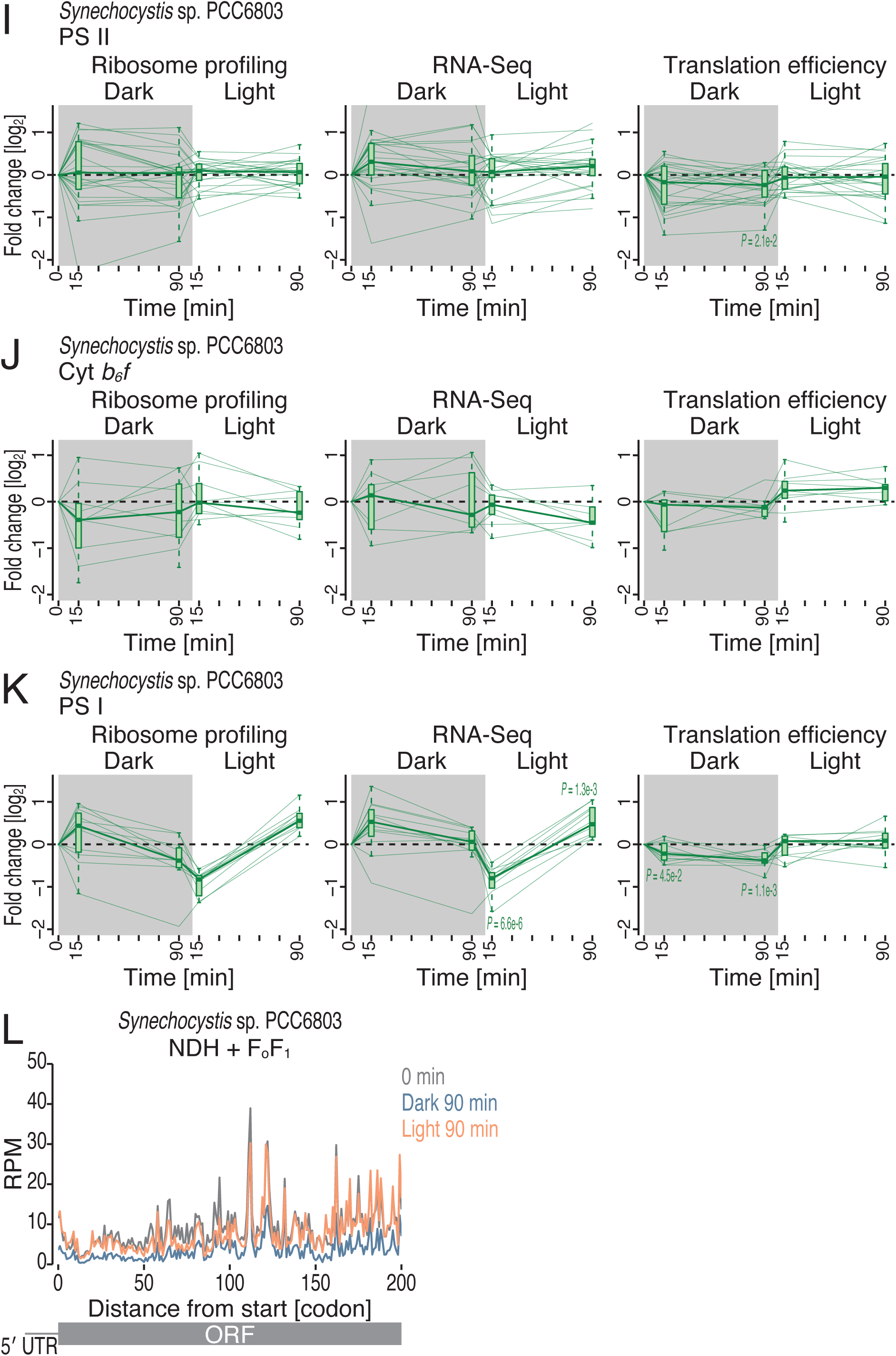
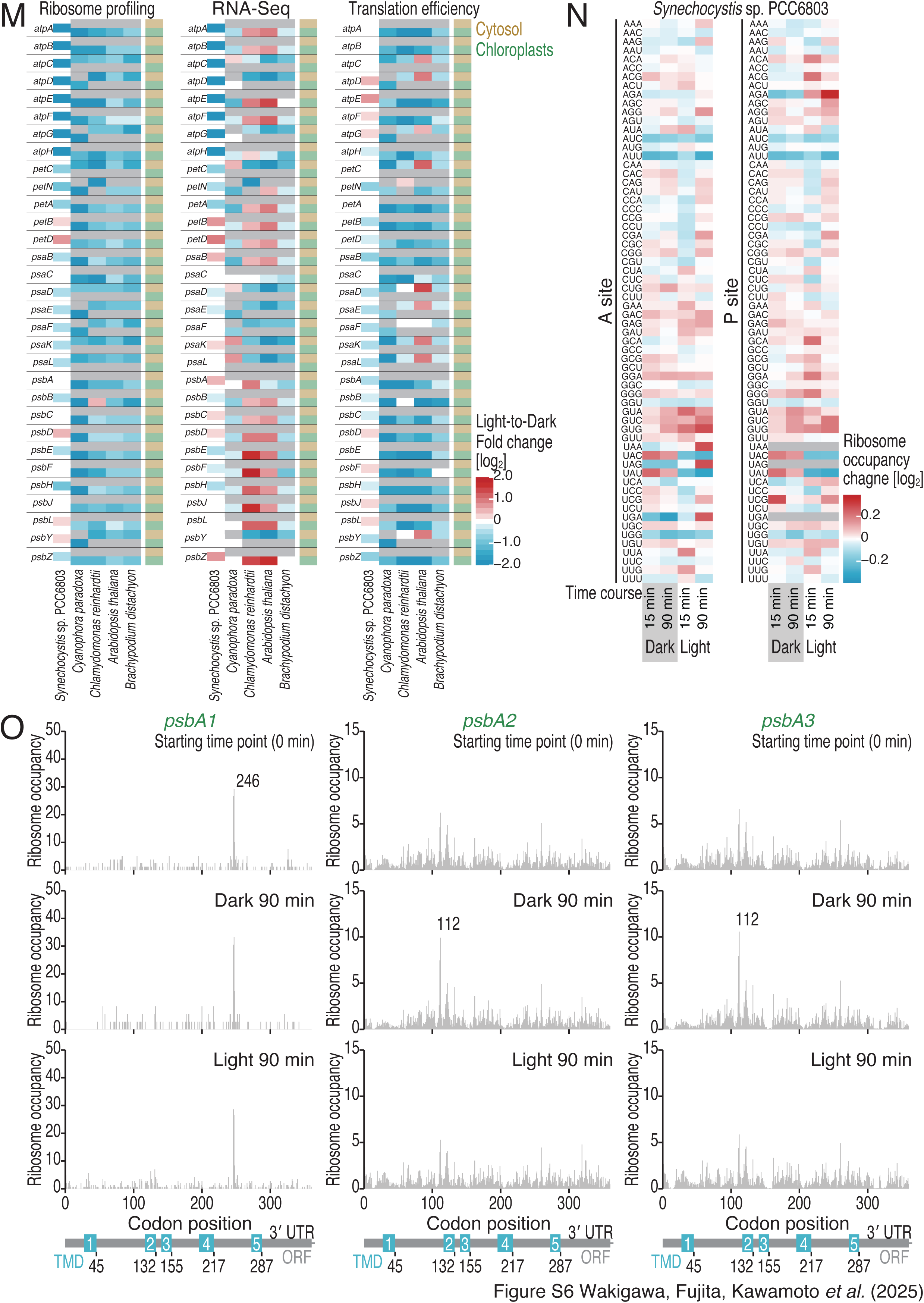
Characterization of ribosome profiling data in *Synechocystis* sp. PCC6803, related to Figure 5. (A) Length distributions of ribosome footprints and chlororibosome footprints in *Synechocystis* sp. PCC6803 under the indicated conditions. (B) Tile plot of corresponding reading frames for the 5′ end of ribosome footprints. The color scale indicates the read abundance. RPM, reads per million mapped reads. (C) Discrete Fourier transform of ribosome footprints (27 nt) downstream of the start codon (from position −9 to position 300). (D) Metagene plots of the 5′ ends of ribosome footprints along the length around start codons (the first nucleotide of the start codon was set to 0). The color scale indicates the read abundance. RPM, reads per million mapped reads. (E) Heatmap of the Pearson’s correlation coefficients (*r*) of the ORF-mapped reads in ribosome profiling and RNA-Seq under the indicated conditions. The value of *r* is indicated by the color scale. The black box tiles present the correlations of replicates. (F) Heatmap of the Pearson’s correlation coefficients (*r*) of the footprints on each codon under the indicated conditions. The value of *r* is indicated by the color scale. The black box tiles present the correlations of replicates. (G) Correlation between the normalized read density (reads/codon) and the subunit stoichiometry for the chloroplast FoF1 ATP synthase, with the zoomed-in inset highlighting data for one component factor. (H) MA (M, log ratio; A, mean average) plot for changes in ribosome profiling, RNA-Seq, and translation efficiency in the indicated conditions. Transcripts with significant changes (BH-adjusted *P* value < 0.05) are highlighted. (I-K) Changes in ribosome profiling, RNA-Seq, and translation efficiency during the light-to-dark and dark-to-light transitions for the subunits of PSII (I), Cyt *b6f* (J), and PSI (K) in *Synechocystis* sp. PCC6803. (L) Metagene plots for ribosome footprints (the A site position) around start codons in the indicated conditions. mRNAs encoding the subunits of FoF1 ATP synthase and NDH were analyzed. (M) Heatmaps for changes in ribosome profiling, RNA-Seq, and translation efficiency for the conserved genes [best-best pairs in Kyoto Encyclopedia of Genes and Genomes (KEGG)] in the indicated species. Analyses were restricted to the genes that showed reads in all the conditions, in all species, and both ribosome profiling and RNA-Seq. The data of the late dark time point (180 min for *A. thaliana* and *B. distachyon*; 90 min for *C. reinhardtii*, *C. paradoxa*, and *Synechocystis* sp. PCC6803) are shown. Color code (yellow-orange and green) represents genomes where the genes are encoded. The color scale indicates their changes. (N) Heatmaps for the ribosome occupancy changes at A-site and P-site codons in the indicated conditions. The color scale indicates ribosome occupancy change. (O) Distributions of chlororibosome footprints along the *psbA1*, *psbA2*, and *psbA3* mRNAs under the indicated conditions. The A-site position of each read (the start codon was set to 0) is indicated. The box plots (I-K) show the median (center line), upper/lower quartiles (box limits), and 1.5× interquartile range (whiskers). The black line indicates the median at each time point.

**Figure S7.**
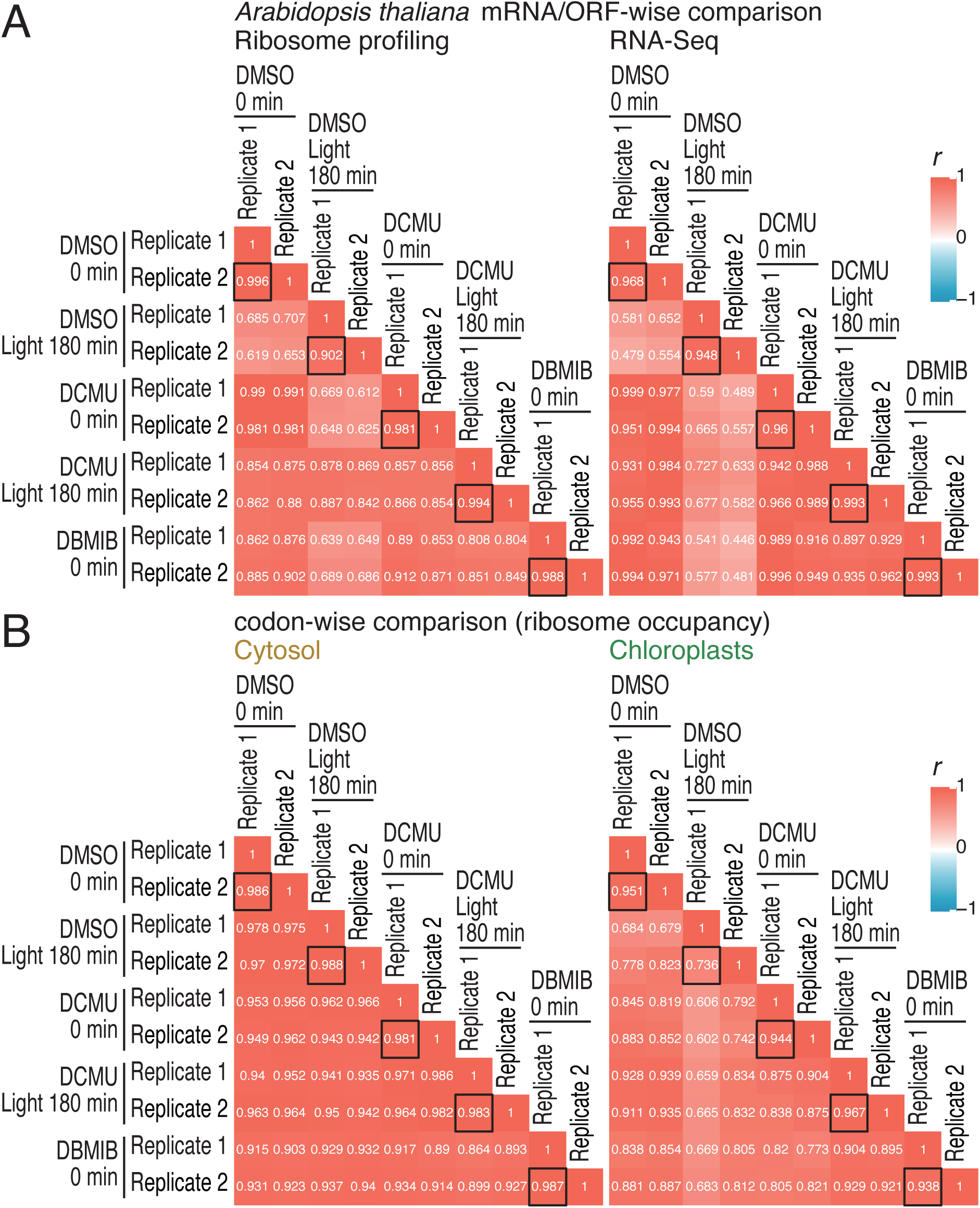
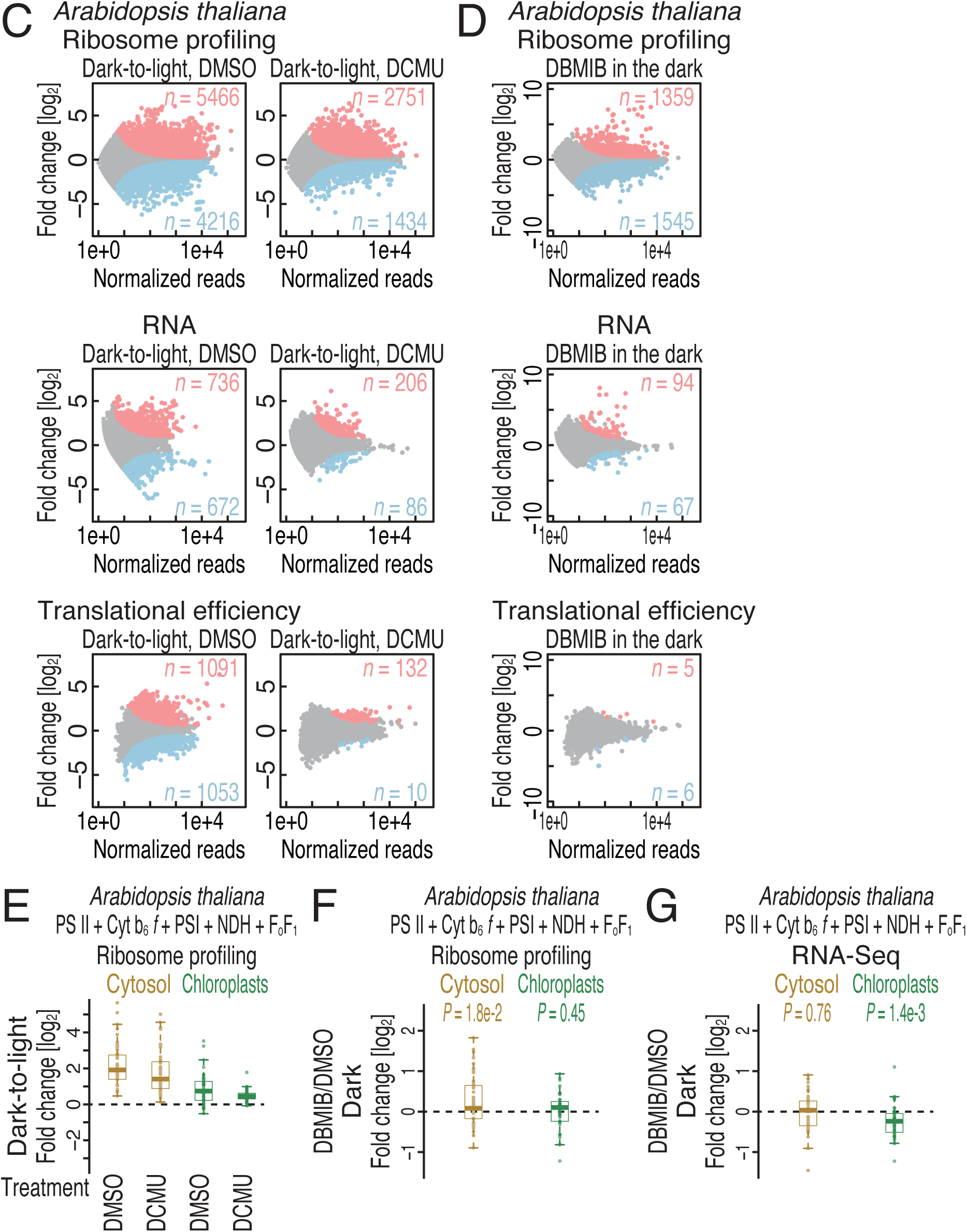
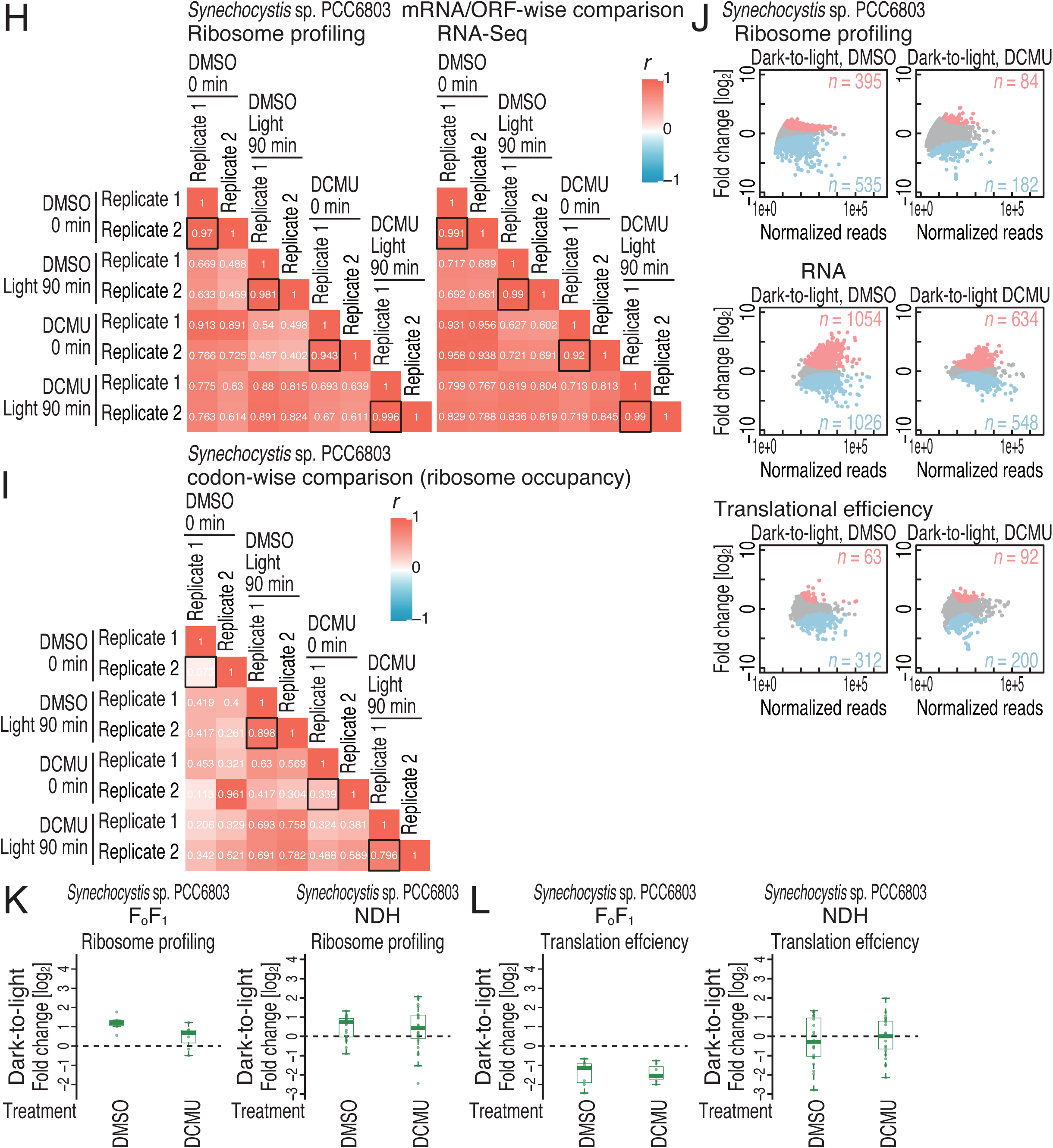
Characterization of DCMU and DBMIB treatment, related to Figure 6. (A and H) Heatmap of the Pearson’s correlation coefficients (*r*) of the ORF-mapped reads in ribosome profiling and RNA-Seq under the indicated conditions. The value of *r* is indicated by the color scale. The black box tiles present the correlations of replicates. (B) Heatmap of the Pearson’s correlation coefficients (*r*) of the cytoribosome footprints and chlororibosome footprints on each codon under the indicated conditions. The value of *r* is indicated by the color scale. The black box tiles present the correlations of replicates. (C-D and J) MA (M, log ratio; A, mean average) plot for changes in ribosome profiling, RNA-Seq, and translation efficiency in the indicated conditions. Transcripts with significant changes (BH-adjusted *P* value < 0.05) are highlighted. (E) Fold change in ribosome footprints following DCMU treatment of *A. thaliana* during the dark-to-light transition. Cytosolic mRNAs and chloroplast mRNAs encoding the subunits of FoF1 ATP synthase, PSII, Cyt *b6f*, PSI, and NDH were analyzed. (F) Fold change in ribosome footprints following DBMIB treatment of *A. thaliana* in the dark. Cytosolic mRNAs and chloroplast mRNAs encoding the subunits of FoF1 ATP synthase, PSII, Cyt *b6f*, PSI, and NDH were analyzed. (G) Fold change in RNA abundances following DBMIB treatment of *A. thaliana* in the dark. Cytosolic mRNAs and chloroplast mRNAs encoding the subunits of FoF1 ATP synthase, PSII, Cyt *b6f*, PSI, and NDH were analyzed. (I) Heatmap of the Pearson’s correlation coefficients (*r*) of the ribosome footprints on each codon under the indicated conditions. The value of *r* is indicated by the color scale. The black box tiles present the correlations of replicates. (K) Fold changes in ribosome footprints following DCMU treatment of *Synechocystis* sp. PCC6803 during the dark-to-light transition. mRNAs encoding the subunits of FoF1 ATP synthase (left) and NDH (right) were analyzed. (L) Fold change in the translation efficiency following DCMU treatment of *Synechocystis* sp. PCC6803 during the dark-to-light transition. mRNAs encoding the subunits of FoF1 ATP synthase (left) and NDH (right) were analyzed. The box plots (E-G and K-L) show the median (center line), upper/lower quartiles (box limits), and 1.5× interquartile range (whiskers). Significance was determined by the Mann‒Whitney *U* test.

**Figure S8.**
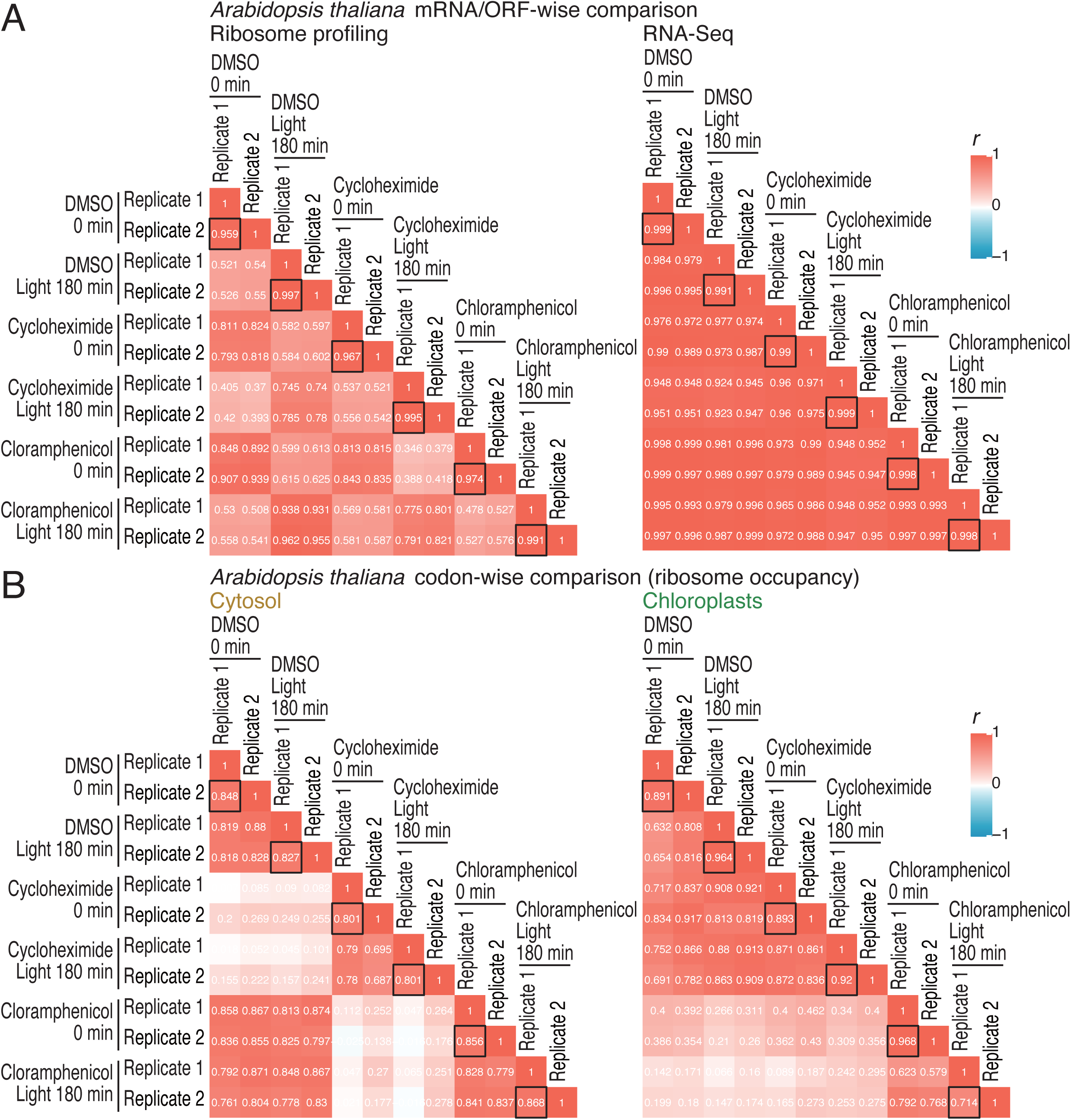
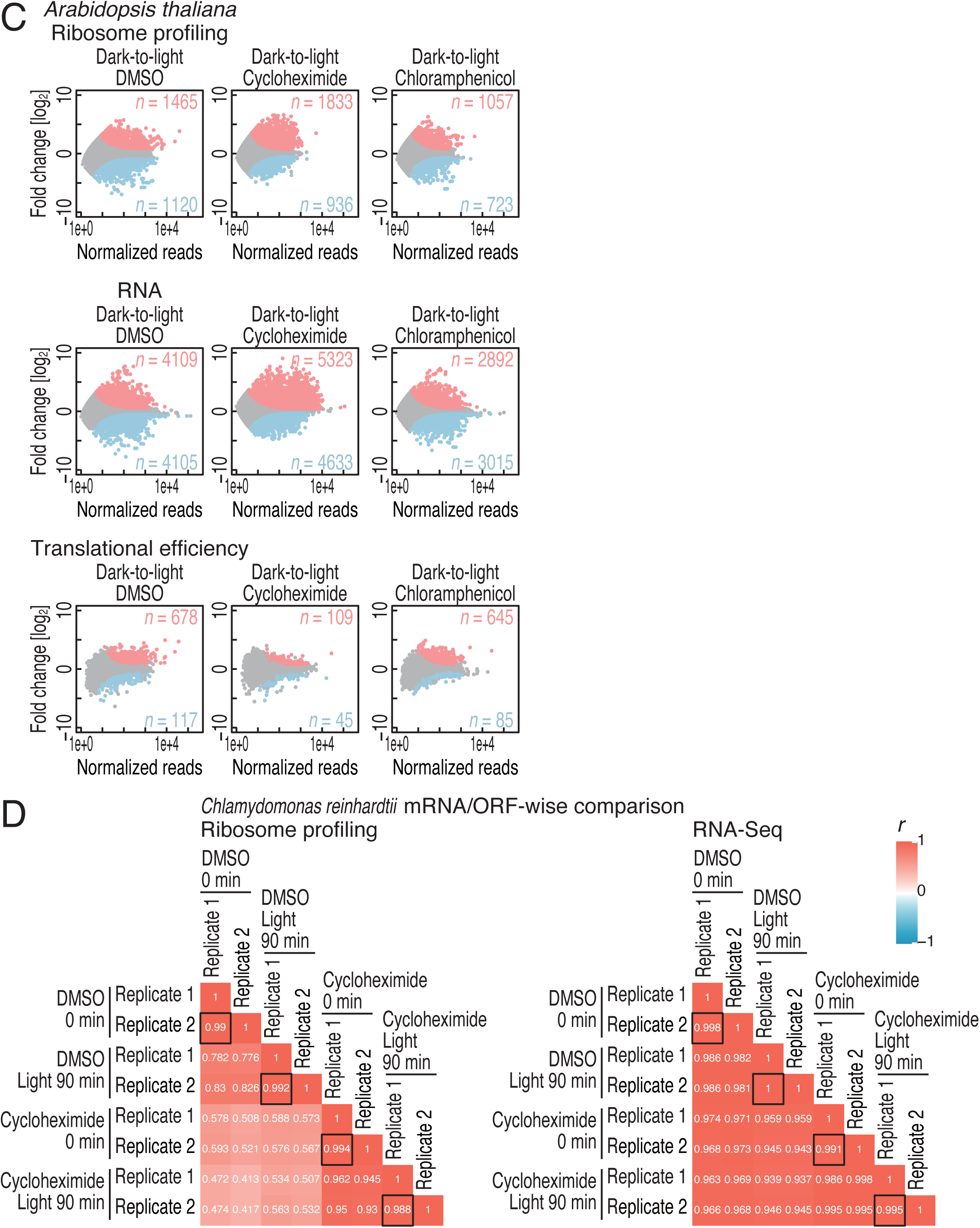
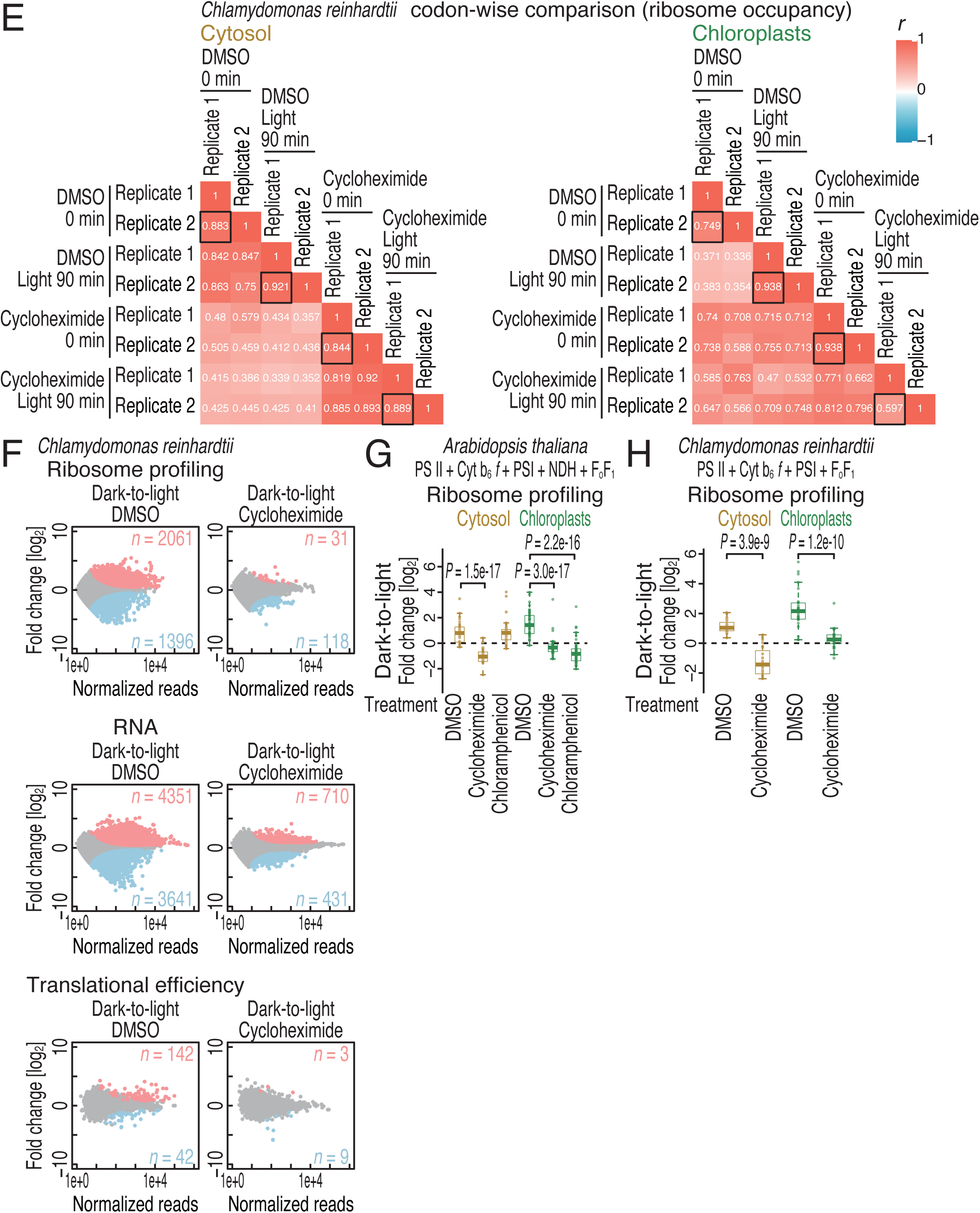
Characterization of translation inhibitor treatment, related to Figure 7. (A and D) Heatmap of the Pearson’s correlation coefficients (*r*) of the ORF-mapped reads in ribosome profiling and RNA-Seq under the indicated conditions. The value of *r* is indicated by the color scale. The black box tiles present the correlations of replicates. (B and E) Heatmap of the Pearson’s correlation coefficients (*r*) of the cytoribosome footprints and chlororibosome footprints on each codon under the indicated conditions. The value of *r* is indicated by the color scale. The black box tiles present the correlations of replicates. (C and F) MA (M, log ratio; A, mean average) plot for changes in ribosome profiling, RNA-Seq, and translation efficiency in the indicated conditions. Transcripts with significant changes (BH-adjusted *P* value < 0.05) are highlighted. (G) Fold change in ribosome footprints following translation inhibitor treatment of *A. thaliana* during the dark-to-light transition. Cytosolic mRNAs and chloroplast mRNAs encoding the subunits of FoF1 ATP synthase, PSII, Cyt *b6f*, PSI, and NDH were analyzed. (H) Fold change in ribosome footprints following translation inhibitor treatment of *C. reinhardtii* during the dark-to-light transition. Cytosolic mRNAs and chloroplast mRNAs encoding the subunits of FoF1 ATP synthase, PSII, Cyt *b6f*, and PSI were analyzed. The box plots (G and H) show the median (center line), upper/lower quartiles (box limits), and 1.5× interquartile range (whiskers). Significance was determined by the Mann‒ Whitney *U* test.

Table S1. List of genes encoding subunits of photosynthesis complexes, related to Figures 1-6.

Complex name, gene name, and transcript ID for species studies in this study are listed.

Table S2. List of ribosome profiling and RNA-Seq libraries, related to Figures 1-6. For each data set, the library name, species, strain name, light/dark condition, drug treatment, replicate number, library type, RNase condition, the read length used for cytoribosome footprint, A-site offset for cytoribosome footprint, the read length used for chlororibosome footprint, A-site offsets for chlororibosome footprint, the read lengths used for mitoribosome footprint, A-site offset for mitoribosome footprint, accession number, and reference are listed.

Table S3. Changes in ribosome footprints, RNAs, and translation efficiency along light-dark transitions in A. thaliana, related to Figures 1-2.

mRNA ID, log2 fold change at each condition compared to the condition before the transition, and BH-adjusted P value are listed.

Table S4. Changes in ribosome footprints, RNAs, and translation efficiency along light-dark transitions in B. distachyon, related to Figure 3.

mRNA ID, log2 fold change at each condition compared to the condition before the transition, and BH-adjusted P value are listed.

Table S5. Changes in ribosome footprints, RNAs, and translation efficiency along light-dark transitions in C. reinhardtii, related to Figure 3.

mRNA ID, log2 fold change at each condition compared to the condition before the transition, and BH-adjusted P value are listed.

Table S6. Changes in ribosome footprints, RNAs, and translation efficiency along light-dark transitions in C. paradoxa, related to Figure 3.

mRNA ID, log2 fold change at each condition compared to the condition before the transition, and BH-adjusted P value are listed.

Table S7. Changes in ribosome footprints, RNAs, and translation efficiency along light-dark transitions in Synechocystis sp. PCC6803, related to Figure 5.

mRNA ID, log2 fold change at each condition compared to the condition before the transition, and BH-adjusted P value are listed.

Table S8. Changes in ribosome footprints, RNAs, and translation efficiency with DCMU treatment in A. thaliana, related to Figure 6.

mRNA ID, log2 fold change during the dark-to-light transition with DMSO or DCMU treatment, and BH-adjusted P value are listed.

Table S9. Changes in ribosome footprints, RNAs, and translation efficiency with DBMIB treatment in A. thaliana, related to Figure 6.

mRNA ID, log2 fold change by DMBIB treatment in the dark, and BH-adjusted P value are listed.

Table S10. Changes in ribosome footprints, RNAs, and translation efficiency with DCMU treatment in Synechocystis sp. PCC6803, related to Figure 6.

mRNA ID, log2 fold change during the dark-to-light transition with DMSO or DCMU, and BH-adjusted P value are listed.

Table S11. Changes in ribosome footprints, RNAs, and translation efficiency with translation inhibitor treatment in A. thaliana, related to Figure 7.

mRNA ID, log2 fold change during the dark-to-light transition with DMSO, cycloheximide, or chloramphenicol, and BH-adjusted P value are listed.

Table S12. Changes in ribosome footprints, RNAs, and translation efficiency with translation inhibitor treatment in C. reinhardtii, related to Figure 7.

mRNA ID, log2 fold change during the dark-to-light transition with DMSO or cycloheximide, and BH-adjusted P value are listed.

